# Molecular Mechanism of Lipid Recognition and Membrane-Guided Gating in Plant Minimal START Proteins

**DOI:** 10.64898/2026.07.22.740216

**Authors:** Kamlesh Kumari, Sanjeet Kumar Mahtha, Megha Parihar, Garima Tiwari, Gitanjali Yadav, Vineet Gaur

**Affiliations:** BRIC-National Institute of Plant Genome Research, Aruna Asaf Ali Marg, New Delhi, India 110067; Pinnacle Medicines US Inc.1980 S. Easton Road, Doylestown, PA 18901

**Author notes:** **Correspondence:** Dr. Vineet Gaur, Scientist, BRIC-National Institute of Plant Genome Research, Aruna Asaf Ali Marg, New Delhi, India 110067.

**Keywords:** Plant minimal START proteins, StARkin superfamily, non-vesicular lipid transport, α/β helix-grip fold, Hydrophobic cavity

## Abstract

The hydrophobic nature of lipids requires specialized transport mechanisms, and one such non-vesicular transport mechanism involves StAR-related lipid transfer (START) domain proteins. Unlike multidomain variants, plant minimal START proteins remain poorly understood.
Integrating structural screening across representative plant species, including *Arabidopsis thaliana, Solanum lycopersicum, Cicer arietinum, Glycine max* and *Oryza sativa*, with experimental validation, we show these proteins specifically bind amphipathic lipids, primarily myristic acid and lysophosphatidylcholine. Detailed analysis via circular dichroism, molecular dynamics simulations, and subcellular localization focused on the *Arabidopsis* protein AT4G14500.
Ligand binding occurs through a bipartite mechanism where conserved basic cavity residues coordinate polar headgroups while hydrophobic tails sit deeper within the pocket. Mechanistically, ligand association induces closure of a lid-like β_5_-β_6_ loop at the cavity entrance, whereas membrane proximity promotes gate reopening and lipid release, supporting a membrane-guided gating mechanism. Consistent with this model, confocal microscopy revealed AT4G14500 associates with key lipid metabolic sites: the endoplasmic reticulum, plastids, Golgi apparatus, and plasma membrane.
Together, our findings establish plant minimal START proteins as bona fide lipid-binding proteins that utilize a bipartite ligand-binding mechanism coupled with membrane-responsive gating dynamics.

Graphical Abstract.
**Proposed model for ligand binding and membrane-associated gating in plant minimal START proteins**. (1) In the apo state, the lid (β_5_–β_6_ loop) adopts an open conformation that permits ligand entry into the hydrophobic cavity. (2) Ligand binding induces closure of the lid, resulting in entrapment of the lipid within the START-domain cavity. (3) Membrane proximity promotes reopening of the loop and facilitates lipid release toward the membrane.

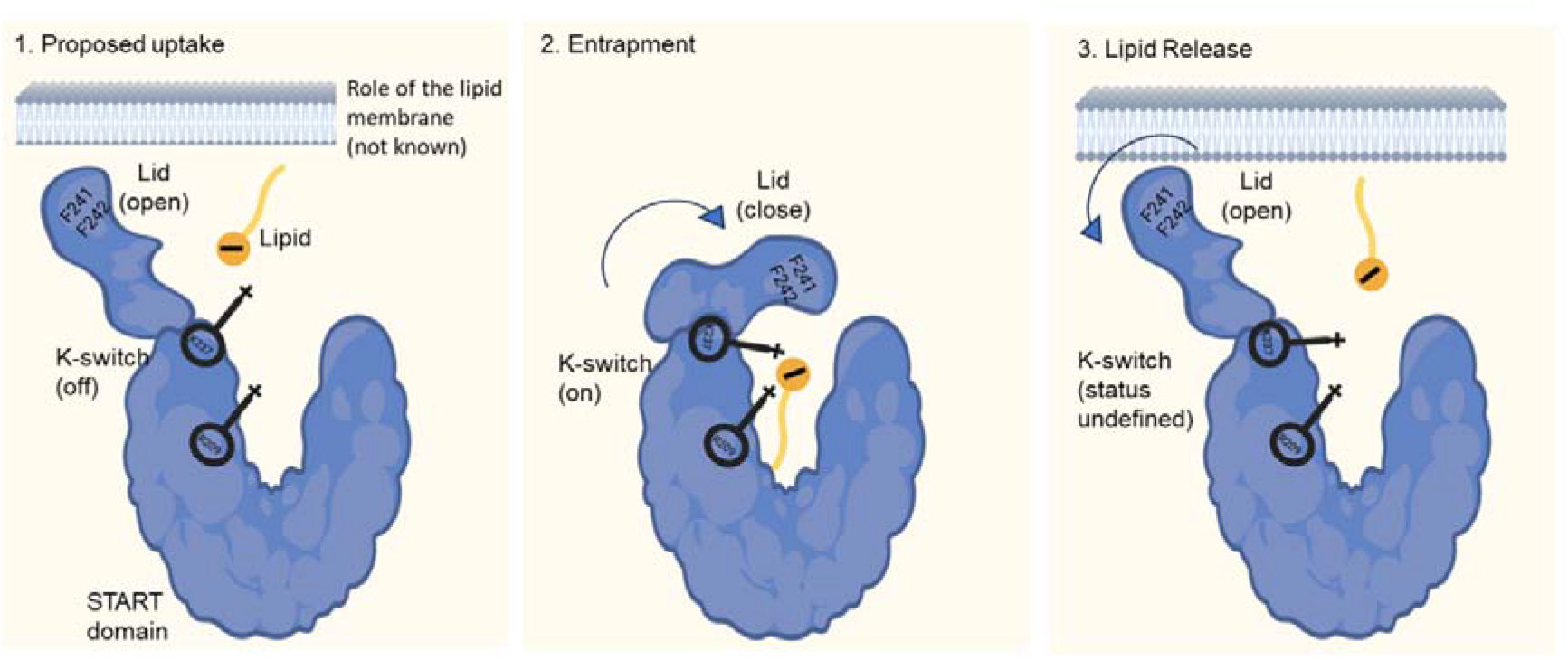

## Introduction

Lipids are essential biomolecules involved in membrane organization, signal transduction, metabolic regulation, and cellular homeostasis (Fahy et al., 2011; Glatz, 2015). They are predominantly hydrophobic and occasionally amphipathic, and are classified by the LIPID MAPS consortium (2005) into eight major categories derived from ketoacyl and isoprene building blocks. However, their predominantly hydrophobic nature prevents free diffusion through the aqueous cellular environment, necessitating specialized lipid transport mechanisms. In eukaryotic cells, lipid biosynthesis occurs across multiple organelles, including the endoplasmic reticulum, Golgi apparatus, peroxisomes, and plastids in plants (Blom et al., 2011; Kim, 2020). Newly synthesized lipids are transported to their destinations through vesicular trafficking, specialized lipid-binding proteins, and membrane contact site (MCS)-mediated transfer (Blom et al., 2011; Prinz, 2010; Wong et al., 2019).

Among lipid-binding modules, the StAR-related lipid transfer (START) domain is an ancient, conserved protein module approximately 210 amino acids in length, characterized by an α/β helix-grip fold that creates an internal hydrophobic lipid-binding cavity (Ponting and Aravind, 1999; Tsujishita and Hurley, 2000). START domains belong to the broader StARkin (*Kin* of *St*eroidogenic *A*cute *R*egulatory protein) superfamily, which is present in all domains of life, and has members like the Bet v1 family and the START family (StAR-related lipid transfer) (Wong et al., 2019; Wong and Levine, 2016). START domain-containing proteins may either consist solely of the START domain, referred to as minimal START proteins (minimal STARTs), or exist as multidomain proteins in combination with other functional domains, making it easier to predict the biological role for later, in contrast to the minimal STARTs (Kanno et al., 2007). In the present study, proteins containing only the START domain were categorized as minimal STARTs, encompassing both strictly minimal proteins and those with putative transmembrane (TM) motifs (TM-STARTs) (*e*.*g*., At4G14500) (Schrick et al., 2004), since these TM motifs are secondary structural elements rather than additional independent functional domains.

Originally identified in mammals, START domains are critical in processes such as steroidogenesis (Caron et al., 1997; Lin et al., 1995), lipid transport, and lipid signaling (Ponting and Aravind, 1999). START domains are most extensively characterized in mammals, where 15 members (STARD1–STARD15) have been identified. Based on sequence and structural similarities, as well as ligand specificities, these proteins are classified into six subfamilies (Clark, 2020). Among the 15 mammalians START domain proteins, STARD1 (also known as StAR), STARD2 (also known as PCTP, Phosphotidylcholine transfer protein), and STARD4–7, along with STARD10, are minimal START proteins (Clark, 2020). STARD1, and STARD4–6 primarily function as sterol-binding proteins (Soccio et al., 2002); notably, STARD1 mediates the transport of cholesterol to mitochondria (Clark et al., 1994; Epstein et al., 1989), while STARD5 mediates the transport of 25-hydroxycholesterol (Rodriguez-Agudo et al., 2005) and can also bind bile acids (Letourneau et al., 2012). In contrast, STARD2, STARD7, and STARD10 are known to bind and transport phosphatidylcholine (Floris et al., 2019; Horibata et al., 2017; Nicholls et al., 2017). The remaining mammalian START proteins, including STARD3, STARD8, STARD9, STARD12, STARD13, STARD14, and STARD15, occur predominantly as multidomain proteins associated with diverse cellular functions such as cholesterol transport, tumor suppression, lipid metabolism, and mitosis, although the ligands and precise roles of several of their START domains remain poorly understood (Frey et al., 2025; Han and Cohen, 2012; Senese et al., 2015; Swarbrick et al., 2014; Torres et al., 2011; Wilhelm et al., 2017). Additional evidence for the ligand diversity of START domains comes from the silkworm *c*arotenoid-*b*inding *p*rotein (CBP1), which binds carotenoids (Slonimskiy et al., 2022; Sluchanko et al., 2022a), and mouse ceramide transfer protein (CERT), which transports ceramide through its START domain (Hanada, 2004; Hanada et al., 2007).

Unlike mammals, START domain proteins have undergone extensive expansion and diversification in plants, with 35 and 29 members identified in *Arabidopsis* and rice, respectively. Genome-wide analyses across Arabidopsis and the rice pangenome further suggest that this expansion occurred through extensive duplication events, highlighting their functional importance during plant evolution and domestication (Mahtha et al., 2021; Schrick et al., 2004). Plants possess both minimal and multidomain START proteins; the majority are multidomain, comprising 26 of 35 in Arabidopsis and 22 of 29 in rice. The majority of multidomain plant START proteins are uniquely associated with homeodomain-leucine zipper (HD-ZIP) transcription factors, a combination absent in animals (Schrick et al., 2004). This association is proposed to couple lipid sensing with gene regulation and is implicated in processes related to cell differentiation during plant development (Mukherjee et al., 2022; Nagata and Abe, 2021). Plants also possess START domains linked with Pleckstrin Homology (PH) and Domain of Unknown Function 1336 (DUF-1336)-containing proteins, further diversifying the function of START domain in lipid signaling and plant defense (Lenoir et al., 2015; Schrick et al., 2004; Tang et al., 2005). HD-START proteins localize to the nucleus, whereas PH-associated START proteins are predicted to localize in mitochondria (Li et al., 2022; Schrick et al., 2004; Yu et al., 2008). Structurally, several multidomain HD-ZIP START proteins possess reduced or highly modified lipid-binding cavities, suggesting an evolutionary shift toward regulatory functions rather than direct lipid transport (Mahtha et al., 2023).

In contrast, minimal START proteins in plants remain the least-studied class of START-domain proteins. Many show similarity to mammalian STARD2, and some also contain predicted transmembrane motifs. These proteins are proposed to function as lipid transporters or mediators of protein interactions, but unlike multidomain START proteins, they lack additional domains that could provide clues about their functions. As a result, the ligands, structural dynamics, and biological roles of most plant minimal START proteins remain poorly understood, leaving a major gap in our understanding of their biology. To address these questions, we carried out a comparative analysis of minimal START proteins across diverse plant lineages. We examined their ligand specificity, cavity architecture, conformational dynamics during ligand binding and release, and subcellular localization. Our findings reveal that despite substantial sequence diversity, plant minimal START proteins share conserved ligand-recognition features and suggest a possible membrane-associated gating mechanism.

## Materials and Methods

### Diversity of Minimal START Proteins

A comprehensive genome-wide dataset of plant START-domain proteins from 62 representative plant species was used (http://nipgr.ac.in/START_FiT). Amino acid length distribution was analyzed using a density histogram in R (version 4.4.1), and sequences of 200–220 amino acid range were selected for downstream analyses. Multiple sequence alignment was performed using MUSCLE v5 (Edgar, 2022). Phylogenetic relationships were inferred using the maximum-likelihood method in RAxML (Kozlov et al., 2019) with 1,000 bootstrap replicates, and the tree was visualized using FigTree v1.4.2 (http://tree.bio.ed.ac.uk/software/figtree/). Sequences were clustered using CD-HIT (Huang et al., 2010) at a 50% sequence identity threshold. Representative proteins were selected based on phylogenetic placement, sequence divergence, and species representation for further analyses.

### Cloning, Expression, and Protein Purification

Minimal START domains from nine genes were codon-optimized for expression in *Escherichia coli* (Biomatik), cloned into the pUC57 (Thermo Scientific™), and subcloned into the pET28a-SUMO (Novagen) using BamHI and XhoI sites (SI Table 1). Site-directed mutagenesis was performed using the QuickChange protocol (SI Table 2), and expression conditions were optimized as described in the Supplementary Methods (SI Table 3). Proteins were expressed in *E. coli* and purified by HisTrap HP affinity chromatography following cell lysis and clarification. Detailed expression, lysis, and affinity-purification conditions are provided in the Supplementary Methods.

For AT4G14500, the His-tag was removed by SUMO protease digestion, followed by purification using a second HisTrap HP and size-exclusion chromatography on a Sephacryl 16/60 S-100 columns (Cytiva). Proteins were verified by 15% SDS-PAGE and concentrated using Amicon® Ultra centrifugal filters (10 kDa MWCO). Protein-lipid overlay assays used His-tagged proteins, whereas all other assays used tag-free AT4G14500. Detailed size-exclusion chromatography and oligomeric-state determination procedures are provided in the Supplementary Methods.

### Lipid Handling and Preparation

For protein-lipid overlay assays, Myristic acid (MA), decanoic acid, dodecanoic acid, sphingomyelin, 2-aminooctanoic acid, and N-acetyl-L-methionine were prepared as 12.5 mM working stocks using appropriate aqueous or organic solvent mixtures. Lysophosphatidylcholine (LPC), a mixture of LPC species (Sigma-Aldrich, PHR2631) was used unless otherwise specified. LPC was prepared as a nominal 12.5 mM stock, with concentration calculations based on C_10_H_22_NO_7_P as the reference species. For overlay assays, 2 µL of each lipid stock was spotted onto the membrane. The same stocks were used for fluorescence assays and subsequently diluted in assay buffer, whereas LPC stocks for BLI were diluted in sodium phosphate buffer containing 150 mM NaCl. Detailed lipid preparation and handling procedures are provided in the Supplementary Materials and Methods.

### Protein-Lipid Overlay Assay

Lipids were spotted onto 4 cm × 2 cm nitrocellulose membranes (Amersham™ Protran® Western blotting membrane) and air-dried for 30 min at room temperature. Also, variable-length LPC strips (seven variants) were customized by Echelon Biosciences for the indicated experiment. Membranes were blocked with 10 mL blocking buffer (5% skimmed milk in TBS: 50 mM Tris-HCl pH 8.5, 150 mM NaCl) at 4 °C for 3 h on a rocker, followed by incubation with purified protein (120 μg mL^−1^ in TBS) at 4 °C for 16 h. Membranes were washed three times with TBST (TBS, 1% skim milk, 0.1% Tween 20) for 10 min at 4 °C. Anti-His HRP-conjugated antibody (1:2000 in TBS; Santa Cruz) was then added and incubated at 4 °C for 4 h, followed by three additional TBST washes. Membranes were rinsed with TBS, developed with freshly prepared Bio-Rad Luminol/Peroxidase solution (1:1) in the dark, and imaged using the Bio-Rad gel documentation system (Blot/Chemiluminescence module). Purified His-tagged protein and sphingomyelin 22 served as positive and negative controls, respectively.

### Circular Dichroism

Circular Dichroism (CD) spectroscopy was performed using a Jasco J-815 spectropolarimeter to analyze wild-type AT4G14500 and mutant proteins. Far-UV spectra (190–260 nm) were recorded at a scanning speed of 100 nm min^−1^, DIT of 4 s, 1 nm bandwidth, and 25 accumulations. Protein samples (0.5 mg mL^−1^) were prepared in phosphate buffer (50 mM phosphate buffer pH 8.5, 150 mM NaCl). Spectra were recorded after buffer blank subtraction and analyzed for characteristic secondary-structure features.

### Fluorescence Spectroscopy

Protein-ligand interactions were monitored by internal fluorescence using an Agilent Technologies Cary Eclipse Fluorescence Spectrophotometer with Cary Eclipse v1.2(146) software. AT4G14500 and mutant proteins (2 μM in 500 μL of 20 mM Tris-HCl, 150 mM NaCl) were analyzed in a 10 mm quartz cuvette. Samples were excited at 290 nm, and emission spectra were recorded from 300-500 nm and averaged over 10 scans. MA, LPC mix, and decanoic acid were titrated from 2-22 μM. Measurements were performed at 25 °C, and buffer blanks were subtracted from all spectra.

### Biolayer Interferometry (BLI)

Binding affinity of wild-type AT4G14500 and its mutant proteins for LPC mix was measured using BLI on an Octet RED96 system (Pall ForteBio). Proteins were biotinylated at 1 mg mL^−1^ using a 1:20 protein-to-biotin molar ratio for 45 min at room temperature. Excess biotin was removed by buffer exchange, and the biotinylated proteins were immobilized on Super-Streptavidin (SSA) biosensors. LPC was prepared in six serial dilutions, with a maximum concentration of 1.25 mM, in 20 mM phosphate buffer (pH 8.5) containing 150 mM NaCl. Association and dissociation kinetics were monitored under standard BLI conditions, and data were analyzed after subtraction of buffer-reference signals. The equilibrium dissociation constant (K_d_) was calculated using Octet BLI Analysis software version 12.2.

### Protein Modeling and Molecular Docking

Tertiary structures of the nine minimal START proteins were predicted using AlphaFold2 (Jumper et al., 2021), and cavity volumes were analyzed using the CASTp server (Tian et al., 2018). Ligands representing diverse lipid families were selected based on the LIPID MAPS classification system, with emphasis on small-to medium-sized plant-relevant lipids and selected plant hormones (SI Table 4). Molecular docking was performed using PyRx v0.8 (Dallakyan and Olson, 2015), which integrates AutoDock Vina, using the AlphaFold2-predicted structures as receptors. Blind docking was performed over the entire protein, and the resulting binding poses were ranked according to predicted binding affinity. Detailed protein and ligand preparation, docking parameters, and pose analysis are provided in the Supplementary Materials and Methods (SI Table 5 and SI Table 6).

### Molecular Dynamics Simulations

AT4G14500 apo and ligand-bound structures obtained after docking were subjected to molecular dynamics simulations using the Desmond package in the Schrödinger suite v2025-02 (Desmond Molecular Dynamics System, D. E. Shaw Research, New York, NY, 2024. Maestro-Desmond Interoperability Tools, Schrödinger, New York, NY, 2025). Protein and protein-ligand complexes were prepared using the Protein Preparation Wizard. Systems were solvated in an orthorhombic TIP3P water box with a 10 Å buffer and neutralized with Na^+^ or Cl^−^ ions at 150 mM. Simulations employed the OPLS4 force field (Lu et al., 2021) and the default Desmond equilibration protocol. Runs were performed in the NPT ensemble at 300 K and 1.01325 bar for 400 ns, with coordinates saved every 400 ps, yielding 1000 frames per simulation. The PubChem CIDs of the ligands used for MD simulations and subsequent *in silico* analyses are 11005 (MA) and 5311264 (LPC).

### Metadynamics

Representative frames at 100 ns from the MA- and LPC-bound simulations were used as starting configurations for metadynamics simulations. Two distinct setups were prepared: a solvated system and a membrane-anchored system, where the protein was embedded in a model lipid bilayer (POPC, POPE, DMPC) with the β_5_-β_6_ loop oriented proximal to the membrane surface. Metadynamics was performed using the Desmond module. The Collective Variable (CV) was defined as the distance between the Center Of Mass (COM) of the protein and the COM of the ligand. The simulation was conducted for 5 ns with an initial Gaussian height of 0.03 kcal/mol and a width of 0.15 Å, deposited every 10 ps with an upper wall at 60 Å. For the bilayer systems, the OPLS_2005 force field was utilized to ensure consistent treatment of lipid-protein interactions (Banks et al., 2005; Shivakumar et al., 2010). The metadynamics simulations yielded a one-dimensional Free-Energy Surface (FES) along the ligand-protein COM distance.

### Confocal Microscopy

The AT4G14500 coding sequence was amplified from *Arabidopsis thaliana* leaf cDNA and cloned into the pENTR/D-TOPO vector (Invitrogen), and recombined into the Gateway destination vector pK7WGF2 to generate an N-terminal GFP-tagged AT4G14500 fusion. Transient expression in *Nicotiana benthamiana* leaves was performed using *Agrobacterium tumefaciens* strain GV3101. GFP-tagged AT4G14500 was co-expressed with fluorescent organelle markers and the P19 silencing suppressor. Organelle markers for the endoplasmic reticulum, Golgi apparatus, peroxisomes, mitochondria, plastids, plasma membrane, and nucleus were used as previously described (Martin et al., 2009; Nelson et al., 2007). Plants were examined 48 h after infiltration using a Leica TCS SP8 confocal microscope. Co-localization was quantified using the Coloc 2 plugin in Fiji by calculating Pearson’s Correlation Coefficient (PCC). Detailed cloning, infiltration, imaging, and image-analysis conditions are provided in the Supplementary Materials and Methods.

## Results

### Plant Minimal START Protein Diversity

The limited understanding of ligand specificity in plant minimal START proteins prompted us to undertake a systematic sequence-based analysis. This analysis was performed on a dataset of 562 minimal START domain sequences from 62 plant species to examine their diversity landscape. An analysis of domain lengths revealed a non-uniform distribution, with 291 sequences falling within a narrow range of 200–220 amino acids, indicating a dominant size class among plant minimal START proteins. Incidentally, this corresponds to the typical length of the START domain (Alpy and Tomasetto, 2005). To focus on this predominant group and reduce variability from potentially truncated variants or large insertions, we further restricted the analysis to the 291 sequences in the 200–220 amino acid range, representing the core population of plant minimal START proteins. Alignment of 291 minimal START proteins highlighted conserved residues and motif-like regions shared across diverse clades (SI Figure 1).

Sequence diversity among the filtered sequences was assessed using a dual approach: redundancy reduction with CD-HIT at a 50% sequence identity cutoff, and in parallel, evaluation of evolutionary relationships through phylogenetic analysis. CD-HIT clustering at 50% identity resulted in 26 distinct clusters. The distribution of sequences across these clusters was markedly uneven. Most clusters (23 clusters) were sparsely populated, containing only 1–7 sequences, whereas only three clusters contained 22, 54, and 175 proteins, respectively. Clustering at 40% or 60% sequence identity resulted in either the merging of distantly related groups (17 clusters) or the over-fragmentation of closely related groups (40 clusters), supporting the use of 50% as a balanced threshold. In parallel, phylogenetic analysis of the 291 sequences resolved the minimal START family into distinct major clades (Figure 1A). Mapping taxonomic divisions onto the phylogenetic tree revealed that proteins from diverse plant lineages, including angiosperms, gymnosperms, pteridophytes, bryophytes, and chlorophytes, were distributed across multiple clades rather than forming lineage-specific groups. This broad distribution suggests that the major minimal START protein groups originated early and diversified prior to the evolutionary separation of major plant divisions.

**Figure 1.**
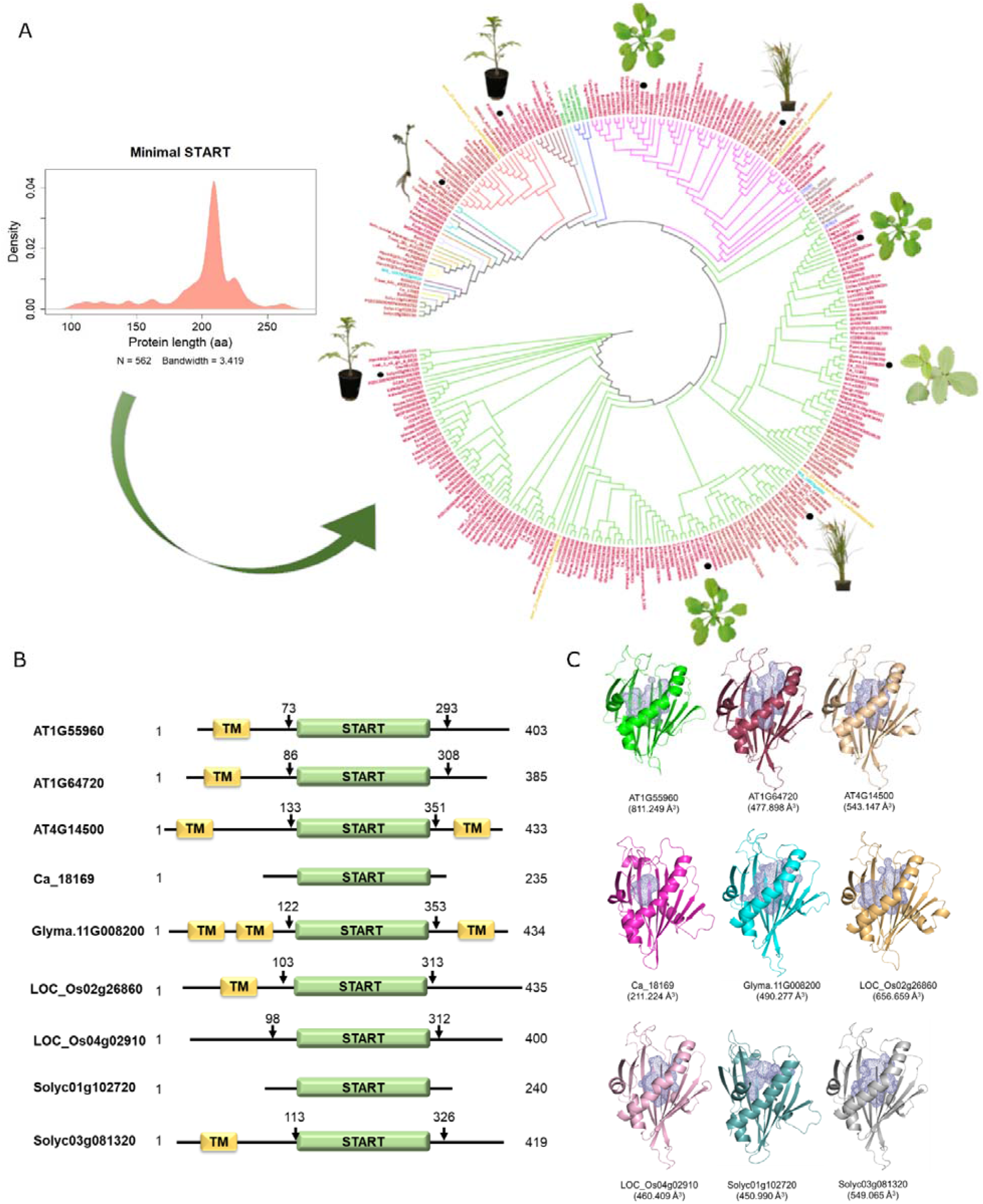
Diversity in plant minimal START proteins and their structural organization. **(A)** Length distribution and phylogenetic analysis of minimal START domain–containing proteins. The length distribution of minimal START domains was estimated using kernel density estimation with a Gaussian kernel and a fixed bandwidth of 3.419. The resulting probability density function is normalized such that the total area under the curve equals one. N denotes the total number of sequences analyzed. A phylogenetic tree was constructed for 291 minimal START domain–containing proteins identified from 62 plant species. Distinct clusters obtained from CD-HIT within the tree are indicated by colored branches. Taxa are color-coded according to major plant divisions: Angiosperm (red), Pteridophytes (blue), Bryophytes (brown), Gymnosperms (cyan), Chlorophytes (green), and Magnoliopsida (yellow). All protein accession numbers correspond to entries in the START_FiT database (http://nipgr.ac.in/START_FiT). Nine diverse proteins selected for further analysis are depicted as black dots along with respective plant images. **(B)** Domain organization of representative minimal START proteins. Schematic representation of nine representative minimal START proteins showing the presence and relative position of the START domain (green) and predicted transmembrane (TM) regions (yellow). The proteins include AT4G14500, AT1G55960, and AT1G64720 from Arabidopsis thaliana; Solyc03g081320 and Solyc01g102720 from Solanum lycopersicum; LOC_Os02g26860 and LOC_Os04g02910 from Oryza sativa; Ca_18169 from Cicer arietinum; and Glyma11G008200 from Glycine max. All protein accession numbers correspond to entries in the START_FiT database (http://nipgr.ac.in/START_FiT). **(C)** Three-dimensional structures of representative minimal START domain proteins predicted using AlphaFold2 (Jumper et al., 2021). Models are displayed in cartoon representation to highlight the conserved START fold architecture. The internal ligand-binding cavity is shown as a blue mesh, as identified using PyMOL (The PyMOL Molecular Graphics System, Version 3.0 Schrödinger, LLC.). Protein accession numbers correspond to the Phytozome database (Goodstein et al., 2012). Cavity volumes (indicated in parentheses) were calculated using the CASTp server (Tian et al., 2018). The following proteins are shown with color coding: AT1G55960 (green), AT1G64720 (red), and AT4G14500 (yellow) from *Arabidopsis thaliana*; Ca_18169 (magenta) from *Cicer arietinum*; Glyma.11G008200 (cyan) from *Glycine max*; LOC_Os02g26860 (orange) and LOC_Os04g02910 (pink) from *Oryza sativa*; and Solyc01g102720 (teal) and Solyc03g081320 (gray) from *Solanum lycopersicum*. A stereoview of the superimposed structures is shown alongside, highlighting conservation of the core fold across species. The color scheme is maintained as described above.

Given the limited representation in most clusters, subsequent analysis focused on the three major clusters identified by CD-HIT. Five proteins were chosen from the largest cluster, while two representative proteins were selected from each of the remaining clusters. While all nine proteins contained a conserved START domain, variation was observed in the presence and positioning of predicted transmembrane (TM) motifs (Figure 1B). Notably, these additional features lie outside the START domain and were not included in the sequence alignment, CD-HIT clustering, or phylogenetic analyses, which were restricted to the conserved START domain region. Alignment of nine representative proteins further showed that the conserved features are maintained across diverse clades, despite sequence divergence in surrounding regions (SI Figure 2). In summary, this resulted in a final set of nine representative minimal START proteins, encompassing both inter- and intra-clade diversity, which were subsequently subjected to further analysis.

### Structural conservation and cavity diversity in minimal START proteins

The representative minimal START proteins identified across diverse clades were analysed at the structural level using AlphaFold2-generated three-dimensional models. All the models exhibited high confidence, with pLDDT scores ranging from 89.46 to 96.08, indicating reliable structural predictions (SI Table 1). All proteins adopted a conserved START domain fold, characterized by a central cavity (Figure 1C. SI Figure 3). All nine proteins superimposed with an rmsd varying from 0.354 Å (LOC_Os04g02910) to 1.778 Å (Ca_18169) with respect to AT1G55960. All nine structures depict a typical α/β helix-grip fold, characterized by a curved antiparallel β-sheet that grips a long C-terminal α-helix. The curved β-sheet composed of nine strands (β_1_-β_9_) and the concave face of the β-sheet form a part of the hydrophobic cavity. A loop connecting β_5_ and β_6_ strands (referred to as β_5_-β_6_ loop) sits at the opening of the cavity (SI Figure 4A). The α/β helix-grip architecture of START domains observed in the AlphaFold2 models is further supported by circular dichroism analysis of AT4G14500. The CD spectrum shows characteristic minima at ~208 and ~222 nm, indicative of α-helical content, along with a broader contribution in the ~215–218 nm region consistent with β-sheet structure, thereby supporting a mixed α/β architecture characteristic of the canonical START-domain fold (SI Figure 4B). Visualization of the models revealed that a cavity is consistently present across all proteins. Quantitative analysis of cavity volumes indicated variability among the proteins, with values ranging from ~211 Å^3^ to ~811 Å^3^ (Figure 1C). While there is significant sequence divergence (SI Figure 2), overall fold remains conserved (SI Figure 3); most of the variation is observed in cavity sizes, suggesting that minimal START proteins of plant origin may accommodate ligands of differing sizes or properties. In summary, these proteins share a conserved fold and cavity architecture, but exhibit variability in cavity dimensions, suggesting differential ligand specificities and prompting a systematic exploration of potential ligands.

### Selection of a diverse ligand set for minimal START proteins

The conserved fold of minimal START proteins, together with variability in cavity dimensions and the absence of a defined ligand for plant minimal START proteins, motivated the rational selection of representative molecules for further analysis. A diverse panel of lipids and small molecules was therefore curated to represent major classes of plant-associated metabolites, with emphasis on hydrophobic or amphipathic molecules compatible with the predominantly hydrophobic cavity. Ligands were selected based on their molecular size, guided by the range of cavity volumes, with preference given to plant-derived lipids involved in membrane structure, signalling, and stress responses. This approach ensured the selection of physiologically relevant ligands that could be accommodated within the minimal START protein cavities (SI Table 4). Additionally, a few synthetic or non-native ligands were also included as probes, based on prior reports (Schrick et al., 2014), to evaluate intrinsic ligand recognition and cross-kingdom binding potential. The final ligand set encompassed representatives from fatty acyls, glycerolipids, glycerophospholipids, sphingolipids, sterols, prenols, and polyketides, capturing a broad chemical space relevant to plant biology. Structure-based docking analysis was performed for all selected lipid molecules to evaluate their compatibility with the START domain cavity. In parallel, a subset of plant hormones was included, and, given their established biological roles and relatively small size, these compounds were evaluated directly using biochemical assays without prior docking-based filtering (SI Figure 5A). Collectively, this ligand library captures the chemical diversity of potential minimal START protein ligands and provides a framework for subsequent *in silico* docking to evaluate their compatibility with the START domain cavity (Figure 2A).

**Figure 2.**
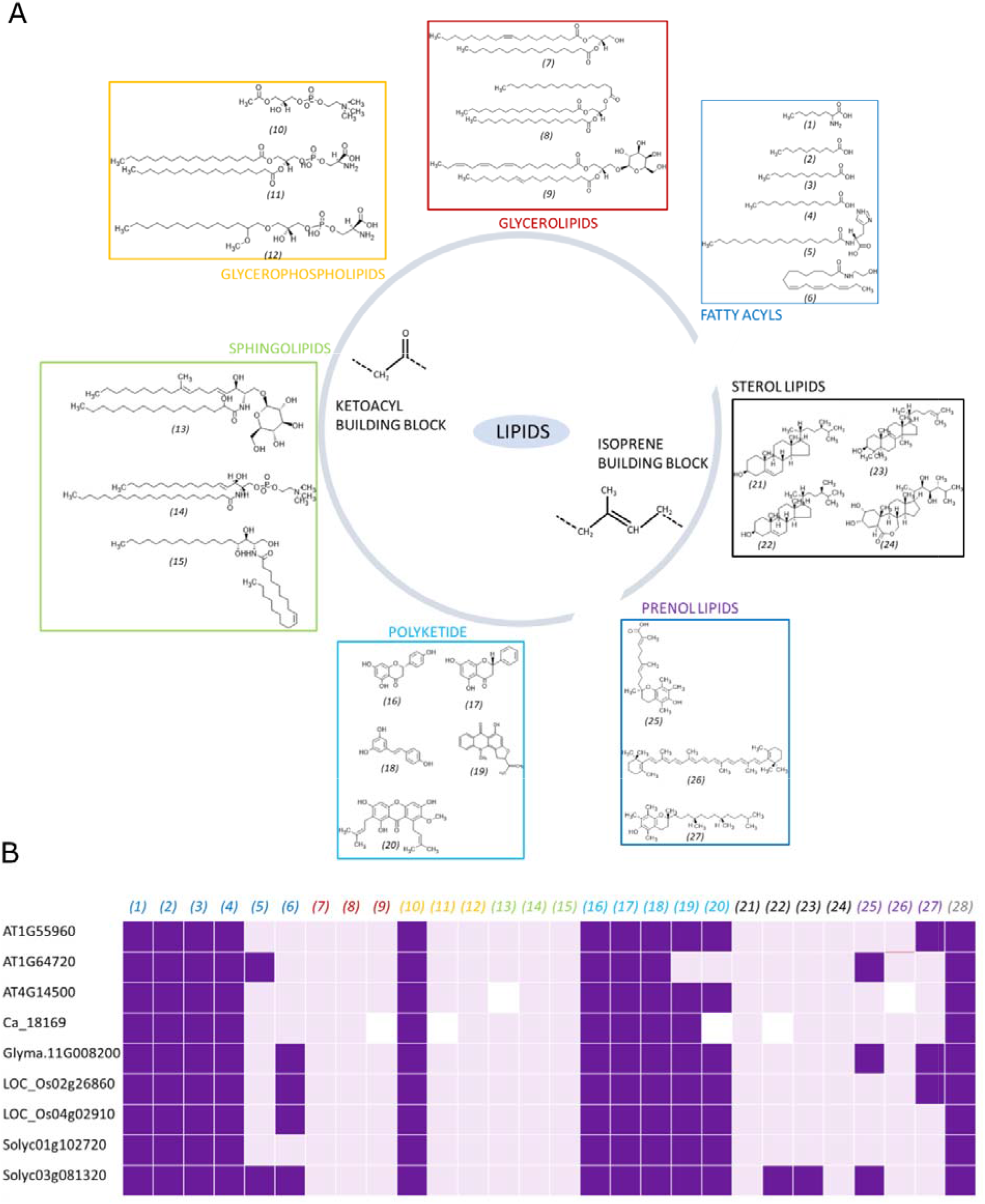
Diversity of lipid ligands and their interaction with minimal START proteins. **(A)** Representative lipid molecules spanning major classes derived from ketoacyl and isoprene building blocks used in docking studies. **(B)** Heatmap summarizing *in silico* docking results of minimal START proteins against the lipid panel. Ligands are numbered as in (A), and the numbering is color-coded according to lipid class. Dark purple indicates ligand binding fully accommodated within the START domain cavity with appropriate stereochemistry; light purple indicates binding with partial cavity occupancy and/or distorted stereochemistry; white indicates no cavity-compatible binding. Fatty acyls: *(1)* 2-aminooctanoic acid (CID: 69522), *(2)* decanoic acid (CID: 16217385), *(3)* dodecanoic acid (CID: 3893), *(4)* myristic acid (CID: 11005), *(5)* N-octadecanoyl-histidine (CID: 152252), *(6)* α-linolenoyl ethanolamide (CID: 5283449). Glycerolipids: *(7)* diacylglycerol (CID: 5283471), *(8)* triacylglycerol (CID: 11146), *(9)* monogalactosyldiacylglycerol (CID: 90657729). Glycerophospholipids: *(10)* lysophosphatidylcholine (LPC 2:0, CID: 5311264), *(11)* phosphatidylserine (CID: 9547096), *(12)* glycerophosphoserine (CID: 137323940). Sphingolipids: *(13)* cerebroside B (CID: 11498616), *(14)* C22 sphingomyelin (CID: 44260125), *(15)* ceramide 3B (CID: 57378373). Polyketides: *(16)* naringenin (CID: 439246), *(17)* pinocembrin (CID: 68071), *(18)* resveratrol (CID: 445154), *(19)* rutacridone (CID: 11623705), *(20)* mangostin (CID: 5281650). Sterols: *(21)* campesterol (CID: 173183), *(22)* β-sitosterol (CID: 222284), *(23)* lanosterol (CID: 246983), *(24)* 24-epibrassinolide (CID: 3239). Prenols: *(25)* 911-carboxy-α-tocotrienol (CID: 53481535), *(26)* β-carotene (CID: 5280489), *(27)* α-tocopherol (CID: 14985). Miscellaneous: *(28)* N-acetyl-L-methionine (CID: 448580). CID denotes the PubChem identifier.

### Docking-based evaluation of ligand binding

The large and chemically diverse ligand library required a systematic and structure-guided screening approach. *In silico* docking was therefore employed to evaluate potential ligand binding within the START domain cavity. This enabled prioritization of selected candidate binders for subsequent in-depth biochemical and biophysical analyses. While docking results are typically evaluated based on predicted binding affinity, we adopted a structure-guided filtering approach rather than relying solely on docking scores. First, only those ligands that were fully accommodated within the START domain cavity were considered, whereas ligands binding partially or outside the cavity were excluded, irrespective of their predicted binding energies. Second, given the high conformational flexibility of many ligands in the dataset, poses exhibiting distorted stereochemistry, such as unrealistic torsional angles or steric clashes, were excluded. These two criteria were used to prioritize cavity-centric interactions with geometrically consistent, physically realistic binding poses over partial or surface binding (SI Table 6). These criteria do not rule out alternative interactions but prioritize geometrically consistent poses to support downstream analyses. Each binding pose was evaluated individually based on the abovementioned two structural criteria. Although the top-ranked pose corresponds to the highest predicted binding affinity, it was not always stereochemically optimal in our analysis. Applying these criteria, all nine START domains were found to interact with some of the selected fatty acids, lysophosphatidylcholine (LPC), all tested polyketides, and N-acetyl-L-methionine, with additional interactions observed for some sterols and prenols. In contrast, many long-chain lipids, including fatty acids, glycerolipids, glycerophospholipids, and sphingolipids, frequently exhibited distorted hydrophobic chain conformations. Overall, despite differences in cavity size, the START domains showed similar preferences for a common subset of ligands (Figure 2B, SI Figure 6A). Despite lower confidence and flexibility in the regions flanking the core START domain, including the predicted TM motifs (SI Table 1), the full-length model was retained for blind docking. These regions did not alter the predicted ligand-binding site (SI Figure 6B). These prioritized ligands were subsequently subjected to biochemical, biophysical, and molecular dynamics analyses to validate docking predictions and to further characterize ligand interactions.

### Experimental and computational validation of ligand interactions

As docking provides a predictive basis for ligand binding, experimental and computational validations were conducted using a subset of ligands identified by docking, including fatty acids, LPC, and N-acetyl-L-methionine. These ligands were prioritized as representative, chemically distinct candidates that consistently exhibited binding within the hydrophobic cavity across multiple minimal START domains and were well-suited for biochemical and biophysical assays. Although polyketides also showed consistent binding, and interactions with sterols and prenols were observed in some cases, they were not included in this study due to their greater hydrophobicity and associated challenges in ensuring consistent assay behavior, warranting future specialized characterization. Additionally, sphingomyelin was included as a negative control. Sphingomyelin shares key physicochemical features with selected ligands, including a zwitterionic headgroup and a hydrophobic tail, it did not exhibit favorable binding in docking. Its inclusion therefore allowed assessment of ligand-recognition specificity beyond general amphipathic or zwitterionic characteristics.

All nine representative START domains were cloned, expressed, and purified (SI Table 3). Of these, six proteins expressed well and were purified in His-tagged form. As an initial qualitative screen for potential lipid ligands, His-tagged proteins were evaluated using protein-lipid overlay assays prior to detailed *in silico* and biophysical characterizations. All six proteins consistently interacted with myristic acid (MA) and LPC, while no detectable interactions were observed with dodecanoic acid, decanoic acid, 2-aminooctanoic acid, N-acetyl-L-methionine, or sphingomyelin. The LPC used in both protein-lipid overlay and subsequent fluorescence assays comprised a mixture of species with varying acyl chain lengths, reflecting a more physiologically relevant scenario. Sphingomyelin was included as a negative control to validate the docking predictions experimentally. His-tagged proteins served as positive controls in all protein-lipid overlay assays (Figure 3A, SI Figure 7). Since overlay assays alone cannot distinguish surface association from binding within the internal hydrophobic cavity, site-directed mutagenesis of key active-site residues was used to further assess the binding site. Given the similar lipid preferences observed across all nine minimal START proteins despite differences in cavity size, AT4G14500 was selected as a representative protein for subsequent analyses.

**Figure 3.**
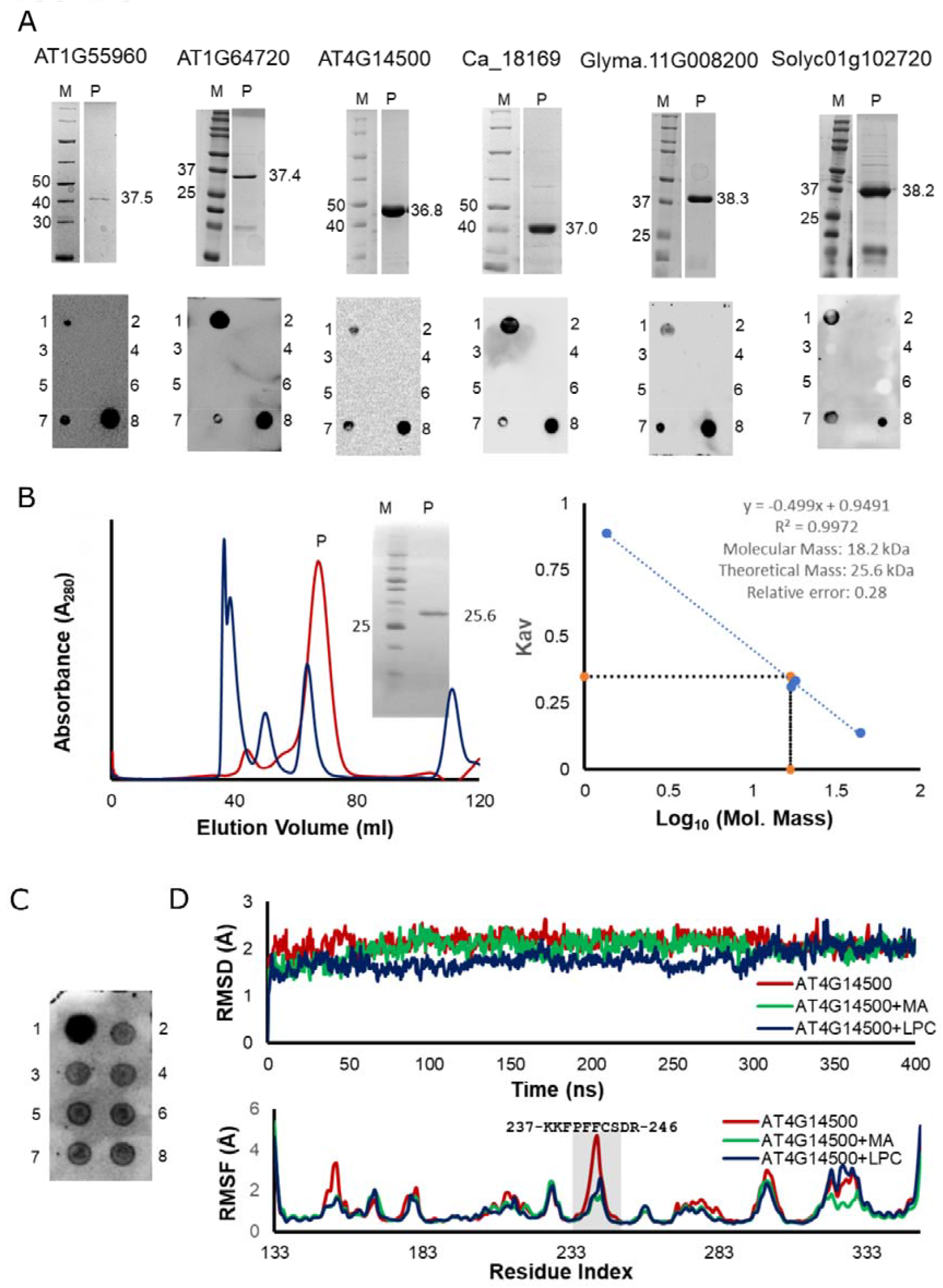
Experimental and computational validation of ligand binding to minimal START proteins. **(A)** Purification and ligand-binding analysis of representative START domains. Upper panels show 15% SDS–PAGE analysis of purified proteins (M: molecular weight marker; P: purified protein), with approximate molecular weights indicated. The lanes corresponding to M and P are from the same SDS-PAGE gel. The lower panels show protein-lipid overlay assays of ligand interactions. Spots correspond to: (1) myristic acid (MA), (2) dodecanoic acid, (3) decanoic acid, (4) 2-aminooctanoic acid, (5) N-acetyl-L-methionine, (6) sphingomyelin, (7) lysophosphatidylcholine (LPC), and (8) positive control (His-tagged protein). **(B)** Oligomeric state analysis of AT4G14500 by size-exclusion chromatography on a Sephacryl S100 gel filtration column (left), showing an elution profile consistent with a monomeric species. Inset: 15% SDS–PAGE of the peak fraction. The red chromatogram corresponds to AT4G14500, and the blue chromatogram corresponds to the standard gel filtration markers. Right: calibration curve used to estimate molecular mass. Kav was calculated as (Ve − Vo)/(Vt − Vo) where Ve, Vo and Vt are elution volume, void volume and total volume, respectively. Thyroglobulin (670 kDa) was used to determine the column’s void volume. **(C)** Protein-lipid overlay assay evaluating binding to glycerophosphocholine and LPC molecules with defined chain lengths. Spots correspond to: (1) positive control (His-tagged protein); (2) glycerophosphocholine (C_8_H_20_NO_6_P; ligand 38; CID: 657272), representing the deacylated LPC headgroup; and (3–8) LPC species with increasing acyl chain lengths: (3) C_14_H_30_NO_7_P (ligand 39; CID: 13917464), (4) C_16_H_34_NO_7_P (ligand 40; CID: 14420997), (5) C_20_H_42_NO_7_P (ligand 41; CID: 460605), (6) C_22_H_46_NO_7_P (ligand 42; CID: 460604), (7) C_24_H_50_NO_7_P (ligand 43; CID: 460602), and (8) C_26_H_54_NO_7_P (ligand 44; CID: 497299). Binding observed across LPC variants, including glycerophosphocholine, highlights the contribution of the phosphocholine headgroup to ligand recognition. **(D)** Molecular dynamics simulations of AT4G14500 in the apo and ligand-bound states. Top: Root mean square deviation (RMSD) plotted as a function of simulation time (400 ns), showing the equilibration and structural stability of the apo protein and the complexes with MA and LPC. Bottom: Root mean square fluctuation (RMSF) profiles showing residue-wise flexibility of AT4G14500 in the apo and ligand-bound states. The β_5_-β_6_ loop (residues 237–246) is highlighted by the shaded region, illustrating localized stabilization upon ligand binding, while the overall protein core remains largely unaffected.

The *in silico* docking analysis was performed assuming that START domains function as monomeric box-like lipid shuttles containing a single hydrophobic binding cavity. However, oligomerization in some proteins can generate additional hydrophobic pockets, either at protein-protein interfaces or between domains (Ahn et al., 2003; Luz et al., 2015; Qiu et al., 2007). This possibility became particularly relevant given that certain ligands were observed to bind partially within the cavity or at the protein surface in docking analyses. To address this, the oligomeric state of AT4G14500 (with the His-tag removed) was examined by size-exclusion chromatography and found to be monomeric in solution (Figure 3B). The presence of a monomeric population supports the relevance of the docking-based analysis.

To address potential bias from the heterogeneous LPC preparation, we performed protein-lipid overlay assays using LPC species with defined acyl chain lengths and the deacylated phosphatidylcholine backbone, glycerophosphocholine. All tested LPC variants bound AT4G14500, and *in silico* analyses consistently positioned them within the protein cavity, indicating that LPC recognition is not restricted to a specific molecular species (Figure 3C; SI Figure 8). Inspired by recent findings on Arabidopsis PROTODERMAL FACTOR2 (Wojciechowska et al., 2024), blind docking of LPC 18:1 and LPC 18:2 similarly placed both ligands in the same cavity (SI Figure 8C). Unlike fatty acids, LPC recognition appeared less dependent on acyl-chain length, potentially reflecting greater flexibility and headgroup-driven recognition, as supported by binding of deacylated glycerophosphocholine (structure 38).

Molecular dynamics simulations for 400 ns were performed to further assess ligand-binding stability (Figure 3D). Analysis of the root mean square deviation (RMSD) profiles showed that all systems: apo AT4G14500 as well as ligand-bound complexes with MA and LPC, rapidly reached equilibrium and remained stable over the course of the simulation. This indicates the overall structural stability of the protein. Notably, no large deviations (assessed by root-mean-square deviation, RMSD) were observed upon ligand binding, suggesting that ligand association does not perturb the global fold of the START domain. Residue-wise flexibility was assessed by root-mean-square fluctuation (RMSF) analysis. Comparable profiles were observed between the apo and ligand-bound states. Localized reductions in flexibility were evident in a few loops, especially the β_5_-β_6_ loop upon ligand binding, while the core structure remained largely unaffected. The molecular basis of these changes is explored in the subsequent section. These observations indicate that both MA and LPC remain stably accommodated within the cavity throughout the simulation, reinforcing the binding modes inferred from docking and biochemical analyses and setting the stage for a detailed examination of the residue-level interactions that underlie ligand recognition and stabilization.

### Structural basis of ligand recognition in START domains

Having established stable ligand binding, we next dissected the residue-level interactions that govern ligand recognition. To identify residues involved in ligand recognition by AT4G14500, docking poses of MA and LPC were analyzed. In the case of MA, the carboxylate group was predicted to interact with Lys237 and Arg209, while the acyl chain was accommodated within the hydrophobic cavity. In contrast, LPC binding involved coordination of the phosphate group by Lys237 and Arg209, with the choline moiety forming a cation–π interaction with Trp289, and the acyl chain similarly occupying the hydrophobic pocket (Figure 4A). To evaluate the stability of these interactions during the simulations, the 400 ns molecular dynamics trajectories were analyzed (Figure 3D). In the MA-bound state, Arg209 maintained interactions with both oxygen atoms of the carboxylate group (~64% and ~41% occupancy), while Lys237 showed weaker engagement. Additionally, Arg252 formed a water-mediated interaction (~75% occupancy). In the LPC-bound state, Arg209 and Lys237 retained interactions with the phosphate group for ~100% and ~56% of the simulation time, respectively, with the Lys237 interaction primarily water-mediated. The predicted cation–π interaction with Trp289 was not maintained; instead, His312 engaged with the choline group (~72% occupancy) (Figure 4B, SI Figure 9).

**Figure 4.**
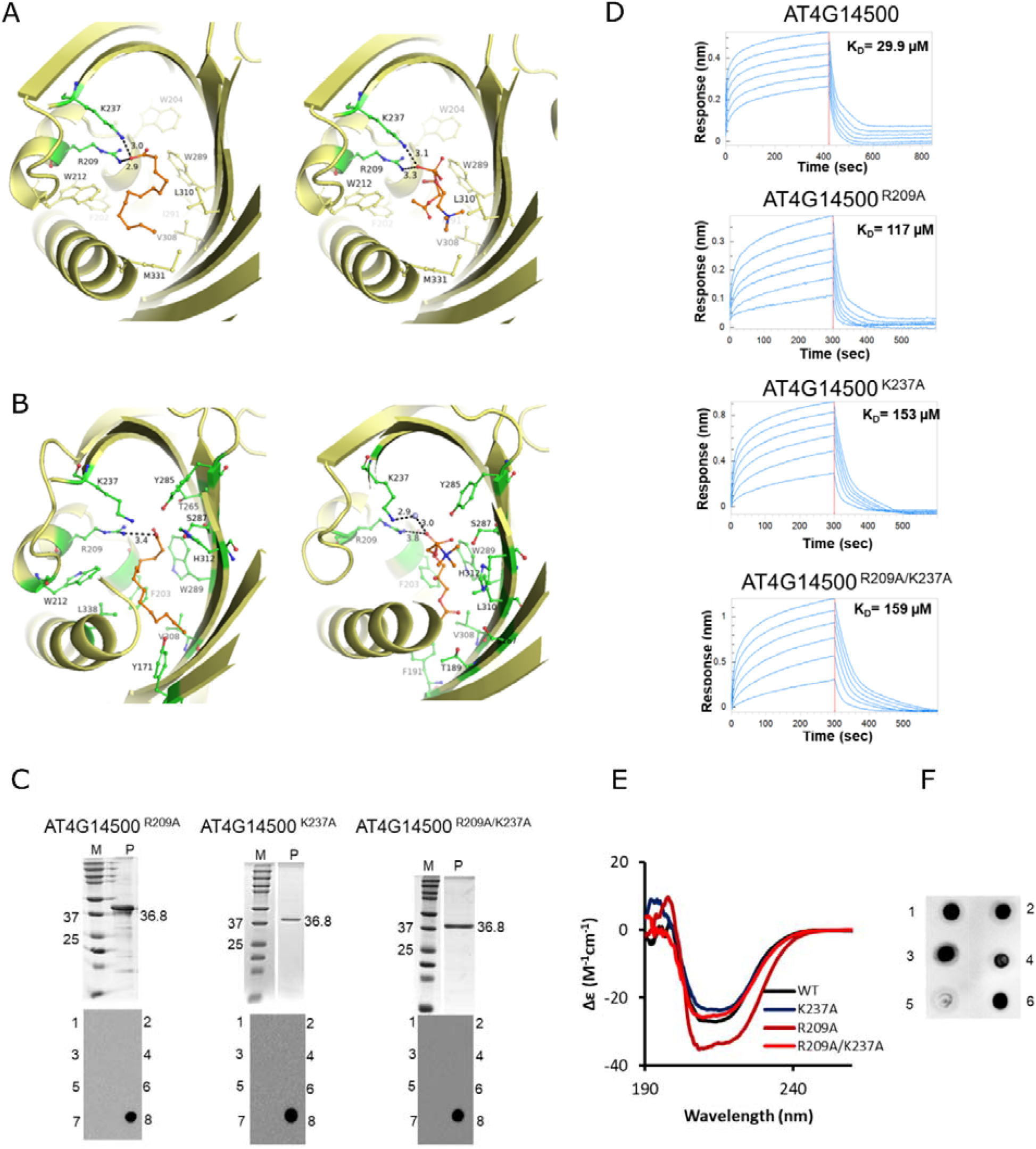
Structural basis of ligand recognition in AT4G14500 START domain. **(A)** Docking-derived models of ligand binding within the START domain cavity. Myristic acid (MA, left) and lysophosphatidylcholine (LPC, right) are shown in stick representation. **(B)** Representative interaction snapshots from molecular dynamics simulations. MA (left) and LPC (right) show stable engagement within the cavity throughout the simulation. **(C)** Biochemical validation of key residues. Purified His-tagged AT4G14500 mutants (R209A, K237A, and R209A/K237A) were analyzed by 15% SDS–PAGE (upper panels; M, molecular weight marker; P, purified protein) and assessed for ligand binding using protein-lipid overlay assays (lower panels). Spots correspond to: (1) MA, (2) dodecanoic acid, (3) decanoic acid, (4) 2-aminooctanoic acid, (5) N-acetyl-L-methionine, (6) sphingomyelin, (7) LPC, and (8) positive control (His-tagged protein). The lanes corresponding to M and P are from the same SDS-PAGE gel for R209A, K237A and R209A/K237A mutants. **(D)** Quantitative binding kinetics measured by bio-layer interferometry (BLI). Representative real-time association and dissociation sensorgrams for LPC binding to wild-type AT4G14500 (K_D_ = 29.9 µM) and cavity mutants (R209A, K_D_ = 117 µM; K237A, K_D_ = 153 µM; and the double mutant R209A/K237A, K_D_ = 159 µM). LPC was tested over a two-fold concentration series ranging from 39.1 µM to 1250 µM. Blue sensorgrams represent the analyte concentration series, fitted using a 1:1 binding model. Equilibrium dissociation constants (K_D_) are indicated for each protein variant. **(E)** Circular dichroism (CD) spectra of wild-type (same as in SI Figure 4B) and mutant proteins. **(F)** Protein-lipid overlay assay with lipid analogues. Spots correspond to: (1) MA, (2) LPC, (3) 14-hydroxymyristic acid, (4) edelfosine, (5) miltefosine, and (6) positive control (His-tagged protein).

Residues Lys237 and Arg209, which were consistently involved in ligand coordination across docking and molecular dynamics simulations, were selected for biochemical validation. Incidentally, both residues are highly conserved across all 291 minimal START proteins (SI Figure 1 and SI Figure 2). Single mutants (K237A and R209A) and a double mutant (K237A/R209A) were generated and evaluated using protein-lipid overlay assays. We failed to purify W289A and H312A mutants. In qualitative protein–lipid overlay assays, the R209A, K237A, and R209A/K237A mutants exhibited a complete loss of binding to all tested lipids (Figure 4C, SI Figure 10). To quantitatively validate these observations, bio-layer interferometry (BLI) was performed using LPC. Quantitative BLI analysis could not be reliably performed with MA because of its poor aqueous solubility. Nevertheless, BLI measurements with LPC revealed a marked 4.0 to 5.3-fold reduction in binding affinity for the R209A (K_D_ = 117 µM), K237A (K_D_ = 153 µM), and double mutant R209A/K237A (K_D_ = 159 µM) compared with wild-type AT4G14500 (K_D_ = 29.9 µM) (Figure 4D), confirming the critical role of these conserved cavity residues in ligand recognition and binding.

To exclude the possibility that loss of binding resulted from structural perturbations, His-tag-free mutant proteins were purified and analyzed for secondary structure (Figure 4E, SI Figure 11). Circular dichroism (CD) analysis showed that mutations at Lys237 and Arg209 did not alter the overall protein fold, indicating that the loss of ligand binding arises specifically from the disruption of interactions with the charged groups of MA and LPC. To further validate the ligand-binding mechanism, we next investigated whether structurally related analogs of MA and LPC could be recognized by the START domain. Two LPC mimetics, Miltefosine (ligand 35) and Edelfosine (ligand 36), along with a MA analog, 14-hydroxymyristic acid (ligand 37), were selected for this analysis (SI Figure 5B, SI Table 4). Protein-lipid overlay assays performed with wild-type AT4G14500 showed that all three analogs exhibited binding (Figure 4F, SI Figure 10B). In contrast, no binding was observed for a panel of plant hormones tested under identical conditions (SI Figure 10C), suggesting that ligand recognition is not promiscuous but is instead selective for lipid-like molecules. Together, these results define a bipartite ligand-recognition mechanism in which conserved basic residues coordinate polar headgroups, while the hydrophobic cavity accommodates lipid tails, thereby enabling selective yet adaptable binding of structurally related ligands. How this binding event is coupled to conformational changes still remains unclear (Figure 3D).

### Conformational gating of the START domain cavity upon ligand binding

We next examined how ligand binding influences conformational dynamics of the START domain. Superimposition of multiple frames from the 400 ns molecular dynamics (MD) simulations revealed striking differences between the apo and ligand-bound states. In the absence of ligand, the β_5_-β_6_ loop exhibited pronounced conformational flexibility, whereas its conformational flexibility was markedly reduced upon binding to either MA or LPC, consistent with the RMSF analysis (Figure 5A, Figure 3D). Interestingly, this loop is positioned at the entrance to the ligand-binding cavity and contains two highly conserved phenylalanine residues, Phe241 and Phe242 (Figure 5B, SI Figures 1 and SI Figure 2). Structural comparison further showed that the loop adopts an open conformation in the apo state, allowing access to the cavity, but transitions to a closed conformation in the ligand-bound state, effectively covering the cavity entrance (Figure 5B). These observations suggest that the β_5_-β_6_ loop functions as a dynamic gate controlling ligand access. To quantify this transition, three residues at the cavity entrance: Phe241 (β_5_-β_6_ loop), Phe214 (loop between α_3_ and β_4_), and Leu325 (α_4_), were used to define a triangular region based on their Cα atoms. The area of this triangle was calculated across the MD trajectories for apo and ligand-bound states. A substantial reduction in the average area was observed in the MA- and LPC-bound simulations (~44.5% and ~45.6%, respectively) compared to the apo form (SI Table 7, SI Figure 12), indicating a significant narrowing of the cavity entrance upon ligand binding. Time-resolved analysis further showed that this reduced cavity entrance was maintained throughout the 400 ns simulations in both ligand-bound systems, whereas the apo protein exhibited pronounced long-timescale fluctuations in entrance geometry (SI Figure 12D), indicating that ligand-induced cavity narrowing is a sustained conformational feature rather than a transient event.

**Figure 5.**
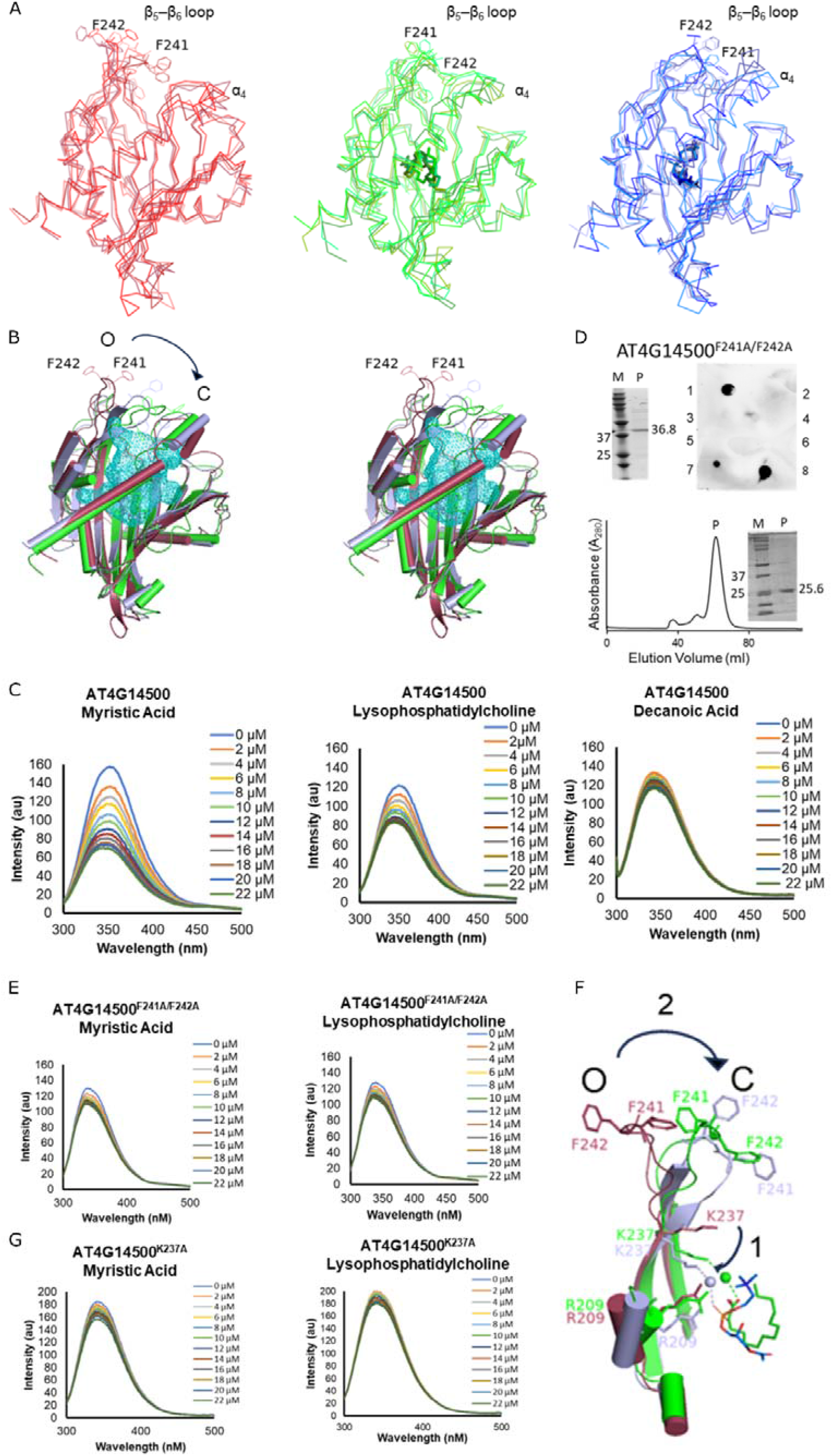
Conformational gating and ligand-induced dynamics of the AT4G14500 START domain. **(A)** Superimposition of representative frames from 400 ns MD simulations for the apo (red), Myristic Acid (MA)-bound (green), and Lysophosphatidylcholine (LPC)-bound (blue) states. The β_5_-β_6_ loop exhibits high flexibility in the apo state but becomes stabilized upon ligand binding. The two ligands are shown in sticks. **(B)** Stereo-view of the structural comparison highlighting the transition of the β_5_-β_6_ loop from an Open (O) conformation in the apo state (red) to a Closed (C) conformation in the ligand-bound state (green/blue), effectively covering the cavity entrance. The cavity has been shown as a mesh. The conserved residues Phe241 and Phe242 are indicated. **(C)** Intrinsic fluorescence spectra of wild-type AT4G14500 titrated with increasing concentrations (0– 22 µM) of MA, LPC, and Decanoic Acid. MA and LPC induce a concentration-dependent quenching of fluorescence, while Decanoic Acid (a non-binder) shows no effect. **(D)** Validation of the AT4G14500^F241A/F242A^ double mutant. Top left: 15% SDS-PAGE of the purified His-tagged mutant protein (M: Marker, P: Protein, protein sizes are in kDa). Top right: Protein-lipid overlay assay confirming that the AT4G14500^F241A/F242A^ mutant retains the ability to bind ligands. Spots correspond to: (1) MA, (2) dodecanoic acid, (3) decanoic acid, (4) 2-aminooctanoic acid, (5) N-acetyl-L-methionine, (6) sphingomyelin, (7) LPC, and (8) positive control (His-tagged protein). Bottom: Size-exclusion chromatography (SEC) profile and SDS-PAGE of AT4G14500^F241A/F242A^ without His-tag (M: Marker, P: Protein, protein sizes are in kDa). **(E)** Intrinsic fluorescence measurements of the AT4G14500^F241A/F242A^ in the presence of MA and LPC. **(F)** Proposed gating mechanism. (1) Ligand binding induces a conformational shift in Lys237, enabling water-mediated interactions with the ligand headgroup. (2) This triggers the closure of the β_5_-β_6_ loop, bringing Phe241/Phe242 into proximity with α_4_ to stabilize the closed state. Apo protein is shown in red, MA-bound protein is shown in green, and LPC-bound protein is shown in blue. **(G)** Intrinsic fluorescence spectra of the AT4G14500^K237A^ mutant in the presence of MA and LPC.

The presence of conserved aromatic residues (Phe241 and Phe242) within the β_5_-β_6_ loop prompted us to investigate whether ligand binding leads to detectable changes in intrinsic fluorescence. The tag-free protein (Figure 3B) was therefore used to monitor interactions with MA and LPC through intrinsic fluorescence measurements. Ligand titration resulted in a concentration-dependent decrease in fluorescence intensity (Figure 5C), whereas decanoic acid, which showed no binding in protein-lipid overlay assays, did not induce any measurable change, supporting the specificity of the response (Figure 5C). To directly assess the contribution of the β_5_-β_6_ loop to this fluorescence signal, a double mutant in which Phe241 and Phe242 were substituted with alanine (AT4G14500^F241A/F242A^) was generated. Interestingly, the mutant does not affect ligand binding (Figure 5D, SI Figure 13A), indicating that these aromatic residues are not essential determinants of ligand recognition. Consistent with this observation, molecular dynamics simulations of the AT4G14500^F241A/F242A^ mutant showed that both ligands remained stably accommodated within the binding cavity despite replacement of the two Phe residues (SI Figure 13B). These findings suggest that ligand binding is maintained through the remaining network of interactions within the binding cavity, whereas Phe241 and Phe242 primarily contribute to the ligand-induced conformational response of the β_5_-β_6_ loop. Consistent with this interpretation, intrinsic fluorescence measurements showed that the mutant failed to exhibit ligand-induced changes in fluorescence intensity (Figure 5E), indicating that the observed fluorescence quenching reflects conformational rearrangement of the β_5_-β_6_ loop rather than direct ligand interaction with Phe241 and Phe242.

While Phe residues on loop β_5_-β_6_ are not essential for ligand binding, ligand binding is coupled to the closure of the β_5_-β_6_ loop. MD simulations suggest that ligand binding induces a conformational change in the Lys237 side chain, enabling water-mediated interactions with the ligand headgroups and inducing conformational changes in the adjacent β_5_-β_6_ loop. This, in turn, brings β_5_-β_6_ loop into closer proximity to alpha helix 4 (α_4_) (SI Figure 4), stabilizing the closed conformation (Figure 5F). Consistent with this, intrinsic fluorescence measurements with the K237A mutant in the presence of MA and LPC showed no detectable change in fluorescence intensity (Figure 5G), in line with its loss of ligand binding. In summary, these findings support a model in which ligand binding induces conformational rearrangements that close the cavity entrance, thereby coupling molecular recognition of ligand to dynamic regulation of ligand access and raising the question of how ligand release is achieved once the cavity is closed.

### Membrane proximity facilitates ligand release

While ligand binding stabilizes a closed cavity conformation, it remains unclear how the ligand is subsequently released from this enclosed state. To investigate the effect of membrane proximity on ligand dissociation, we performed metadynamics simulations using the distance between the center of mass (COM) of the ligand and the COM of the entire protein as the collective variable under two conditions: in the absence of a lipid bilayer and in the presence of model lipid bilayers composed of POPC, POPE, or DMPC. It is important to emphasize that the resulting free-energy profiles describe the free-energy landscape associated with ligand dissociation from the protein rather than the equilibrium thermodynamics of lipid partitioning between water and the membrane. In the absence of a membrane, the free-energy profile exhibited a markedly deeper minimum than the membrane-associated systems, indicating a less favorable free-energy landscape for ligand dissociation along the selected coordinate. In contrast, membrane proximity promoted opening of the β_5_-β_6_ loop, facilitating ligand escape from the binding cavity (Figure 6A). Structural superimposition of representative conformations extracted from the metadynamics trajectories further showed that ligand dissociation is accompanied by membrane-assisted opening of the β_5_-β_6_ loop, supporting a preferred ligand exit region adjacent to the loop (Figure 6B).

**Figure 6.**
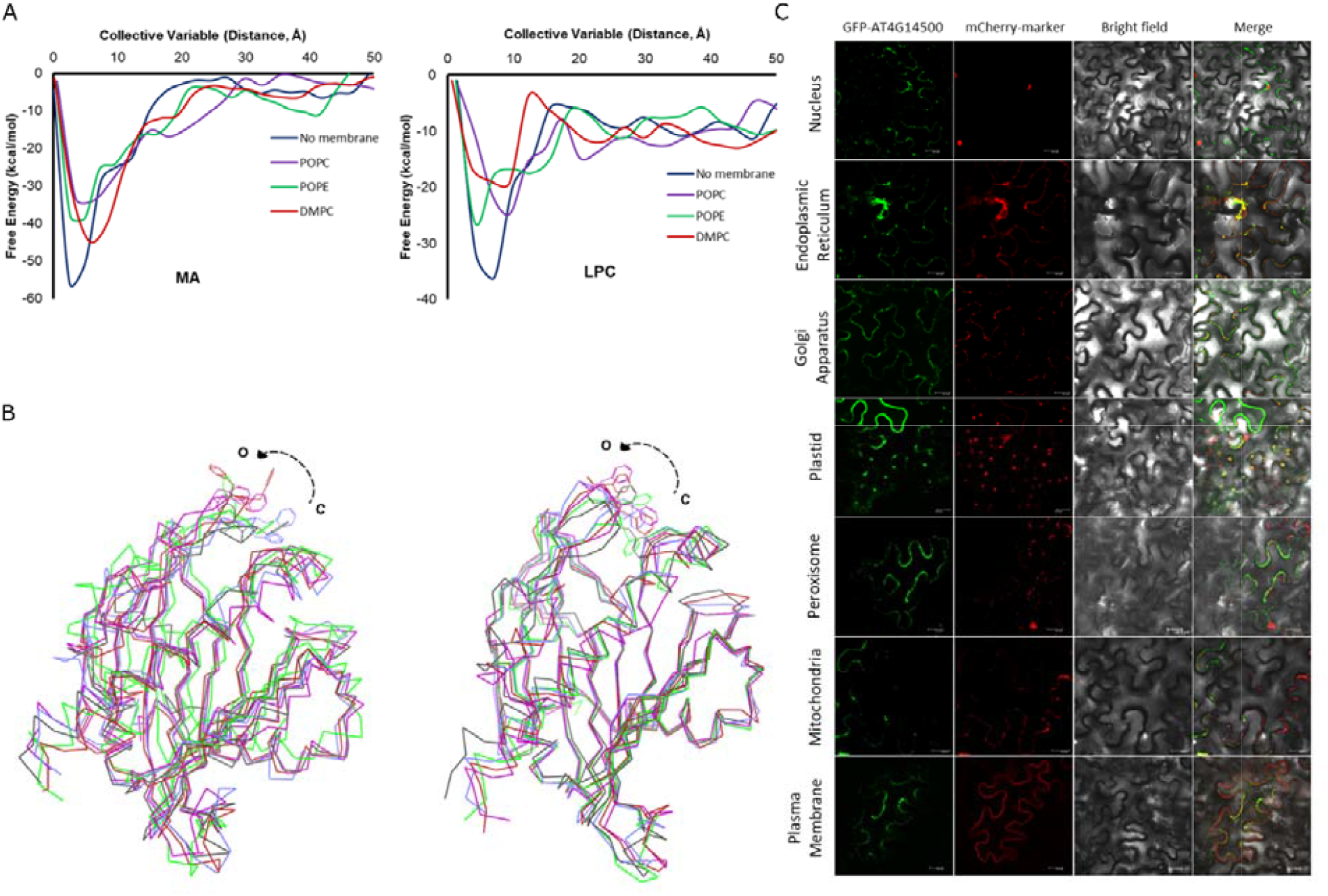
Membrane-Guided Gating and Subcellular Localization. **(A)** Free-energy landscapes of ligand release obtained from metadynamics simulations for myristic acid (MA, left) and lysophosphatidylcholine (LPC, right). The profiles show ligand dissociation along the protein-ligand COM distance in a lipid bilayer-free environment (No membrane) and in the presence of POPC, POPE, and DMPC bilayers. **(B)** Structural superimposition of representative conformations from the metadynamics trajectories for MA (left) and LPC (right). The colored structures represent conformations sampled from metadynamics trajectories following ligand exit in the membrane-free and membrane-associated systems. The black structure represents the initial input ligand-bound closed conformation. The color scheme is the same as in (A). The β_5_-β_6_ loop adopts an open conformation following ligand release, with the positions of Phe241 and Phe242 shown to highlight their location within the loop. Ligands are not displayed to facilitate visualization of the loop conformational changes. **(C)** Organelle-specific localization of AT4G14500. Confocal laser scanning microscopy images showing expression of GFP-tagged AT4G14500 (green) together with organelle-specific mCherry markers (red). Co-localization signals (yellow in merged images) are observed with the endoplasmic reticulum, plastids, Golgi apparatus, and plasma membrane. Quantification of co-localization using Pearson’s correlation coefficient yielded values of 0.65 ± 0.05 (ER), 0.69 ± 0.09 (plastids), 0.61 ± 0.06 (Golgi), 0.58 ± 0.08 (plasma membrane), 0.43 ± 0.11 (peroxisomes), 0.43 ± 0.11 (mitochondria), and 0.25 ± 0.14 (nucleus). Values are presented as mean ± SD from 3–6 independent biological replicates. Scale bars: 20 μm.

To assess the cellular relevance of this mechanism, we performed confocal microscopy– based localization studies. The protein showed preferential localization to the endoplasmic reticulum, Golgi apparatus, plastids, and plasma membrane, consistent with the highest Pearson’s correlation coefficients obtained for these organelles (Figure 6C, SI Figure 14). This distribution is consistent with a model in which ligand exchange is not driven by diffusion through the cytosol but is instead facilitated by membrane proximity, with the β_5_-β_6_ loop acting as a membrane-responsive gate. By localizing to membranes rather than the cytosol, the protein is positioned where its gating mechanism is most effective.

## Discussion

Plant minimal START proteins are the least-studied branches of the StARkin superfamily despite their widespread distribution across plant lineages. Our sequence- and structure-based analyses revealed that minimal START proteins retain a conserved core START-domain architecture. Most proteins fall within the characteristic 200–220-amino-acid range. The close correspondence between sequence clustering and phylogenetic relationships suggests that major minimal START protein groups emerged early during plant evolution and subsequently diversified across multiple plant lineages. Similar conservation of domain size and overall fold is also observed in mammalian START domain proteins (Ponting and Aravind, 1999; Soccio and Breslow, 2003) and other StARkin members, such as the Bet v1 family (Radauer et al., 2008). The amino acid residues lining the cavity are far more conserved than the surface-exposed residues (Figure 7). Such conservation likely reflects structural constraints and evolutionary pressure required to maintain the characteristic α/β helix-grip fold that accommodates hydrophobic ligands. Although the overall fold remained highly conserved, cavity volumes varied substantially among the proteins. This variation is important because ligand specificity in START-domain proteins is closely linked to cavity architecture, and many multidomain plant START proteins possess smaller or partially restricted cavities due to bulkier cavity-lining residues (Mahtha et al., 2023). In contrast, plant minimal START proteins appear to retain comparatively larger and more accessible cavities resembling those of mammalian START proteins.

**Figure 7.**
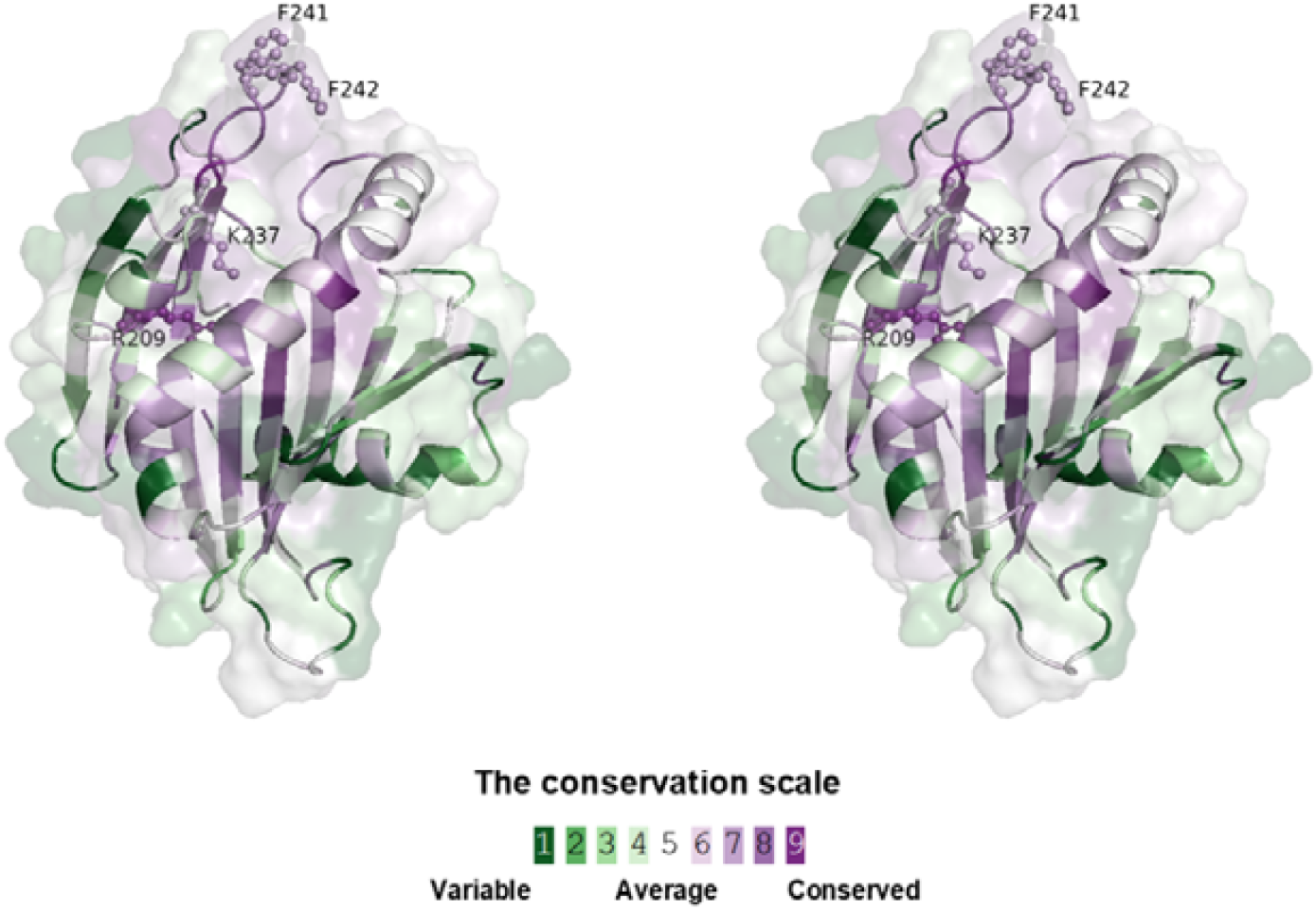
Stereoview representation of ConSurf-based evolutionary conservation mapped onto the Arabidopsis minimal START protein AT4G14500. Surface and cartoon stereoview representations of the predicted three-dimensional structure of AT4G14500 showing residue conservation mapped using ConSurf analysis based on 291 homologous sequences (Armon et al., 2001; Glaser et al., 2003). Residues are colored according to the ConSurf conservation scale, ranging from variable (green) to highly conserved (purple). The β5–β6 loop region (containing F241 and F242) and ligand-interacting residues (R209 and K237) display strong evolutionary conservation. Notably, residues lining the ligand-binding cavity are more conserved than solvent-exposed surface residues, and the β5–β6 loop is considerably more conserved than other loop regions.

Despite substantial variation in sequence and cavity dimensions, the representative plant minimal START proteins displayed remarkably similar ligand-binding behavior. Docking analyses consistently identified a common subset of compatible ligands, particularly fatty acids and lysophosphatidylcholines. Experimental validation further showed that multiple representative minimal START proteins consistently interacted with MA and LPC, suggesting that ligand recognition is not entirely random despite the family’s apparent structural diversity. Specifically, they utilize a common bipartite ligand-recognition mechanism in which conserved basic residues interact with the charged headgroups, while the hydrophobic cavity accommodates lipid tails. In AT4G14500, the highly conserved residues Arg209 and Lys237 coordinated the carboxylate group of MA and the phosphate group of LPC. LPC recognition also involved cation–π interactions with aromatic residues. This bipartite mechanism of ligand recognition in plants’ minimal START proteins mirrors that of mammalian STARD2 (PC-TP), which uses an Arg residue (Arg78) to coordinate the phosphate group of LPC, while the tail occupies the hydrophobic pocket (Druzak et al., 2023; Roderick et al., 2002). Similarly, in STARD14 (Them1), Arg449 anchors the carboxyl headgroup of fatty acids as the acyl chain is accommodated internally (Tillman et al., 2020). This striking alignment of molecular recognition strategies across kingdoms suggests that plant minimal START proteins have retained a conserved structural blueprint for binding amphipathic lipids.

Although these findings highlighted a conserved mechanism for ligand recognition in plant minimal START proteins, they also raise an important mechanistic question: how do amphipathic ligands come out through the deeply buried hydrophobic cavity of the START domain? Our analyses suggest that ligand exchange is regulated by conformational dynamics of the highly conserved β_5_-β_6_ (Ω1) loop positioned at the cavity entrance. In the apo state, this loop adopts an open-close, flexible conformation, whereas ligand binding stabilizes a closed conformation that narrows the cavity entrance. The highly conserved aromatic Phe residues of β_5_-β_6_ loop appear to primarily regulate this gating mechanism rather than directly interacting with the ligand. Metadynamics simulations utilizing simple lipid bilayer models further showed that membrane proximity destabilizes the bound state and promotes ligand release through reopening of the β_5_-β_6_ loop. A similar β_5_-β_6_ (Ω1) loop-mediated gating mechanism has been proposed for several mammalian START domain proteins, including human STARD2 and STARD4 (Iaea et al., 2015; Romanowski et al., 2002). In line with this, apo structures of mammalian START domains frequently display weak electron density or elevated B-factors for the β_5_-β_6_ loop, reflecting its ligand-dependent conformational flexibility as an essential gate (Kudo et al., 2008; Letourneau et al., 2016; Moqadam et al., 2024; Murcia et al., 2006; Sluchanko et al., 2022b; Tan et al., 2019; Tong et al., 2018). Similar lid-like gating mechanisms are also observed in structurally unrelated lipid transport proteins such as Sec14 domains (Ryan et al., 2007; Sha et al., 1998) and non-specific lipid transfer proteins (Missaoui et al., 2022). This suggests that dynamic loop-mediated control of cavity access may represent a convergently evolved strategy for lipid exchange across diverse lipid-binding protein families involved in non-vesicular lipid transport. Together, these observations suggest that plant minimal START proteins may function as lipid-binding proteins, in which selective ligand recognition is coupled to dynamic β_5_-β_6_ loop gating to spatially regulate lipid exchange. Although the precise site and mechanism of lipid uptake remain to be fully defined for plant minimal START proteins, existing paradigms suggest that START domains often associate with lipid membranes or work in tandem with membrane-anchored regions (such as putative TM domains) to facilitate lipid binding and transfer (Tsujishita and Hurley, 2000). The predicted TM regions identified in several plant minimal START proteins may contribute to membrane association, although their functional significance remains unclear and warrants further investigation.

The proposed membrane-guided gating mechanism corroborates with subcellular localization of AT4G14500 to the endoplasmic reticulum (ER), Golgi apparatus, plastids, and plasma membrane. These four cellular compartments represent major sites of lipid metabolism and membrane remodeling. The localization of AT4G14500 to the ER and plastids is consistent with the role of the *Marchantia polymorpha* START protein MpSTART2 in facilitating the transfer of ER-derived fatty acids to chloroplasts for glycolipid synthesis (Hirashima et al., 2021). Plastids and the ER are the main sites of *de novo* fatty acid biosynthesis and LPC production during phosphatidylcholine (PC) metabolism and remodeling. In contrast, the Golgi apparatus functions as a major center for lipid modification, membrane trafficking, and plasma membrane biogenesis (Agliarulo and Parashuraman, 2022; Karki et al., 2019; Ohlrogge et al., 1979). Collectively, these observations suggest that plant minimal START domain proteins may participate in localized lipid exchange between membrane compartments.

In conclusion, this study establishes that plant minimal START proteins have all the characteristics of bona fide lipid-binding proteins that recognize amphipathic lipid molecules through a conserved bipartite mechanism. Ligand binding is coupled to a dynamic β_5_-β_6_ loop-mediated lid mechanism that likely maintains a protected hydrophobic environment during lipid exchange, a feature shared with several box-type lipid transport proteins, although direct lipid-transfer activity has yet to be experimentally demonstrated. The specific interactions of these proteins with biologically important molecules, such as myristic acid and lysophosphatidylcholines, highlight their potential roles in membrane remodeling, lipid signaling, stress responses, and protein N-myristoylation–dependent pathways. Although myristic acid and LPC were identified as *in vitro* binding ligands, the endogenous physiological ligand(s) remain unknown, and the ligand specificity observed under simplified *in vitro* conditions may not fully reflect the repertoire of ligands recognized in the native environment. Even so, these findings establish that minimal START proteins possess intrinsic lipid-binding activity and should not be overlooked simply because they lack additional regulatory domains, though further *in planta* studies and lipid transfer assays using reconstituted lipid vesicles will be important to establish their precise biological functions. Beyond their biological significance, the substantial diversity in cavity architecture, together with the ability to accommodate structurally distinct lipid-like drug molecules without disrupting the overall fold, suggests that minimal START domains may serve as tuneable scaffolds for engineering selective lipid-binding systems or membrane-targeted delivery platforms.

## Acknowledgments

We thank the Central Instrumentation Facility at BRIC–National Institute of Plant Genome Research and the University of Delhi South Campus for providing access to instrumentation facilities. We are particularly grateful to Dr. Manisha Goel, Department of Biophysics, University of Delhi South Campus, for access to the fluorescence spectroscopy facility; Dr. Dinakar M. Salunke, ICGEB, New Delhi, for access to BLI equipment; and Dr. Pinky Aggarwal, BRIC–NIPGR, for confocal data analysis. We also acknowledge the Director, BRIC–National Institute of Plant Genome Research, for continued support.

The authors used artificial intelligence (AI)-assisted tools, specifically ChatGPT and Grammarly, exclusively to improve the clarity, grammar, and readability of the manuscript. These tools were not used to generate scientific content, analyze data, interpret data, or formulate conclusions. All AI-assisted edits were thoroughly reviewed and validated by the authors to ensure scientific accuracy.

## Competing interests

None declared.

## Author contributions

G.Y. and V. G. conceptualization; K.K., S.K.M., M.P., G.T., G.Y., and V.G. methodology; K.K., G.Y., and V.G. validation; K.K., G.Y., and V.G. formal analysis; K.K., G.Y., and V.G. investigation; G.Y. and V. G. resources; K.K. and V. G. writing–original draft; K.K., G.Y., and V. G. writing–review & editing; G.Y. and V. G. supervision; G.Y. and V. G. funding acquisition.

## Data Availability

All data supporting the findings of this study are available in the main text and Supporting Information. **Supporting Information 1** contains SI Figures 1–14, SI Tables 1–7, and the Supplementary Materials and Methods. **Supporting Information 2** contains the sequences of all 562 plant minimal START proteins analyzed in this study. **Supporting Information 3** provides a high-resolution version of the phylogenetic tree shown in Figure 1A. **Supporting Information 4** contains the input structures, trajectory files, and analysis outputs for the MD and metadynamics simulations. Supporting Information 4 is available on Zenodo at https://doi.org/10.5281/zenodo.22246737.

## Funding

This work was supported by the BRIC-National Institute of Plant Genome Research core grant (to V. G. and G.Y).

## Supporting Information

1. Supporting Information 1: SI Figures, SI Tables, Supplementary material and methods
2. Supporting Information 2: Sequences of all 562 minimal START proteins from Plants
3. Supporting Information 3: High-resolution Family Tree
4. Supporting Information 4: MD and metadynamics data (available on Zenodo at https://doi.org/10.5281/zenodo.22246737)

## Notes

### Competing Interest Statement

The authors have declared no competing interest.

### Summary of Updates

new experiments are added: MD simulations and BLI experiments

## References

Agliarulo, I., and Parashuraman, S. (2022). Golgi Apparatus Regulates Plasma Membrane Composition and Function. Cells 11.

Ahn, V.E., Faull, K.F., Whitelegge, J.P., Fluharty, A.L., and Prive, G.G. (2003). Crystal structure of saposin B reveals a dimeric shell for lipid binding. Proc Natl Acad Sci U S A 100, 38–43.

Alpy, F., and Tomasetto, C. (2005). Give lipids a START: the StAR-related lipid transfer (START) domain in mammals. J Cell Sci 118, 2791–2801.

Armon, A., Graur, D., and Ben-Tal, N. (2001). ConSurf: an algorithmic tool for the identification of functional regions in proteins by surface mapping of phylogenetic information. J Mol Biol 307, 447–463.

Banks, J.L., Beard, H.S., Cao, Y., Cho, A.E., Damm, W., Farid, R., Felts, A.K., Halgren, T.A., Mainz, D.T., Maple, J.R., et al. (2005). Integrated Modeling Program, Applied Chemical Theory (IMPACT). J Comput Chem 26, 1752–1780.

Blom, T., Somerharju, P., and Ikonen, E. (2011). Synthesis and biosynthetic trafficking of membrane lipids. Cold Spring Harb Perspect Biol 3, a004713.

Caron, K.M., Soo, S.C., Wetsel, W.C., Stocco, D.M., Clark, B.J., and Parker, K.L. (1997). Targeted disruption of the mouse gene encoding steroidogenic acute regulatory protein provides insights into congenital lipoid adrenal hyperplasia. Proc Natl Acad Sci U S A 94, 11540–11545.

Clark, B.J. (2020). The START-domain proteins in intracellular lipid transport and beyond. Mol Cell Endocrinol 504, 110704.

Clark, B.J., Wells, J., King, S.R., and Stocco, D.M. (1994). The purification, cloning, and expression of a novel luteinizing hormone-induced mitochondrial protein in MA-10 mouse Leydig tumor cells. Characterization of the steroidogenic acute regulatory protein (StAR). J Biol Chem 269, 28314–28322.

Dallakyan, S., and Olson, A.J. (2015). Small-molecule library screening by docking with PyRx. Methods Mol Biol 1263, 243–250.

Druzak, S.A., Tardelli, M., Mays, S.G., El Bejjani, M., Mo, X., Maner-Smith, K.M., Bowen, T., Cato, M.L., Tillman, M.C., Sugiyama, A., et al. (2023). Ligand dependent interaction between PC-TP and PPARdelta mitigates diet-induced hepatic steatosis in male mice. Nat Commun 14, 2748.

Edgar, R.C. (2022). Muscle5: High-accuracy alignment ensembles enable unbiased assessments of sequence homology and phylogeny. Nat Commun 13, 6968.

Epstein, L.F., Alberta, J.A., Pon, L.A., and Orme-Johnson, N.R. (1989). Subcellular localization of a protein produced in adrenal cortex cells in response to ACTH. Endocr Res 15, 117–127.

Fahy, E., Cotter, D., Sud, M., and Subramaniam, S. (2011). Lipid classification, structures and tools. Biochim Biophys Acta 1811, 637–647.

Floris, A., Luo, J., Frank, J., Zhou, J., Orru, S., Biancolella, M., Pucci, S., Orlandi, A., Campagna, P., Balzano, A., et al. (2019). Star-related lipid transfer protein 10 (STARD10): a novel key player in alcohol-induced breast cancer progression. J Exp Clin Cancer Res 38, 4.

Frey, Y., Lungu, C., and Olayioye, M.A. (2025). Regulation and functions of the DLC family of RhoGAP proteins: Implications for development and cancer. Cell Signal 125, 111505.

Glaser, F., Pupko, T., Paz, I., Bell, R.E., Bechor-Shental, D., Martz, E., and Ben-Tal, N. (2003). ConSurf: identification of functional regions in proteins by surface-mapping of phylogenetic information. Bioinformatics 19, 163–164.

Glatz, J.F. (2015). Lipids and lipid binding proteins: a perfect match. Prostaglandins Leukot Essent Fatty Acids 93, 45–49.

Goodstein, D.M., Shu, S., Howson, R., Neupane, R., Hayes, R.D., Fazo, J., Mitros, T., Dirks, W., Hellsten, U., Putnam, N., et al. (2012). Phytozome: a comparative platform for green plant genomics. Nucleic Acids Res 40, D1178–1186.

Han, S., and Cohen, D.E. (2012). Functional characterization of thioesterase superfamily member 1/Acyl-CoA thioesterase 11: implications for metabolic regulation. J Lipid Res 53, 2620–2631.

Hanada, K. (2004). [The molecular machinery CERT for intracellular trafficking of ceramide]. Seikagaku 76, 562–570.

Hanada, K., Kumagai, K., Tomishige, N., and Kawano, M. (2007). CERT and intracellular trafficking of ceramide. Biochim Biophys Acta 1771, 644–653.

Hirashima, T., Jimbo, H., Kobayashi, K., and Wada, H. (2021). A START domain-containing protein is involved in the incorporation of ER-derived fatty acids into chloroplast glycolipids in Marchantia polymorpha. Biochem Biophys Res Commun 534, 436–441.

Horibata, Y., Ando, H., Satou, M., Shimizu, H., Mitsuhashi, S., Shimizu, Y., Itoh, M., and Sugimoto, H. (2017). Identification of the N-terminal transmembrane domain of StarD7 and its importance for mitochondrial outer membrane localization and phosphatidylcholine transfer. Sci Rep 7, 8793.

Huang, Y., Niu, B., Gao, Y., Fu, L., and Li, W. (2010). CD-HIT Suite: a web server for clustering and comparing biological sequences. Bioinformatics 26, 680–682.

Iaea, D.B., Dikiy, I., Kiburu, I., Eliezer, D., and Maxfield, F.R. (2015). STARD4 Membrane Interactions and Sterol Binding. Biochemistry 54, 4623–4636.

Jumper, J., Evans, R., Pritzel, A., Green, T., Figurnov, M., Ronneberger, O., Tunyasuvunakool, K., Bates, R., Zidek, A., Potapenko, A., et al. (2021). Highly accurate protein structure prediction with AlphaFold. Nature 596, 583–589.

Kanno, K., Wu, M.K., Agate, D.S., Fanelli, B.J., Wagle, N., Scapa, E.F., Ukomadu, C., and Cohen, D.E. (2007). Interacting proteins dictate function of the minimal START domain phosphatidylcholine transfer protein/StarD2. J Biol Chem 282, 30728–30736.

Karki, N., Johnson, B.S., and Bates, P.D. (2019). Metabolically Distinct Pools of Phosphatidylcholine Are Involved in Trafficking of Fatty Acids out of and into the Chloroplast for Membrane Production. Plant Cell 31, 2768–2788.

Kim, H.U. (2020). Lipid Metabolism in Plants. Plants (Basel) 9.

Kozlov, A.M., Darriba, D., Flouri, T., Morel, B., and Stamatakis, A. (2019). RAxML-NG: a fast, scalable and user-friendly tool for maximum likelihood phylogenetic inference. Bioinformatics 35, 4453–4455.

Kudo, N., Kumagai, K., Tomishige, N., Yamaji, T., Wakatsuki, S., Nishijima, M., Hanada, K., and Kato, R. (2008). Structural basis for specific lipid recognition by CERT responsible for nonvesicular trafficking of ceramide. Proc Natl Acad Sci U S A 105, 488–493.

Lenoir, M., Kufareva, I., Abagyan, R., and Overduin, M. (2015). Membrane and Protein Interactions of the Pleckstrin Homology Domain Superfamily. Membranes (Basel) 5, 646–663.

Letourneau, D., Bedard, M., Cabana, J., Lefebvre, A., LeHoux, J.G., and Lavigne, P. (2016). STARD6 on steroids: solution structure, multiple timescale backbone dynamics and ligand binding mechanism. Sci Rep 6, 28486.

Letourneau, D., Lorin, A., Lefebvre, A., Frappier, V., Gaudreault, F., Najmanovich, R., Lavigne, P., and LeHoux, J.G. (2012). StAR-related lipid transfer domain protein 5 binds primary bile acids. J Lipid Res 53, 2677–2689.

Li, Y., Yang, Z., Zhang, Y., Guo, J., Liu, L., Wang, C., Wang, B., and Han, G. (2022). The roles of HD-ZIP proteins in plant abiotic stress tolerance. Front Plant Sci 13, 1027071.

Lin, D., Sugawara, T., Strauss, J.F., 3rd, Clark, B.J., Stocco, D.M., Saenger, P., Rogol, A., and Miller, W.L. (1995). Role of steroidogenic acute regulatory protein in adrenal and gonadal steroidogenesis. Science 267, 1828–1831.

Lu, C., Wu, C., Ghoreishi, D., Chen, W., Wang, L., Damm, W., Ross, G.A., Dahlgren, M.K., Russell, E., Von Bargen, C.D., et al. (2021). OPLS4: Improving Force Field Accuracy on Challenging Regimes of Chemical Space. J Chem Theory Comput 17, 4291–4300.

Luz, J.G., Antonysamy, S., Kuklish, S.L., Condon, B., Lee, M.R., Allison, D., Yu, X.P., Chandrasekhar, S., Backer, R., Zhang, A., et al. (2015). Crystal Structures of mPGES-1 Inhibitor Complexes Form a Basis for the Rational Design of Potent Analgesic and Anti-Inflammatory Therapeutics. J Med Chem 58, 4727–4737.

Mahtha, S.K., Kumari, K., Gaur, V., and Yadav, G. (2023). Cavity architecture based modulation of ligand binding tunnels in plant START domains. Comput Struct Biotechnol J 21, 3946–3963.

Mahtha, S.K., Purama, R.K., and Yadav, G. (2021). StAR-Related Lipid Transfer (START) Domains Across the Rice Pangenome Reveal How Ontogeny Recapitulated Selection Pressures During Rice Domestication. Front Genet 12, 737194.

Martin, K., Kopperud, K., Chakrabarty, R., Banerjee, R., Brooks, R., and Goodin, M.M. (2009). Transient expression in Nicotiana benthamiana fluorescent marker lines provides enhanced definition of protein localization, movement and interactions in planta. Plant J 59, 150–162.

Missaoui, K., Gonzalez-Klein, Z., Pazos-Castro, D., Hernandez-Ramirez, G., Garrido-Arandia, M., Brini, F., Diaz-Perales, A., and Tome-Amat, J. (2022). Plant non-specific lipid transfer proteins: An overview. Plant Physiol Biochem 171, 115–127.

Moqadam, M., Gartan, P., Talandashti, R., Chiapparino, A., Titeca, K., Gavin, A.C., and Reuter, N. (2024). A Membrane-Assisted Mechanism for the Release of Ceramide from the CERT START Domain. J Phys Chem B 128, 6338–6351.

Mukherjee, T., Subedi, B., Khosla, A., Begler, E.M., Stephens, P.M., Warner, A.L., Lerma-Reyes, R., Thompson, K.A., Gunewardena, S., and Schrick, K. (2022). The START domain mediates Arabidopsis GLABRA2 dimerization and turnover independently of homeodomain DNA binding. Plant Physiol 190, 2315–2334.

Murcia, M., Faraldo-Gomez, J.D., Maxfield, F.R., and Roux, B. (2006). Modeling the structure of the StART domains of MLN64 and StAR proteins in complex with cholesterol. J Lipid Res 47, 2614–2630.

Nagata, K., and Abe, M. (2021). The lipid-binding START domain regulates the dimerization of ATML1 via modulating the ZIP motif activity in Arabidopsis thaliana. Dev Growth Differ 63, 448–454.

Nelson, B.K., Cai, X., and Nebenfuhr, A. (2007). A multicolored set of in vivo organelle markers for co-localization studies in Arabidopsis and other plants. Plant J 51, 1126–1136.

Nicholls, H.T., Hornick, J.L., and Cohen, D.E. (2017). Phosphatidylcholine transfer protein/StarD2 promotes microvesicular steatosis and liver injury in murine experimental steatohepatitis. Am J Physiol Gastrointest Liver Physiol 313, G50–G61.

Ohlrogge, J.B., Kuhn, D.N., and Stumpf, P.K. (1979). Subcellular localization of acyl carrier protein in leaf protoplasts of Spinacia oleracea. Proc Natl Acad Sci U S A 76, 1194–1198.

Ponting, C.P., and Aravind, L. (1999). START: a lipid-binding domain in StAR, HD-ZIP and signalling proteins. Trends Biochem Sci 24, 130–132.

Prinz, W.A. (2010). Lipid trafficking sans vesicles: where, why, how? Cell 143, 870–874.

Qiu, X., Mistry, A., Ammirati, M.J., Chrunyk, B.A., Clark, R.W., Cong, Y., Culp, J.S., Danley, D.E., Freeman, T.B., Geoghegan, K.F., et al. (2007). Crystal structure of cholesteryl ester transfer protein reveals a long tunnel and four bound lipid molecules. Nat Struct Mol Biol 14, 106–113.

Radauer, C., Lackner, P., and Breiteneder, H. (2008). The Bet v 1 fold: an ancient, versatile scaffold for binding of large, hydrophobic ligands. BMC Evol Biol 8, 286.

Roderick, S.L., Chan, W.W., Agate, D.S., Olsen, L.R., Vetting, M.W., Rajashankar, K.R., and Cohen, D.E. (2002). Structure of human phosphatidylcholine transfer protein in complex with its ligand. Nat Struct Biol 9, 507–511.

Rodriguez-Agudo, D., Ren, S., Hylemon, P.B., Redford, K., Natarajan, R., Del Castillo, A., Gil, G., and Pandak, W.M. (2005). Human StarD5, a cytosolic StAR-related lipid binding protein. J Lipid Res 46, 1615–1623.

Romanowski, M.J., Soccio, R.E., Breslow, J.L., and Burley, S.K. (2002). Crystal structure of the Mus musculus cholesterol-regulated START protein 4 (StarD4) containing a StAR-related lipid transfer domain. Proc Natl Acad Sci U S A 99, 6949–6954.

Ryan, M.M., Temple, B.R., Phillips, S.E., and Bankaitis, V.A. (2007). Conformational dynamics of the major yeast phosphatidylinositol transfer protein sec14p: insight into the mechanisms of phospholipid exchange and diseases of sec14p-like protein deficiencies. Mol Biol Cell 18, 1928–1942.

Schrick, K., Bruno, M., Khosla, A., Cox, P.N., Marlatt, S.A., Roque, R.A., Nguyen, H.C., He, C., Snyder, M.P., Singh, D., et al. (2014). Shared functions of plant and mammalian StAR-related lipid transfer (START) domains in modulating transcription factor activity. BMC Biol 12, 70.

Schrick, K., Nguyen, D., Karlowski, W.M., and Mayer, K.F. (2004). START lipid/sterol-binding domains are amplified in plants and are predominantly associated with homeodomain transcription factors. Genome Biol 5, R41.

Senese, S., Cheung, K., Lo, Y.C., Gholkar, A.A., Xia, X., Wohlschlegel, J.A., and Torres, J.Z. (2015). A unique insertion in STARD9’s motor domain regulates its stability. Mol Biol Cell 26, 440–452.

Sha, B., Phillips, S.E., Bankaitis, V.A., and Luo, M. (1998). Crystal structure of the Saccharomyces cerevisiae phosphatidylinositol-transfer protein. Nature 391, 506–510.

Shivakumar, D., Williams, J., Wu, Y., Damm, W., Shelley, J., and Sherman, W. (2010). Prediction of Absolute Solvation Free Energies using Molecular Dynamics Free Energy Perturbation and the OPLS Force Field. J Chem Theory Comput 6, 1509–1519.

Slonimskiy, Y.B., Egorkin, N.A., Ashikhmin, A.A., Friedrich, T., Maksimov, E.G., and Sluchanko, N.N. (2022). Reconstitution of the functional Carotenoid-Binding Protein from silkworm in E. coli. Int J Biol Macromol 214, 664–671.

Sluchanko, N.N., Slonimskiy, Y.B., Egorkin, N.A., Varfolomeeva, L.A., Faletrov, Y.V., Moysenovich, A.M., Parshina, E.Y., Friedrich, T., Maksimov, E.G., Boyko, K.M., et al. (2022a). Silkworm carotenoprotein as an efficient carotenoid extractor, solubilizer and transporter. Int J Biol Macromol 223, 1381–1393.

Sluchanko, N.N., Slonimskiy, Y.B., Egorkin, N.A., Varfolomeeva, L.A., Kleymenov, S.Y., Minyaev, M.E., Faletrov, Y.V., Moysenovich, A.M., Parshina, E.Y., Friedrich, T., et al. (2022b). Structural basis for the carotenoid binding and transport function of a START domain. Structure 30, 1647–1659 e1644.

Soccio, R.E., Adams, R.M., Romanowski, M.J., Sehayek, E., Burley, S.K., and Breslow, J.L. (2002). The cholesterol-regulated StarD4 gene encodes a StAR-related lipid transfer protein with two closely related homologues, StarD5 and StarD6. Proc Natl Acad Sci U S A 99, 6943–6948.

Soccio, R.E., and Breslow, J.L. (2003). StAR-related lipid transfer (START) proteins: mediators of intracellular lipid metabolism. J Biol Chem 278, 22183–22186.

Swarbrick, C.M., Roman, N., Cowieson, N., Patterson, E.I., Nanson, J., Siponen, M.I., Berglund, H., Lehtio, L., and Forwood, J.K. (2014). Structural basis for regulation of the human acetyl-CoA thioesterase 12 and interactions with the steroidogenic acute regulatory protein-related lipid transfer (START) domain. J Biol Chem 289, 24263–24274.

Tan, L., Tong, J., Chun, C., and Im, Y.J. (2019). Structural analysis of human sterol transfer protein STARD4. Biochem Biophys Res Commun 520, 466–472.

Tang, D., Ade, J., Frye, C.A., and Innes, R.W. (2005). Regulation of plant defense responses in Arabidopsis by EDR2, a PH and START domain-containing protein. Plant J 44, 245–257.

Tian, W., Chen, C., Lei, X., Zhao, J., and Liang, J. (2018). CASTp 3.0: computed atlas of surface topography of proteins. Nucleic Acids Res 46, W363–W367.

Tillman, M.C., Imai, N., Li, Y., Khadka, M., Okafor, C.D., Juneja, P., Adhiyaman, A., Hagen, S.J., Cohen, D.E., and Ortlund, E.A. (2020). Allosteric regulation of thioesterase superfamily member 1 by lipid sensor domain binding fatty acids and lysophosphatidylcholine. Proc Natl Acad Sci U S A 117, 22080–22089.

Tong, J., Manik, M.K., and Im, Y.J. (2018). Structural basis of sterol recognition and nonvesicular transport by lipid transfer proteins anchored at membrane contact sites. Proc Natl Acad Sci U S A 115, E856–E865.

Torres, J.Z., Summers, M.K., Peterson, D., Brauer, M.J., Lee, J., Senese, S., Gholkar, A.A., Lo, Y.C., Lei, X., Jung, K., et al. (2011). The STARD9/Kif16a kinesin associates with mitotic microtubules and regulates spindle pole assembly. Cell 147, 1309–1323.

Tsujishita, Y., and Hurley, J.H. (2000). Structure and lipid transport mechanism of a StAR-related domain. Nat Struct Biol 7, 408–414.

Wilhelm, L.P., Wendling, C., Vedie, B., Kobayashi, T., Chenard, M.P., Tomasetto, C., Drin, G., and Alpy, F. (2017). STARD3 mediates endoplasmic reticulum-to-endosome cholesterol transport at membrane contact sites. EMBO J 36, 1412–1433.

Wojciechowska, I., Mukherjee, T., Knox-Brown, P., Hu, X., Khosla, A., Subedi, B., Ahmad, B., Mathews, G.L., Panagakis, A.A., Thompson, K.A., et al. (2024). Arabidopsis PROTODERMAL FACTOR2 binds lysophosphatidylcholines and transcriptionally regulates phospholipid metabolism. New Phytol 244, 1498–1518.

Wong, L.H., Gatta, A.T., and Levine, T.P. (2019). Lipid transfer proteins: the lipid commute via shuttles, bridges and tubes. Nat Rev Mol Cell Biol 20, 85–101.

Wong, L.H., and Levine, T.P. (2016). Lipid transfer proteins do their thing anchored at membrane contact sites… but what is their thing? Biochem Soc Trans 44, 517–527.

Yu, H., Chen, X., Hong, Y.Y., Wang, Y., Xu, P., Ke, S.D., Liu, H.Y., Zhu, J.K., Oliver, D.J., and Xiang, C.B. (2008). Activated expression of an Arabidopsis HD-START protein confers drought tolerance with improved root system and reduced stomatal density. Plant Cell 20, 1134–1151.

