## Supplemental Information for "Molecular Mechanism of Lipid Recognition and Membrane-Guided Gating in Plant Minimal START Proteins"

[*] **Correspondence:**

### SI FIGURE 1

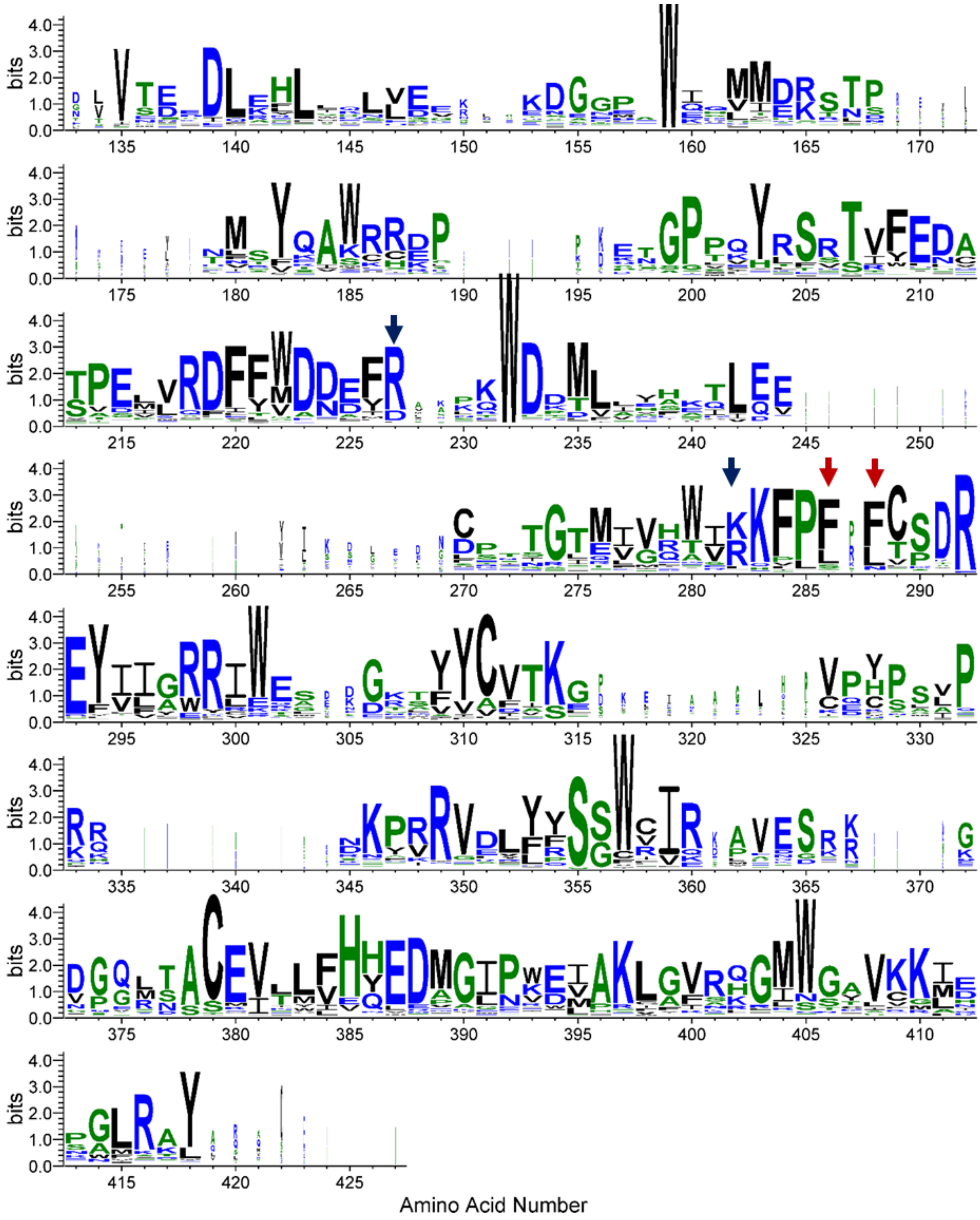


**SI Figure 1: Amino acid conservation landscape of plant minimal START proteins.** The sequences (n = 291) were aligned in Clustal Omega (1), and a conservation profile was visualized using WebLogo 3 (2,3). Blue arrows and red arrows highlight ligand-interacting residues and loop residues, respectively.

### SI FIGURE 2

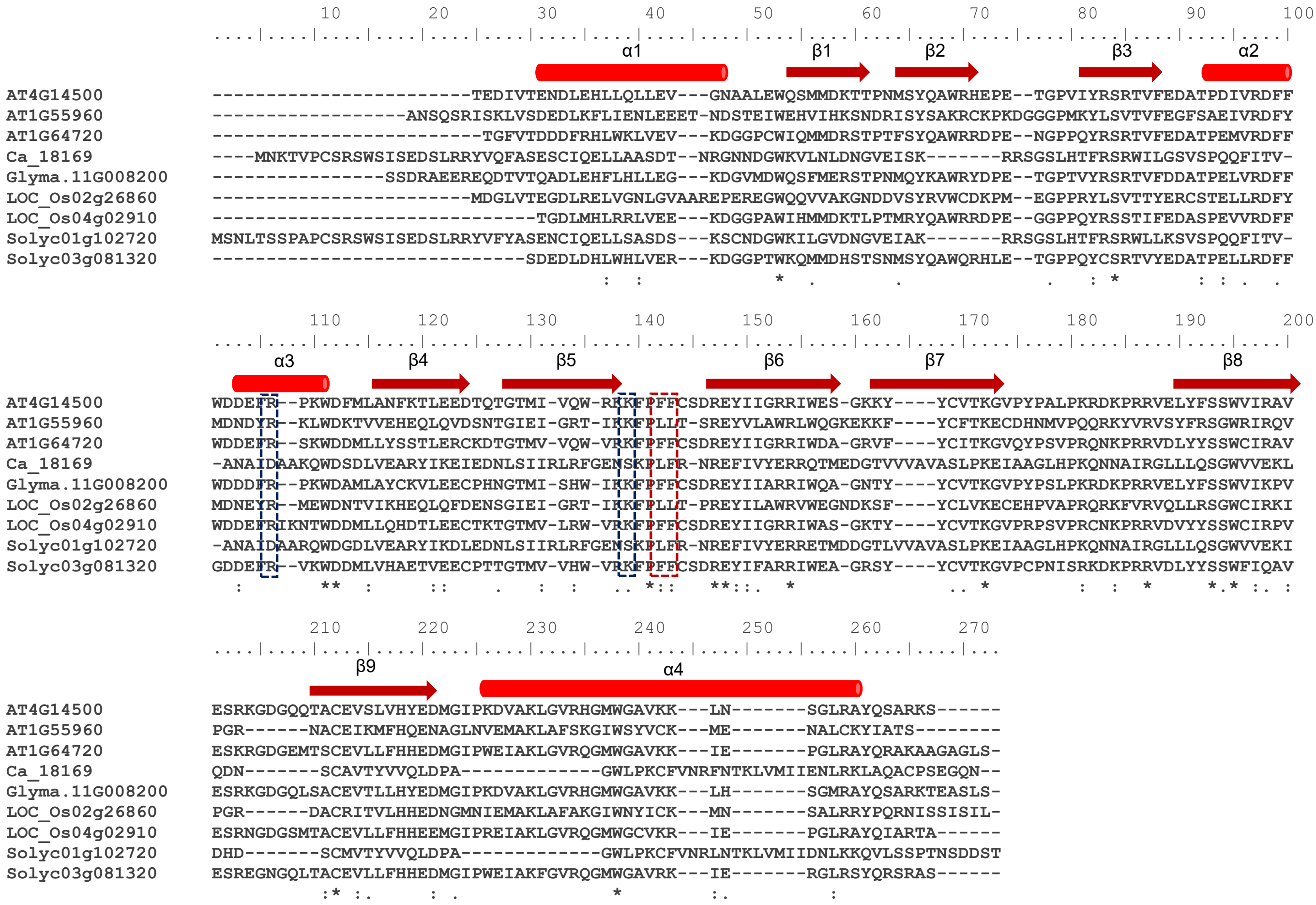


**SI Figure 2: Multiple sequence alignment of representative minimal START proteins with predicted secondary structure.** Amino acid sequences of nine representative minimal START proteins (without TM regions) were aligned, and the corresponding secondary structure elements were superimposed based on predictions from AlphaFold2 models (4). α-helices are indicated by cylinders and β-strands by arrows. The proteins include AT4G14500, AT1G55960, and AT1G64720 from *Arabidopsis thaliana*; Solyc03g081320 and Solyc01g102720 from *Solanum lycopersicum*; LOC_Os02g26860 and LOC_Os04g02910 from *Oryza sativa*; Ca_18169 from *Cicer arietinum*; and Glyma11G008200 from *Glycine max*. All protein accession numbers correspond to entries in the START_FiT database (http://nipgr.ac.in/START_FiT). Blue boxes and red box highlight ligand-interacting residues and loop residues, respectively.

### SI FIGURE 3

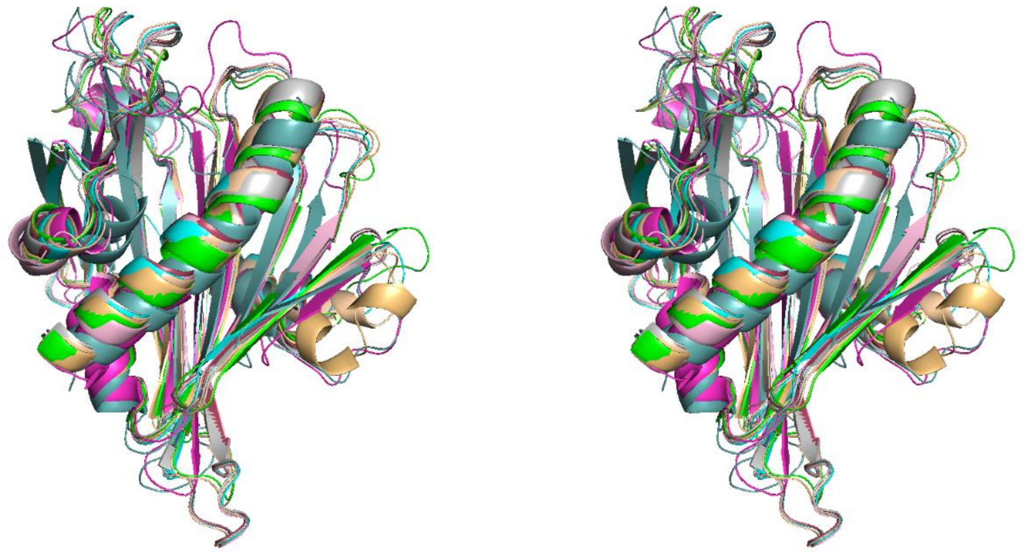

**SI Figure 3: Structural conservation of the minimal START domain across plant species.** Stereo view showing the structural superposition of representative minimal START domain proteins corresponding to Figure 1C, highlighting high structural conservation of the core fold. Proteins from different plant species are color-coded as follows: Arabidopsis thaliana [AT1G55960 (green), AT1G64720 (red), AT4G14500 (yellow)]; Cicer arietinum [Ca_18169 (magenta)]; Glycine max [Glyma.11G008200 (cyan)]; Oryza sativa [LOC_Os02g26860 (orange), LOC_Os04g02910 (pink)]; and Solanum lycopersicum [Solyc01g102720 (teal), Solyc03g081320 (gray)].

### SI FIGURE 4

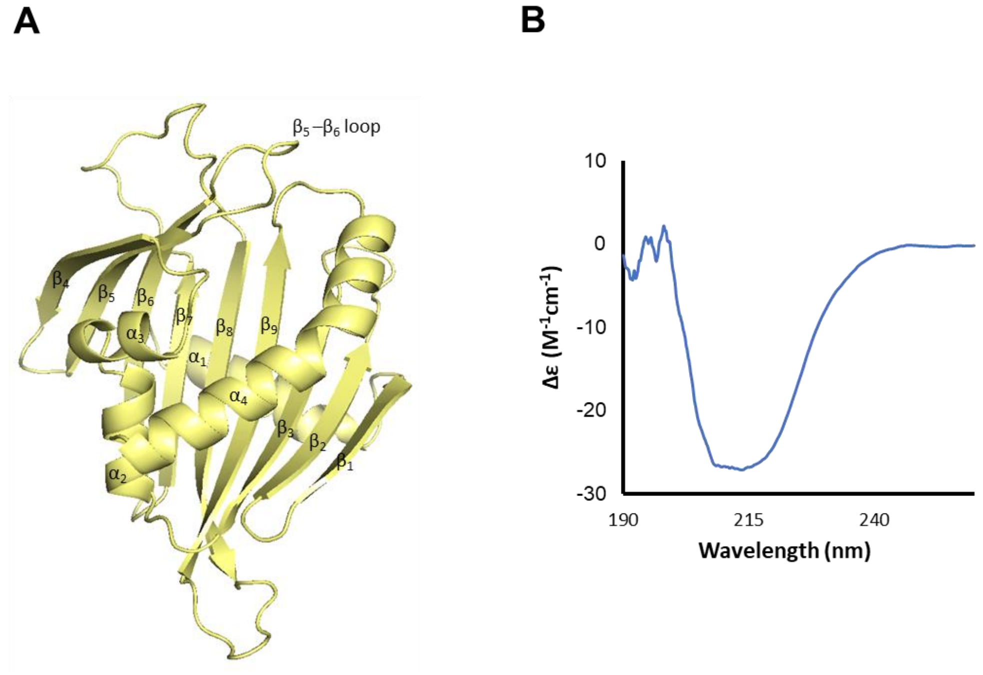


**SI Figure 4: Structural architecture and secondary structure validation of a representative minimal START protein (AT4G14500).**

**(A)** AlphaFold2-predicted structure of the START domain (AT4G14500) showing the characteristic α/β helix-grip fold, comprising a curved antiparallel β-sheet (β1–β9) enclosing a central cavity and flanked by α-helices (α1–α4). The β5–β6 loop, positioned at the cavity entrance, is highlighted.

**(B)** Far-UV circular dichroism spectrum of AT4G14500, displaying characteristic features of a mixed α/β architecture.

### SI FIGURE 5

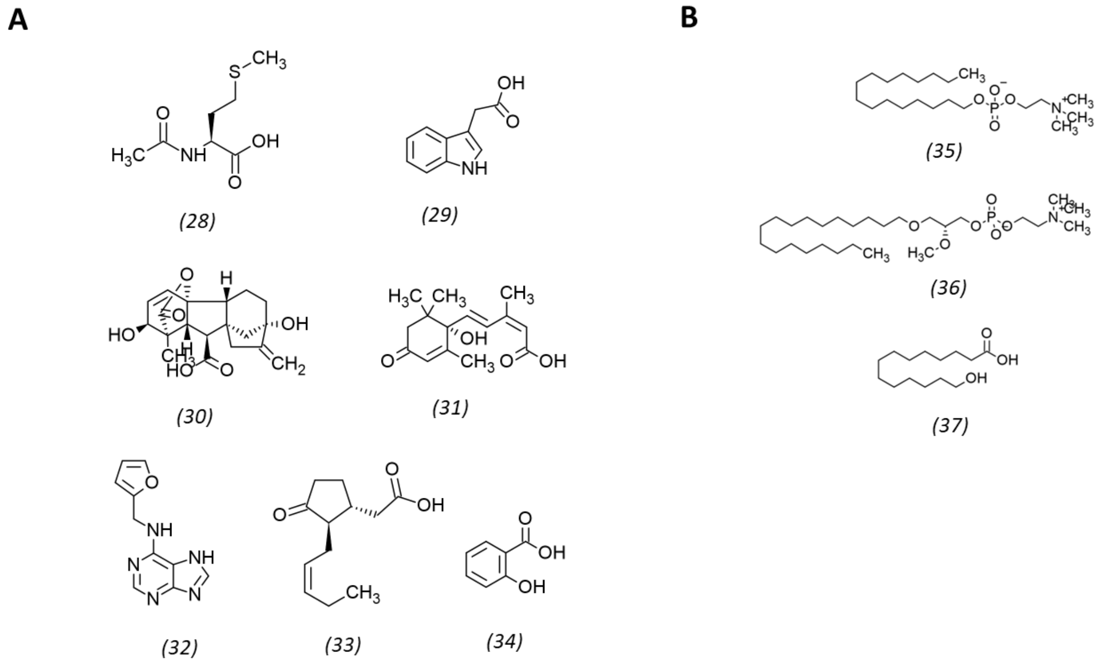


**SI Figure 5: Additional ligands included in the study.**

**(A)** Non-lipid molecules: *(28)* N-acetyl-L-methionine (CID: 448580), *(29)* indole-3-acetic acid (CID: 802), *(30)* gibberellin A3 (CID: 6466), *(31)* abscisic acid (CID: 5280896), *(32)* kinetin (CID: 3830), *(33)* jasmonic acid (CID: 5281166), and *(34)* salicylic acid (CID: 338).

**(B)** Drug-like molecules included as probes: *(35)* miltefosine (CID: 3599), *(36)* edelfosine (CID: 1392), and *(37)* 14-hydroxymyristic acid (CID: 3084276). CID denotes the PubChem identifier.

### SI FIGURE 6

**(A)**

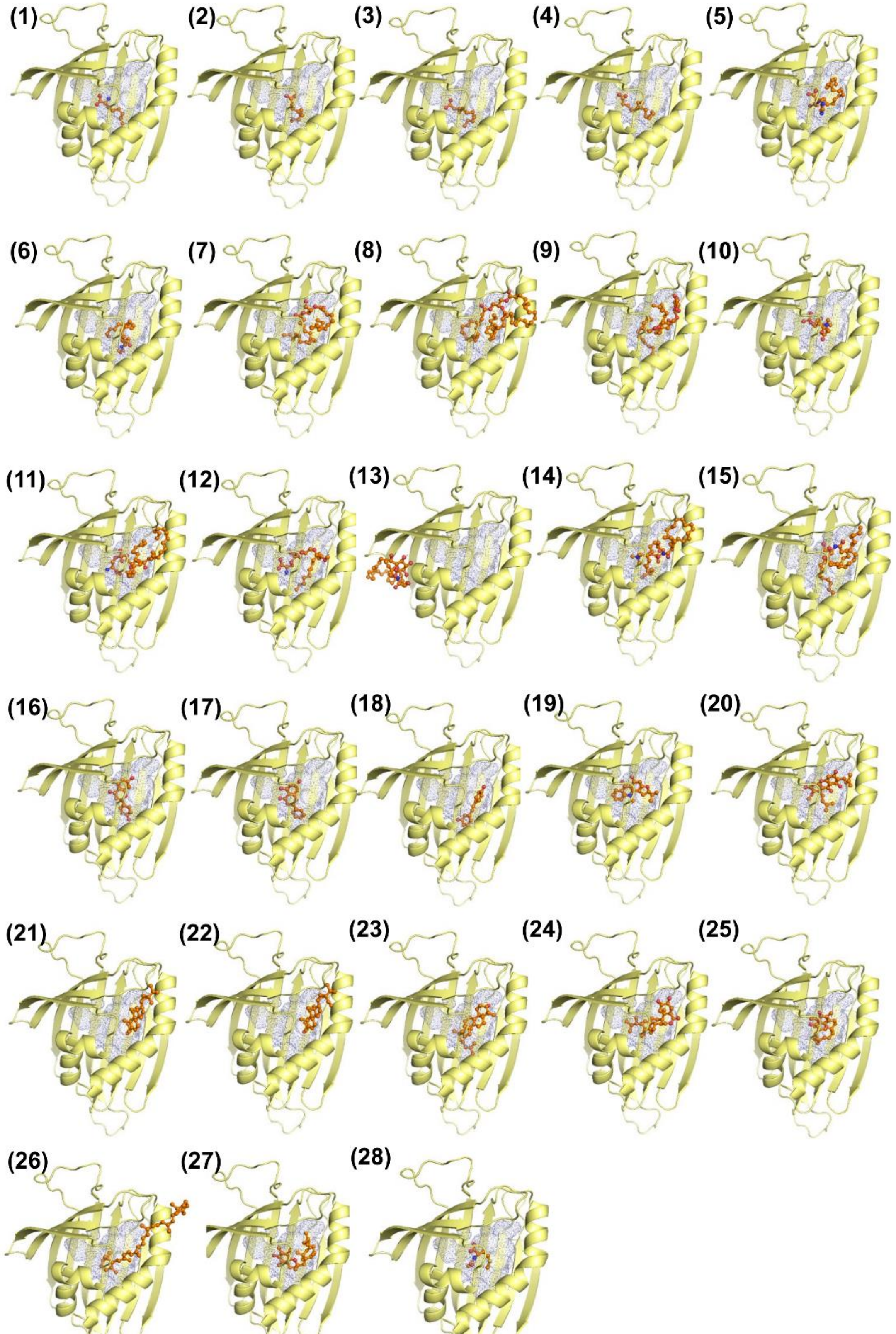

**(B)**

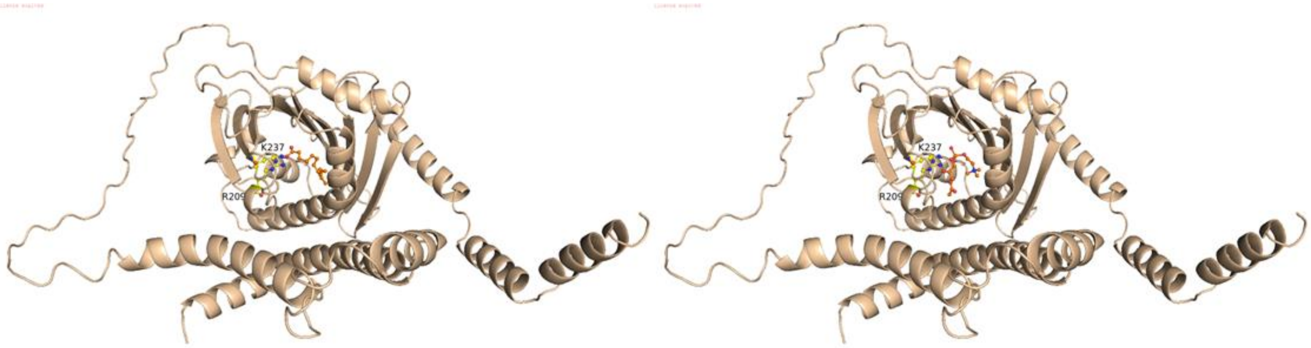


**SI Figure 6: Docking of diverse ligands into the START domain of AT4G14500.**

**(A)** Representative docking poses of 28 ligands within the START domain of AT4G14500. Ligands are numbered as in Figure 2A. The cavity is shown as a blue mesh, and ligands are displayed as sticks (red). Poses illustrate the relative positioning of ligands within or near the predicted binding cavity. While several ligands are accommodated within the cavity, others display partial or surface-associated binding, highlighting variability in ligand accommodation within the START domain.

**(B)** Blind molecular docking of full-length AT4G14500 with myristic acid and LPC. Blind docking poses of full-length AT4G14500 (wheat cartoon) showing the structural accommodation of myristic acid (MA, left panel) and lysophosphatidylcholine (LPC, right panel) within the central START domain binding cavity. Ligands are rendered as orange ball-and-stick models. Key catalytic and contact residues, R209 and K237, are highlighted as yellow sticks and labeled.

#### SI FIGURE 7

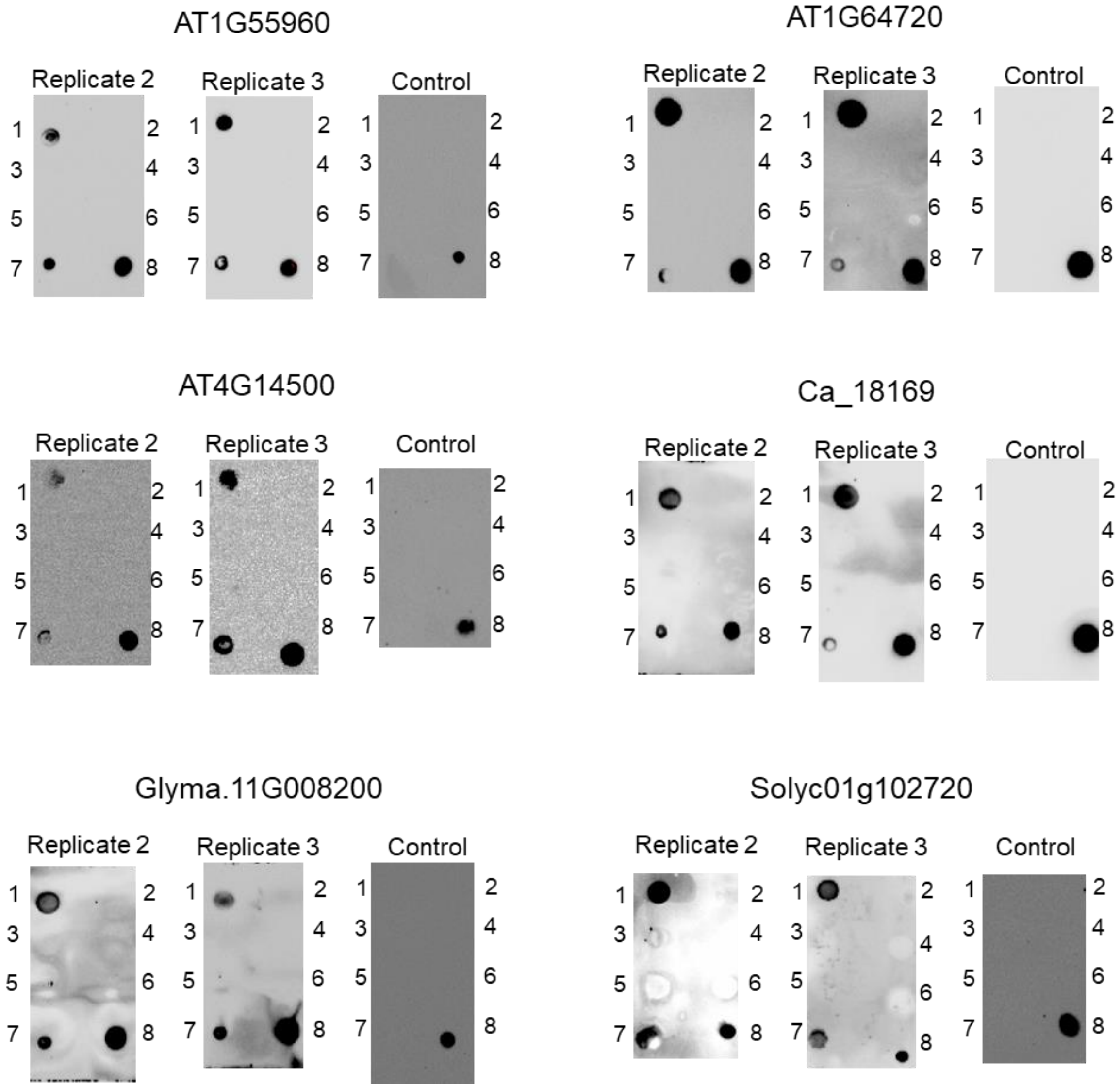


**SI Figure 7: Replicates and controls for protein-lipid overlay assays of ligand binding to minimal START domains.** Independent replicate experiments corresponding to the protein-lipid overlay analyses shown in Figure 3A (lower panel) are presented. Each panel represents a repeat of ligand-binding assays performed under identical conditions. Spots correspond to: (1) myristic acid, (2) dodecanoic acid, (3) decanoic acid, (4) 2-aminooctanoic acid, (5) N-acetyl-L-methionine, (6) sphingomyelin, (7) lysophosphatidylcholine, and (8) positive control (His-tagged protein). In control experiments, protein-lipid overlay assays were performed identically except that buffer was used in place of protein. The reproducibility of binding patterns across replicates supports the specificity of interactions observed for myristic acid and lysophosphatidylcholine.

#### SI FIGURE 8

**A**

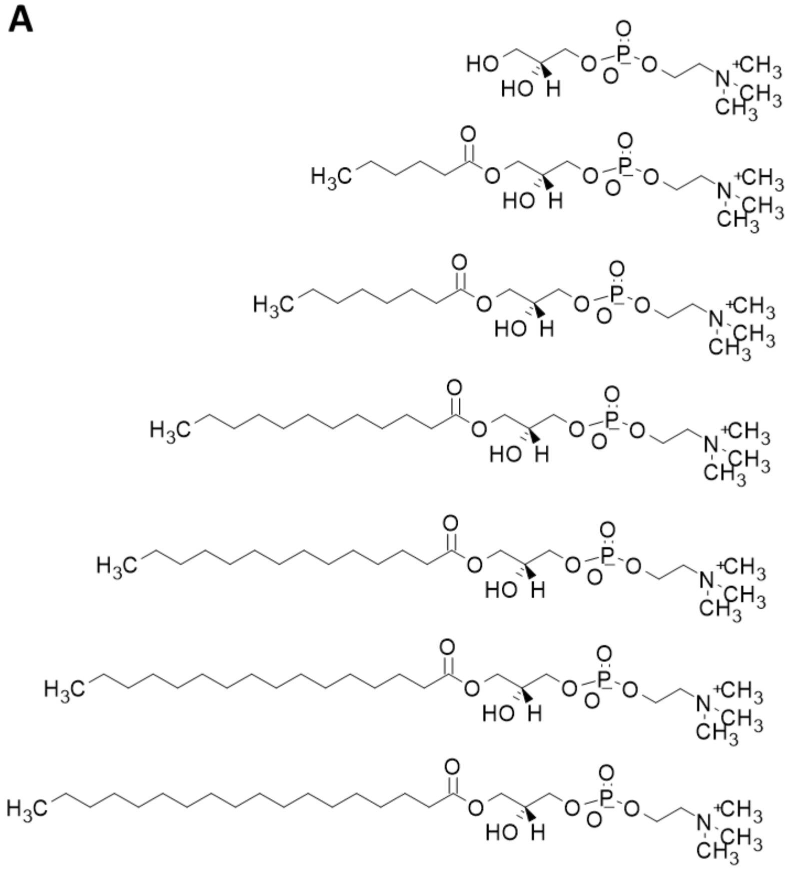

(38)

(39)

(40)

(41)

(42)

(43)

(44)

**B**

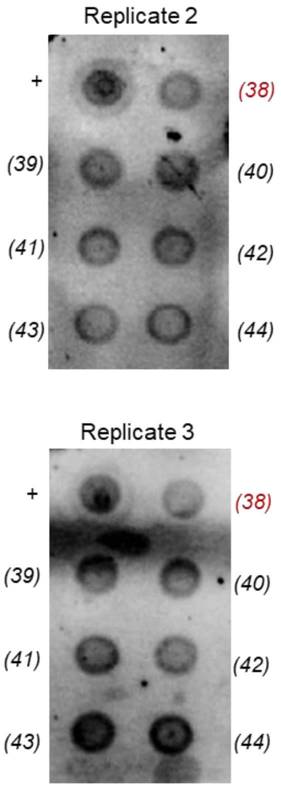


**C**

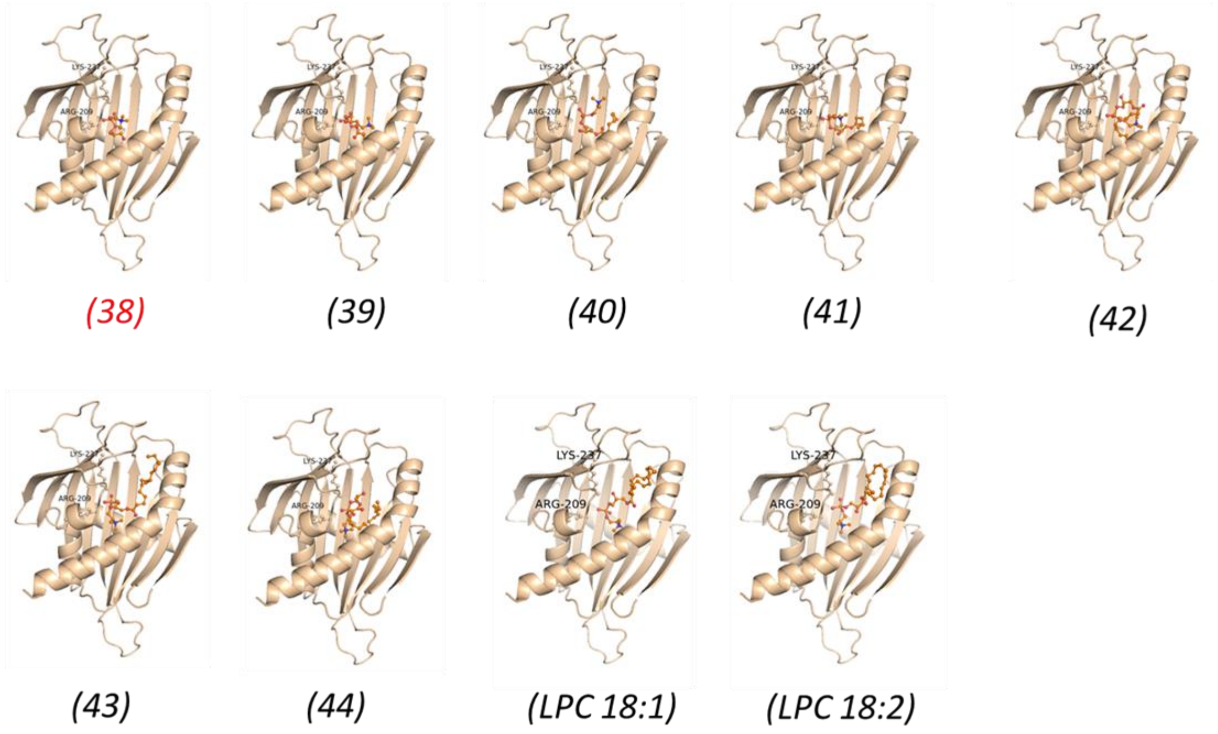

**SI Figure 8: Glycerophosphocholine and lysophosphatidylcholine variants and replicate protein-lipid overlay assays.**

**(A)** Chemical structures of glycerophosphocholine and lysophosphatidylcholine (LPC) variants used in this study, corresponding to ligands 38–44. Ligand 38 represents glycerophosphocholine ($C_8H_{20}NO_6P$), the deacylated phosphocholine headgroup. Ligands 39–44 correspond to LPC molecules with increasing acyl chain lengths: (39) $C_{14}H_{30}NO_7P$, (40) $C_{16}H_{34}NO_7P$, (41) $C_{20}H_{42}NO_7P$, (42) $C_{22}H_{46}NO_7P$, (43) $C_{24}H_{50}NO_7P$, and (44) $C_{26}H_{54}NO_7P$.

**(B)** Independent replicate protein-lipid overlay assays for LPC binding to AT4G14500. Two independent experimental replicates (Replicate 2, top; Replicate 3, bottom) showing protein-lipid overlay assays for AT4G14500. Spots correspond to: (+) positive control (His-tagged protein); (38) glycerophosphocholine ($C_8H_{20}NO_6P$), and (39–44) LPC chain-length variants.

**(C)** Molecular docking of glycerophosphocholine, LPC variants, and unsaturated LPC species into AT4G14500. Blind molecular docking poses showing the structural accommodation of glycerophosphocholine (ligand 38), LPC chain-length variants (ligands 39–44), and unsaturated species: LPC 18:1 ($C_{26}H_{52}NO_7P$, PubChem CID: 16081932) and LPC 18:2 ($C_{26}H_{50}NO_7P$, PubChem CID:11005824) within the hydrophobic cavity of AT4G14500.

### SI FIGURE 9

A

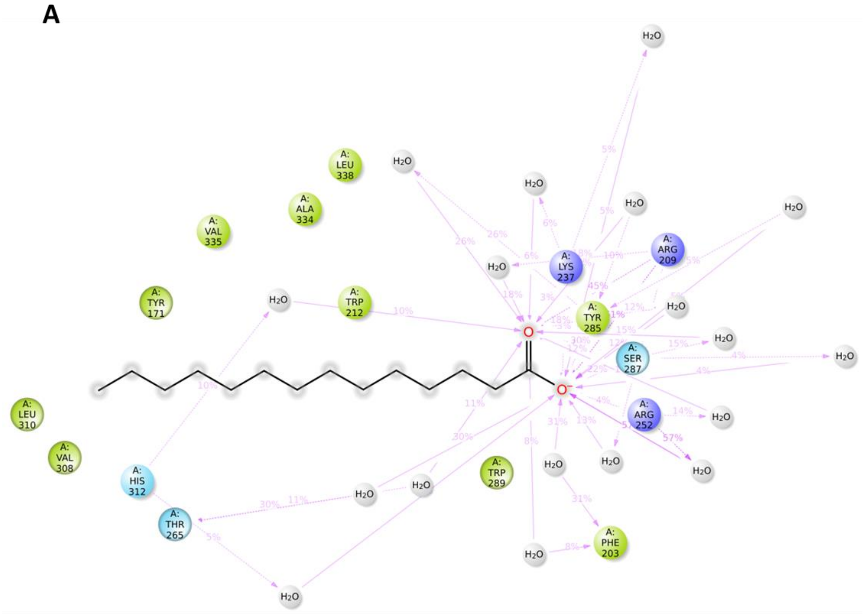


B

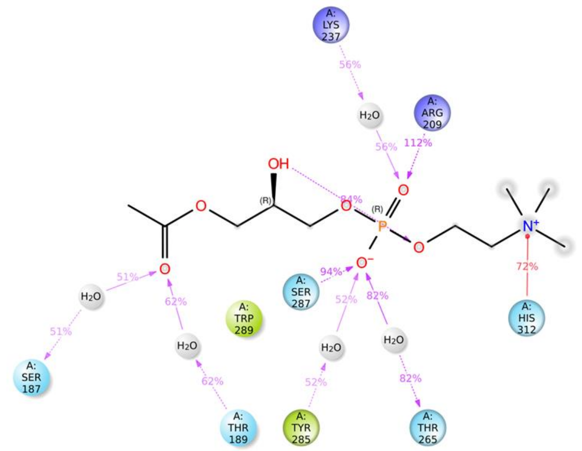


**SI Figure 9: Residue-level interaction networks of AT4G14500 with (A) myristic acid and (B) lysophosphatidylcholine derived from molecular dynamics simulations.** Key residues involved in interactions are shown, with interaction occupancies indicated as percentages.

### SI FIGURE 10

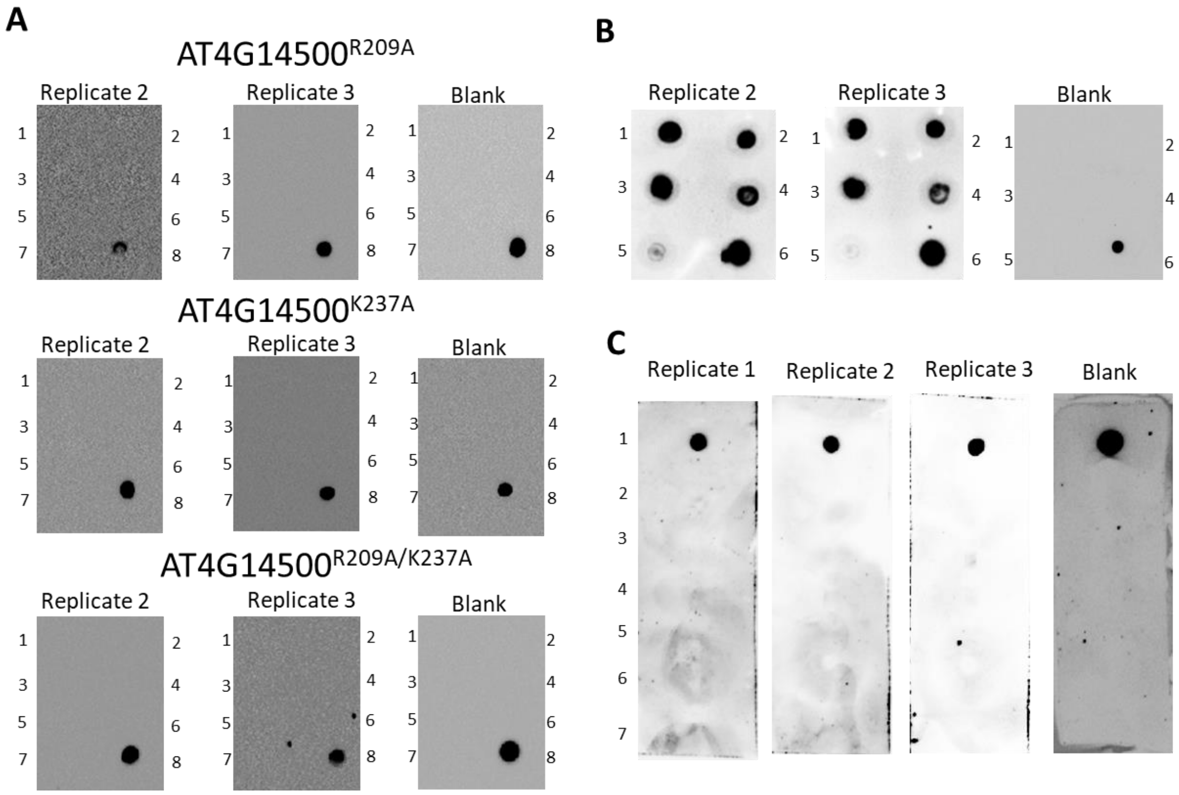


**SI Figure 10: Protein-lipid overlay assays demonstrating ligand binding specificity of AT4G14500.**

**(A)** Protein-lipid overlay assay for AT4G14500 variants. Purified His-tagged AT4G14500$^{R209A}$, AT4G14500$^{K237A}$, and AT4G14500$^{R209A/K237A}$ proteins were incubated with membrane strips spotted with various lipids. Spots correspond to: (1) myristic acid, (2) dodecanoic acid, (3) decanoic acid, (4) 2-aminooctanoic acid, (5) N-acetyl-L-methionine, (6) sphingomyelin, (7) lysophosphatidylcholine, and (8) positive control (His-tagged protein). Independent experiments (Replicates 2 and 3) are shown alongside a Blank membrane control in which protein was omitted. Bound proteins were detected using an anti-His antibody.

**(B)** Protein-lipid overlay assay with selected lipid analogues. Membranes were spotted with representative ligands and incubated with purified His-tagged AT4G14500. Spots correspond to: (1) myristic acid, (2) lysophosphatidylcholine, (3) 14-hydroxymyristic acid, (4) edelfosine, (5) miltefosine, and (6) positive control (His-tagged protein). Independent replicates (Replicates 2 and 3) are shown along with a Blank control.

**(C)** Protein-lipid overlay assay with selected plant hormones. Membranes were spotted with plant hormones and incubated with purified His-tagged AT4G14500. Spots correspond to: (1) positive control (His-tagged protein), (2) indole-3-acetic acid, (3) gibberellin A3, (4) abscisic acid, (5) kinetin, (6) jasmonic acid, and (7) salicylic acid. Replicates (1–3) are shown alongside a Blank membrane control.

### SI FIGURE 11

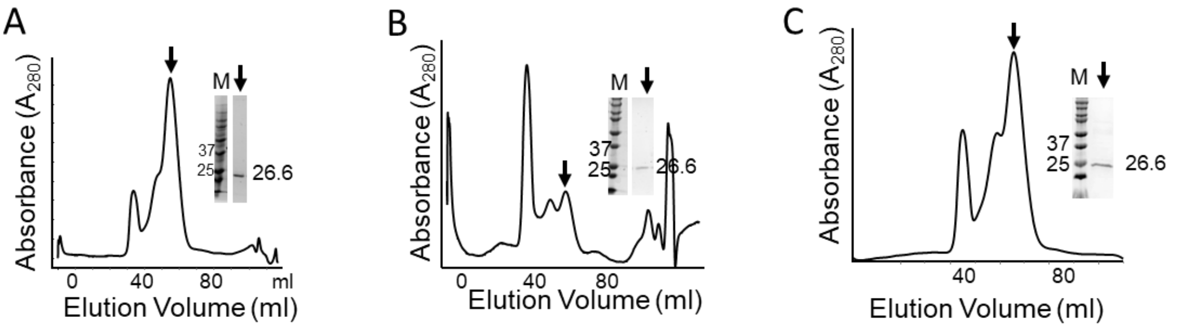


**SI Figure 11: Purification of His-tag free AT4G14500 mutant proteins by size-exclusion chromatography (SEC). (A)** Purification of AT4G14500$^{R209A}$. **(B)** Purification of AT4G14500$^{K237A}$. **(C)** Purification of AT4G14500$^{R209A/K237A}$. The chromatogram shows the elution profile from a Sephacryl S200 gel filtration column. The black arrow indicates the major peak containing the target protein. Inset: 15% SDS–PAGE analysis of the peak fraction; Lane M corresponds to the molecular weight marker (kDa), and the arrow indicates the purified protein band at 26.6 kDa.

### SI FIGURE 12

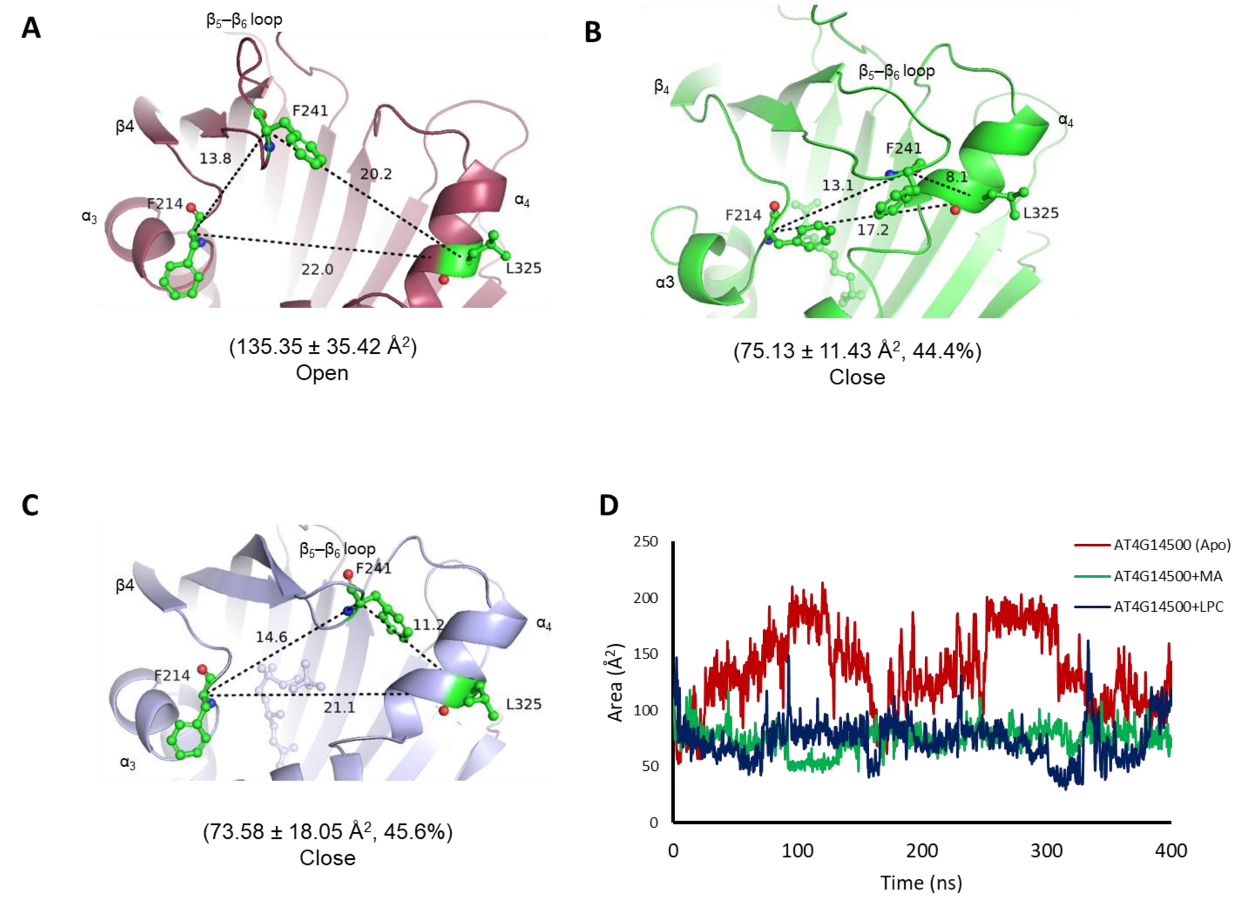


**SI Figure 12: Ligand-induced narrowing of the START domain cavity opening quantified by geometric analysis.** Representative frames from molecular dynamics simulations of AT4G14500 are shown for **(A)** the apo state, **(B)** the MA-bound state, and **(C)** the LPC-bound state. Distances between Cα atoms defining the cavity entrance are indicated in Å. Values in parentheses denote the mean triangle area ± standard deviation calculated over 400 ns MD trajectories, along with the percentage reduction in area for ligand-bound structures relative to the apo state. **(D)** Time-resolved analysis of the cavity entrance area during the 400 ns molecular dynamics simulations. The entrance-gate area, defined by the Cα atoms of residues F214, F241 and L325, is plotted as a function of simulation time for the apo (red), MA-bound (green) and LPC-bound (blue) systems.

### SI FIGURE 13

## A

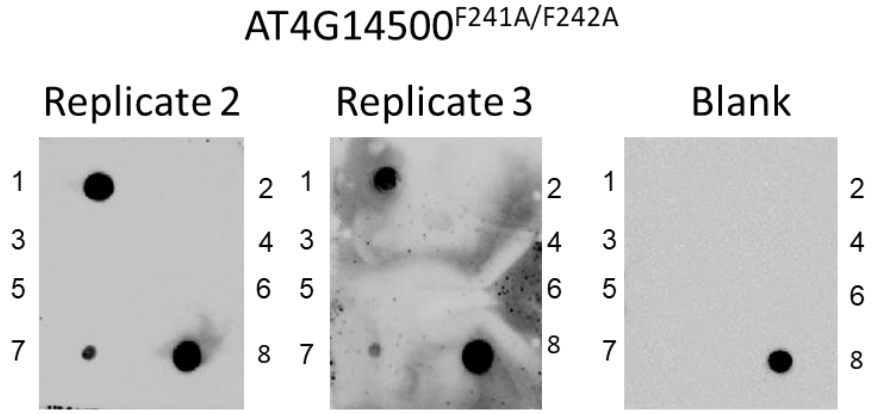


## B

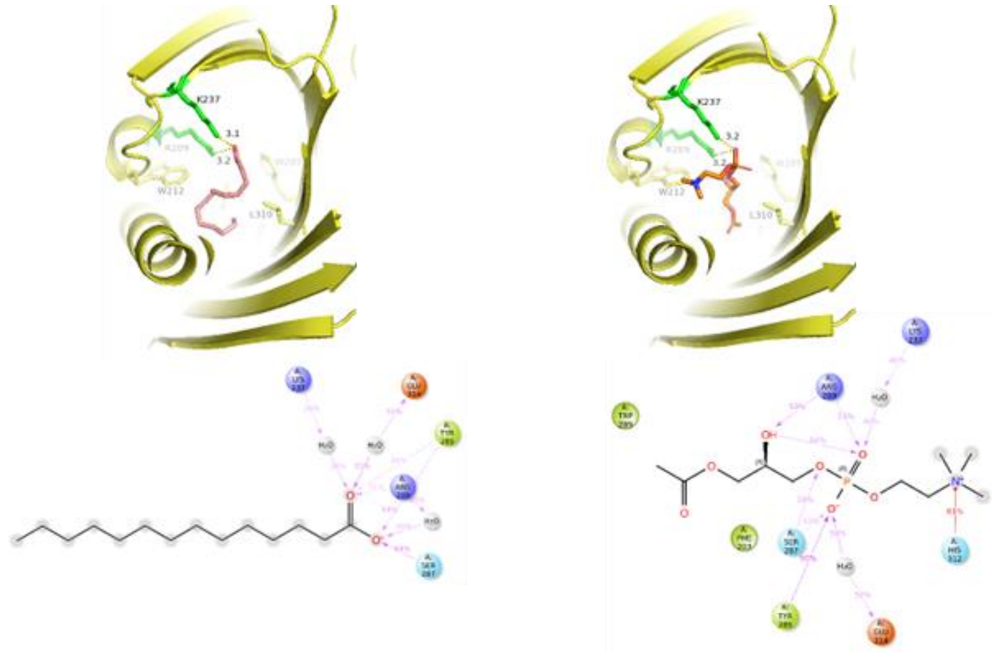


**SI Figure 13. Ligand binding and molecular dynamics analysis of the AT4G14500$^{F241A/F242A}$ mutant. (A)** Protein–lipid overlay assays demonstrating the ligand-binding specificity of AT4G14500$^{F241A/F242A}$. Lipids spotted on the membrane correspond to: (1) myristic acid, (2) dodecanoic acid, (3) decanoic acid, (4) 2-aminooctanoic acid, (5) N-acetyl-L-methionine, (6) sphingomyelin, (7) lysophosphatidylcholine (LPC), and (8) positive control (His-tagged protein). Independent experiments (Replicates 2 and 3) are shown alongside a blank membrane control in which the protein was omitted. Bound protein was detected using an anti-His antibody. **(B)** Representative binding poses and two-dimensional interaction maps obtained from molecular dynamics simulations for 400 ns of the AT4G14500$^{F241A/F242A}$ mutant in complex with myristic acid (left) and lysophosphatidylcholine (right).

### SI Figure 14

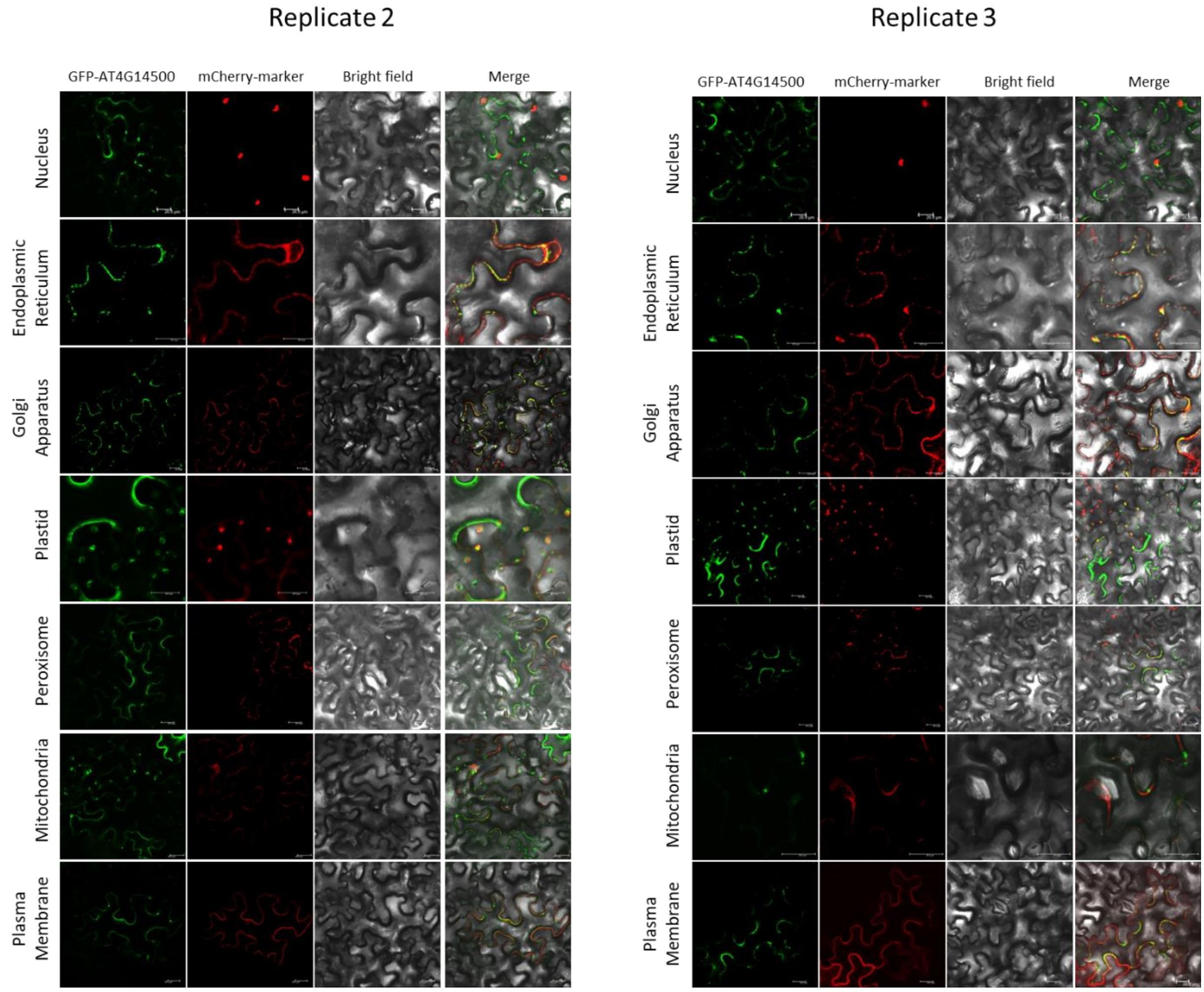


**SI Figure 14: Biological replicates for organelle-specific localization of AT4G14500.** Confocal microscopy images displaying independent biological replicates (Replicate 2, left panel; Replicate 3, right panel) for the subcellular localization of AT4G14500 in *Nicotiana benthamiana* leaves. microscopy Images show expression of GFP-tagged AT4G14500 (green) together with organelle-specific mCherry markers (red). Strong co-localization signals (yellow in merged images) are observed in the endoplasmic reticulum, plastids, Golgi apparatus, and plasma membrane. Scale bars: 20 micrometers.

**SI Table 1: Minimal START domain construct boundaries, AlphaFold2 pLDDT scores, and RMSD values from superimposition with respect to AT1G55960.**

| Protein | Source | Boundary | pLDDT Score | RMSD (Å) |
|---|---|---|---|---|
| AT1G55960 | *Arabidopsis thaliana* | 73-293 | 93.31 | - |
| AT1G64720 | *Arabidopsis thaliana* | 86-308 | 96.08 | 0.857 |
| AT4G14500 | *Arabidopsis thaliana* | 133-351 | 94.61 | 0.832 |
| Ca_18169 | *Cicer arietinum* | 1-235 | 92.86 | 1.778 |
| Glyma11G008200 | *Glycine max* | 122-353 | 94.85 | 0.941 |
| LOC_Os02g26860 | *Oryza sativa* | 103-313 | 93.94 | 0.354 |
| LOC_Os04g02910 | *Oryza sativa* | 98-312 | 95.23 | 0.964 |
| Solyc01g102720 | *Solanum lycopersicum* | 1-240 | 89.46 | 4.380 |
| Solyc03g081320 | *Solanum lycopersicum* | 113-326 | 94.46 | 0.770 |
| AT4G14500 | *Arabidopsis thaliana* | 1-433 | 75.55 | - |

**SI Table 2: Sequences of primers used in site-directed mutagenesis.**

| Primers | Sequence |
|---|---|
| **Mutations in cavity residues** | |
| At4G14500$^{R209A}$R | AAAATCCCATTTCGGAGCAAATTCATCATCCCAAAAGAAATCACGAACA |
| At4G14500$^{R209A}$F | TGTTCGTGATTTCTTTTGGGATGATGAATTTGCTCCGAAATGGGATTTT |
| At4G14500$^{K237A}$R | CTACAGAAAAACGGAAATTTTGCGCGCCACTGCACAATCATGGT |
| At4G14500$^{K237A}$F | ACCATGATTGTGCAGTGGCGCGCAAAATTTCCGTTTTTCTGTAG |
| **Mutations in $\beta_5$-$\beta_6$ loop residues** | |
| At4G14500$^{F241A,F242A}$R | CCGATAATATATTCACGATCACTACAGGCAGCCGGAAATTTTTTGCGCCACTGCACAAT |
| At4G14500$^{F241A,F242A}$F | ATTGTGCAGTGGCGCAAAAAATTTCCGGCTGCCTGTAGTGATCGTGAATATATTATCGG |
| **Primers for cloning in pENTR-D-Topo** | |
| At4G14500$^{cDNA}$F | CACCATGGATGAGACCTATTTTGATC |
| At4G14500$^{cDNA}$R | TCACCTACGAGCCAGTCTCTGG |

**SI Table 3: Expression optimization of minimal START proteins in *E. coli*.**

| Protein | *E. coli* strain | IPTG (mM) | Temperature (°C) |
|---|---|---|---|
| AT1G55960 | RIL | 0.4 | 16 |
| At1G64720 | BL21 | 0.4 | 12 |
| At4G14500 | BL21 | 0.4 | 16 |
| Ca_18169 | BL21 | 1 | 16 |
| Glyma11G008200 | BL21 | 0.4 | 18 |
| LOC_Os02g26860 | | No Expression | |
| LOC_Os04g02910 | | No Expression | |
| Solyc01g102720 | BL21 | 0.4 | 16 |
| Solyc03g081320 | | No Expression | |

**SI Table 4: Ligands used in the study**

| | Ligand | Volume (Å$^3$)[1] | rationale for inclusion[2] | Ref. |
|---|---|---|---|---|
| **FATTY ACYLS** | | | | |
| 1. | 2-aminooctanoic acid | 79.344 | Model amphipathic probe with a fatty-acyl–length hydrophobic moiety (C8) and an amino acid head to investigate generic lipid-binding features of the protein, independent of specific headgroup complexity.<br>Previously suggested as potential ligands | (5-7) |
| 2. | decanoic acid | 88.984 | Certain plant seed oils, particularly those of the genus Cuphea, are known to synthesize and accumulate medium-chain fatty acids, including caprylic (C8:0) and capric (C10:0) acids.<br>Decanoic acid (capric acid) is an intermediate in plant fatty acid synthesis pathways. | (8,9) |
| 3. | dodecanoic acid | 103.157 | Role in Plant defence mechanisms and drought resistance. | (10-12) |
| 4. | myristic acid | 117.284 | Protein myristoylation and photosynthesis repair. | (13,14) |
| 5. | N-octadecanoyl-histidine | 223.061 | Non-native synthetic probe: to assess potential recognition of lipid–amino acid conjugates by START domains.<br>Previously suggested as potential ligands | (7) |
| 6. | α-linolenoyl ethanolamide | 161.13 | Role in development and defence. | (15-17) |
| **GLYCEROLIPIDS** | | | | |
| 7. | Diacylglycerol | 279.082 | Involved in lipid-mediated signalling pathways in plants | (18) |
| 8. | Triacylglycerols | 338.313 | Major storage compound, development, mitigation of biotic and abiotic stresses. | (19) |
| 9. | Monogalactosyldiacylglycerol | 346.597 | Most abundant lipid in photosynthetic plant membranes | (20) |
| **GLYCEROPHOSPHOLIPIDS** | | | | |
| 10. | Lysophosphatidylcholine | 121.571 | Involved in lipid-mediated signalling pathways in plants | (21) |
| 11. | Phosphatidylserine | 278.103 | Cell plate formation | (22) |
| 12. | Glycerophosphoserine | 250.120 | Phosphate scavenging | (23) |
| **SPHINGOLIPIDS** | | | | |
| 13. | Cerebroside | 328.141 | Defence responses in plants | (24) |
| 14. | Sphingomyelin | 266.997 | Non-native xenoligand: to check cross-kingdom molecular mimicry.<br>Previously suggested as potential ligands. | (7,25,26) |
| 15. | Ceramide | 250.604 | programmed cell death in plants | (27) |
| **POLYKETIDES** | | | | |
| 16. | Naringenin | 109.676 | Pathogen resistance, plant growth and development | (28-30) |
| 17. | Pinocembrin | 130.003 | A key bioflavonoid found in plants, particularly in *Peperomia* and *Piper* genera, and *Asteraceae* families | (31) |
| 18. | Resveratrol | 102.393 | Naturally occurring polyketide produced by many plants in response to biotic and abiotic stress. | (32) |

| | | | | |
|---|---|---|---|---|
| 19. | Rutacridone | 233.857 | Main alkaloid in tissue cultures of *Ruta graveolens.* | (33) |
| 20. | Mangostin | 209.979 | One of the active compounds with antimicrobial properties found in *Garcinia* fruits. | (34) |
| | | **STEROLS** | | |
| 21. | Campesterol | 198.552 | A major phytosterol found in cell membranes, crucial for plant growth, development, and stress response. | (35) |
| 22. | β-Sitosterol | 164.365 | Crucial for plant growth, development, and stress response | (35) |
| 23. | Lanosterol | 162.421 | Phytosterols production in *Arabidopsis.* | (36,37) |
| 24. | 24-Epibrassinolide | 211.388 | Promotes growth and resistance to abiotic stress. | (38-40) |
| | | **PRENOLS** | | |
| 25. | 9'-Carboxy-alpha-tocotrienol | 166.521 | Abundant in palm oil, rice bran oil, wheat germ, barley, and various nuts and grains. | (41) |
| 26. | β-carotene | 255.166 | Photoprotection, anti-oxidant, precursor to plant hormones | (42) |
| 27. | α-tocopherol | 197.292 | Protects photosynthetic machinery. | (43) |
| | | **MISCELLANEOUS** | | |
| 28. | N-acetyl-L-methionine | 97.743 | Putative modified amino acid: to assess potential recognition of acetylated amino acid derivatives in plant proteins.<br>Previously suggested as potential ligands. | (7) |
| 29. | Indole-3-acetic acid | 79.255 | Primary auxin regulating plant growth and development | (44) |
| 30. | Gibberellin A3 | 113.467 | A bioactive gibberellin regulating plant growth | (45) |
| 31. | Abscisic Acid | 105.321 | A central hormone involved in plant stress responses | (46) |
| 32. | Kinetin | 98.751 | One of the Cytokinins. | (47) |
| 33. | Jasmonic Acid | 113.319 | A key signalling molecule in wound and defence responses | (48) |
| 34. | Salicylic Acid | 61.422 | Improves plant immunity, growth, and stress tolerance, acting as a signaling molecule for systemic acquired resistance (SAR) against pathogens. | (49) |
| | | **DRUG MOLECULES** | | |
| 35. | Miltefosine | 158.155 | Non-native lysophospholipid analogue, anti-cancer drug, repurposed as antileishmanial drug. | (50) |
| 36. | Edelfosine | 209.700 | Non-native lysophospholipid analogue, anticancer drug | (51) |
| 37. | 14-hydroxymyristic acid | 97.950 | Modified fatty acid analogue | (52,53) |

[1] Molecular volumes were calculated using the Sanjeevini web server for target-directed lead discovery (54).

[2] These descriptions are not exhaustive and should not be interpreted as complete functional annotations in plants.

**SI Table 5: Grid parameters used for blind docking of minimal START proteins.**

| Protein | Grid Center (X, Y, Z) (Å) | Grid Size (X, Y, Z) (Å) |
|---|---|---|
| AT1G55960 | -4.360, 2.548, -2.594 | 44.261, 42.407, 52.831 |
| At1G64720 | 0.750, -2.143, 0.836 | 48.708, 44.824, 45.305 |
| AT4G14500 | 0.925, -1.514, 2.354 | 53.162, 49.415, 53.737 |
| Ca_18169 | -1.332, -1.267, -0.168 | 46.570, 49.561, 42.925 |
| Glyma.11G008200 | 0.258, 0.762, 3.026 | 47.558, 44.440, 49.450 |
| LOC_Os02g26860 | -6.281, 3.578, -3.548 | 49.126, 46.035, 52.226 |
| LOC_Os04g02910 | 0.204, -1.958, -0.009 | 56.527, 39.347, 47.175 |
| Solyc01g102720 | 0.242, -0.183, 0.251 | 41.718, 51.695, 46.593 |
| Soly03g081320 | -6.056, 2.724, -4.547 | 54.535, 45.901, 46.664 |
| AT4G14500$^{FL*}$ | 2.627, 10.141, -0.210 | 71.446, 99.893, 92.430 |

* Only AT4G14500FL with putative transmembrane motifs was used for docking, For rest of the proteins only the START domain was used

**SI Table 6: Structure-guided evaluation of docking poses for ligand binding across representative minimal START proteins.**

**SI Table 6A: Docking pose evaluation for AT1G55960**

| Ligand No. | Cavity Binding[1] | Pose[2] | Binding Energy (kcal/mol | Status[3] | Ligand No. | Cavity Binding[1] | Pose[2] | Binding Energy (kcal/mol) | Status |
|---|---|---|---|---|---|---|---|---|---|
| 1 | Full | 6 | -4.7 | Selected | 15 | Full | PS | - | Discarded |
| 2 | Full | 9 | -4.5 | Selected | 16 | Full | 2 | -7.7 | Selected |
| 3 | Full | 9 | -4.8 | Selected | 17 | Full | 3 | -7.6 | Selected |
| 4 | Full | 3 | -5.1 | Selected | 18 | Full | 4 | -7.1 | Selected |
| 5 | Full | PS | - | Discarded | 19 | Full | 4 | -7.7 | Selected |
| 6 | Full | PS | - | Discarded | 20 | Full | 1 | -7.3 | Selected |
| 7 | Full | - | - | Discarded | 21 | Full | PS | - | Discarded |
| 8 | Full | PS | - | Discarded | 22 | Full | PS | - | Discarded |
| 9 | Full | PS | - | Discarded | 23 | Full | PS | - | Discarded |
| 10 | Full | 3 | -5.5 | Selected | 24 | Full | PS | - | Discarded |
| 11 | Full | PS | - | Discarded | 25 | Full | PS | - | Discarded |
| 12 | Full | PS | - | Discarded | 26 | Partial | - | - | Discarded |
| 13 | Full | PS | - | Discarded | 27 | Full | 1 | -8.1 | Selected |
| 14 | Full | PS | - | Discarded | 28 | Full | 1 | -6.8 | Selected |

**SI Table 6B: Docking pose evaluation for AT1G64720**

| Ligand No. | Cavity Binding[1] | Pose[2] | Binding Energy (kcal/mol) | Status[3] | Ligand No. | Cavity Binding[1] | Pose[2] | Binding Energy (kcal/mol) | Status |
|---|---|---|---|---|---|---|---|---|---|
| 1 | Full | 6 | -4.2 | Selected | 15 | Full | PS | - | Discarded |
| 2 | Full | 1 | -4.6 | Selected | 16 | Full | 1 | -7.5 | Selected |
| 3 | Full | 8 | -4.5 | Selected | 17 | Full | 1 | -7.4 | Selected |
| 4 | Full | 1 | -4.5 | Selected | 18 | Full | 1 | -6.4 | Selected |
| 5 | Full | 3 | -5.4 | Selected | 19 | Full | PS | - | Discarded |
| 6 | Full | 6 | -4.9 | Selected | 20 | Full | PS | - | Discarded |
| 7 | Full | PS | - | Discarded | 21 | Full | PS | - | Discarded |
| 8 | Partial | - | - | Discarded | 22 | Partial | - | - | Discarded |
| 9 | Partial | - | - | Discarded | 23 | Full | PS | - | Discarded |
| 10 | Full | 5 | -5.0 | Selected | 24 | Full | PS | - | Discarded |
| 11 | Partial | - | - | Discarded | 25 | Full | 3 | -7.3 | Selected |
| 12 | Full | PS | - | Discarded | 26 | Partial | - | - | Discarded |
| 13 | Partial | - | - | Discarded | 27 | Full | PS | - | Discarded |
| 14 | Full | PS | - | Discarded | 28 | Full | 1 | -6.5 | Selected |

[1] Full= Ligand binding completely inside the cavity; Partial: Ligand binding partially; Zero: Ligand not binding inside the cavity.
[2] Numbers: Pose number with correct stereochemistry of the ligand; PS: Poor stereochemistry of the ligand in all the poses.
[3] Selected: Ligand binding fully accommodated with in the cavity with appropriate stereochemistry; Not Selected: Ligand binding with partial or zero cavity occupancy and/or distorted stereochemistry

**SI Table 6C: Docking pose evaluation for AT4G14500**

| Ligand No. | Cavity Binding[1] | Pose[2] | Binding Energy (kcal/mol) | Status[3] | Ligand No. | Cavity Binding[1] | Pose[2] | Binding Energy (kcal/mol) | Status |
|---|---|---|---|---|---|---|---|---|---|
| 1 | Full | 3 | -4.1 | Selected | 15 | Partial | - | - | Discarded |
| 2 | Full | 6 | -4.4 | Selected | 16 | Full | 1 | -7.2 | Selected |
| 3 | Full | 1 | -4.9 | Selected | 17 | Full | 1 | -7.6 | Selected |
| 4 | Full | 1 | -4.6 | Selected | 18 | Full | 1 | -6.2 | Selected |
| 5 | Full | PS | - | Discarded | 19 | Full | 1 | -8.3 | Selected |
| 6 | Full | PS | - | Discarded | 20 | Full | 1 | -7.3 | Selected |
| 7 | Partial | - | - | Discarded | 21 | Partial | - | - | Discarded |
| 8 | Partial | - | - | Discarded | 22 | Partial | - | - | Discarded |
| 9 | Partial | - | - | Discarded | 23 | Full | PS | - | Discarded |
| 10 | Full | 5 | -4.9 | Selected | 24 | Full | PS | - | Discarded |
| 11 | Partial | - | - | Discarded | 25 | Full | PS | | Discarded |
| 12 | Full | PS | - | Discarded | 26 | Zero | - | - | Discarded |
| 13 | Zero | - | - | Discarded | 27 | Partial | - | - | Discarded |
| 14 | Partial | - | - | Discarded | 28 | Full | 1 | -5.4 | Selected |

**SI Table 6D: Docking pose evaluation for Ca_18169**

| Ligand No. | Cavity Binding[1] | Pose[2] | Binding Energy (kcal/mol) | Status[3] | Ligand No. | Cavity Binding[1] | Pose[2] | Binding Energy (kcal/mol) | Status |
|---|---|---|---|---|---|---|---|---|---|
| 1 | Full | 3 | 4.8 | Selected | 15 | Partial | - | - | Discarded |
| 2 | Full | 5 | 5.1 | Selected | 16 | Full | 1 | -8.2 | Selected |
| 3 | Full | 3 | 5.6 | Selected | 17 | Full | 1 | -9.1 | Selected |
| 4 | Full | 8 | 5.5 | Selected | 18 | Full | 3 | -7.5 | Selected |
| 5 | Full | PS | - | Discarded | 19 | Full | 1 | -8.5 | Selected |
| 6 | Full | PS | - | Discarded | 20 | Zero | - | - | Discarded |
| 7 | Full | PS | - | Discarded | 21 | Full | PS | - | Discarded |
| 8 | Full | PS | - | Discarded | 22 | Zero | - | - | Discarded |
| 9 | Zero | - | - | Discarded | 23 | Full | PS | - | Discarded |
| 10 | Full | 1 | 6.1 | Selected | 24 | Partial | PS | - | Discarded |
| 11 | Zero | - | - | Discarded | 25 | Partial | - | - | Discarded |
| 12 | Full | PS | - | Discarded | 26 | Partial | - | - | Discarded |
| 13 | Partial | - | - | Discarded | 27 | Partial | - | - | Discarded |
| 14 | Partial | - | - | Discarded | 28 | Full | 1 | -5.8 | Selected |

[1] Full= Ligand binding completely inside the cavity; Partial: Ligand binding partially; Zero: Ligand not binding inside the cavity.
[2] Numbers: Pose number with correct stereochemistry of the ligand; PS: Poor stereochemistry of the ligand in all the poses.
[3] Selected: Ligand binding fully accommodated with in the cavity with appropriate stereochemistry; Not Selected: Ligand binding with partial or zero cavity occupancy and/or distorted stereochemistry

**SI Table 6E: Docking pose evaluation for Glyma.11G008200**

| Ligand No. | Cavity Binding[1] | Pose[2] | Binding Energy (kcal/mol) | Status[3] | Ligand No. | Cavity Binding[1] | Pose[2] | Binding Energy (kcal/mol) | Status |
|---|---|---|---|---|---|---|---|---|---|
| 1 | Full | 2 | -4.4 | Selected | 15 | Full | PS | - | Discarded |
| 2 | Full | 9 | -4.5 | Selected | 16 | Full | 1 | -7.0 | Selected |
| 3 | Full | 5 | -4.5 | Selected | 17 | Full | 1 | -7.1 | Selected |
| 4 | Full | 1 | -5.0 | Selected | 18 | Full | 1 | -6.2 | Selected |
| 5 | Full | PS | - | Discarded | 19 | Full | 1 | -8.1 | Selected |
| 6 | Full | 1 | -6.1 | Selected | 20 | Full | 1 | -7.7 | Selected |
| 7 | Partial | - | - | Discarded | 21 | Full | PS | - | Discarded |
| 8 | Partial | - | - | Discarded | 22 | Full | PS | - | Discarded |
| 9 | Full | PS | - | Discarded | 23 | Full | PS | - | Discarded |
| 10 | Full | 1 | -4.9 | Selected | 24 | Full | PS | - | Discarded |
| 11 | Partial | - | - | Discarded | 25 | Full | 1 | -7.8 | Selected |
| 12 | Full | PS | - | Discarded | 26 | Partial | - | - | Discarded |
| 13 | Full | PS | - | Discarded | 27 | Full | 1 | -6.9 | Selected |
| 14 | Full | PS | - | Discarded | 28 | Full | 1 | -6.0 | Selected |

**SI Table 6F: Docking pose evaluation for LOC_Os02g26860**

| Ligand No. | Cavity Binding[1] | Pose[2] | Binding Energy (kcal/mol) | Status[3] | Ligand No. | Cavity Binding[1] | Pose[2] | Binding Energy (kcal/mol) | Status |
|---|---|---|---|---|---|---|---|---|---|
| 1 | Full | 4 | -4.7 | Selected | 15 | Partial | - | - | Discarded |
| 2 | Full | 6 | -4.6 | Selected | 16 | Full | 1 | -8.0 | Selected |
| 3 | Full | 8 | -4.6 | Selected | 17 | Full | 1 | -7.7 | Selected |
| 4 | Full | 7 | -4.8 | Selected | 18 | Full | 1 | -7.1 | Selected |
| 5 | Full | PS | - | Discarded | 19 | Full | 1 | -8.4 | Selected |
| 6 | Full | 7 | -5.4 | Selected | 20 | Full | 1 | -8.9 | Selected |
| 7 | Full | PS | - | Discarded | 21 | Full | PS | - | Discarded |
| 8 | Full | PS | - | Discarded | 22 | Full | PS | - | Discarded |
| 9 | Full | PS | - | Discarded | 23 | Full | PS | - | Discarded |
| 10 | Full | 3 | -5.0 | Selected | 24 | Full | PS | - | Discarded |
| 11 | Full | PS | - | Discarded | 25 | Full | PS | - | Discarded |
| 12 | Full | PS | - | Discarded | 26 | Partial | - | - | Discarded |
| 13 | Full | PS | - | Discarded | 27 | Full | 8 | -7.2 | Selected |
| 14 | Full | PS | - | Discarded | 28 | Full | 1 | -6.5 | Selected |

[1] Full= Ligand binding completely inside the cavity; Partial: Ligand binding partially; Zero: Ligand not binding inside the cavity.
[2] Numbers: Pose number with correct stereochemistry of the ligand; PS: Poor stereochemistry of the ligand in all the poses.
[3] Selected: Ligand binding fully accommodated with in the cavity with appropriate stereochemistry; Not Selected: Ligand binding with partial or zero cavity occupancy and/or distorted stereochemistry

**SI Table 6G: Docking pose evaluation for LOC_Os4g02910**

| Ligand No. | Cavity Binding[1] | Pose[2] | Binding Energy (kcal/mol) | Status[3] | Ligand No. | Cavity Binding[1] | Pose[2] | Binding Energy (kcal/mol) | Status |
|---|---|---|---|---|---|---|---|---|---|
| 1 | Full | 2 | -5.2 | Selected | 15 | Partial | - | - | Discarded |
| 2 | Full | 1 | -5.3 | Selected | 16 | Full | 1 | -8.6 | Selected |
| 3 | Full | 9 | -5.1 | Selected | 17 | Full | 1 | -8.5 | Selected |
| 4 | Full | 3 | -5.4 | Selected | 18 | Full | 1 | -7.5 | Selected |
| 5 | Full | PS | - | Discarded | 19 | Full | 1 | -9.4 | Selected |
| 6 | Full | 6 | -5.6 | Selected | 20 | Full | 1 | -8.6 | Selected |
| 7 | Full | PS | - | Discarded | 21 | Full | PS | - | Discarded |
| 8 | Partial | - | - | Discarded | 22 | Full | PS | - | Discarded |
| 9 | Partial | - | - | Discarded | 23 | Full | PS | - | Discarded |
| 10 | Full | 4 | -5.2 | Selected | 24 | Full | PS | - | Discarded |
| 11 | Partial | | | Discarded | 25 | Full | - | - | Discarded |
| 12 | Full | PS | - | Discarded | 26 | Partial | - | - | Discarded |
| 13 | Partial | - | - | Discarded | 27 | Full | PS | - | Discarded |
| 14 | Partial | - | - | Discarded | 28 | Full | 1 | -6.3 | Selected |

**SI Table 6H: Docking pose evaluation for Solyc01g102720**

| Ligand No. | Cavity Binding[1] | Pose[2] | Binding Energy (kcal/mol) | Status[3] | Ligand No. | Cavity Binding[1] | Pose[2] | Binding Energy (kcal/mol) | Status |
|---|---|---|---|---|---|---|---|---|---|
| 1 | Full | 1 | -4.2 | Selected | 15 | Partial | - | - | Discarded |
| 2 | Full | 1 | -4.7 | Selected | 16 | Full | 1 | -6.3 | Selected |
| 3 | Full | 4 | -3.8 | Selected | 17 | Full | 1 | -6.5 | Selected |
| 4 | Full | 3 | -4.3 | Selected | 18 | Full | 1 | -5.8 | Selected |
| 5 | Partial | - | - | Discarded | 19 | Full | 1 | -6.8 | Selected |
| 6 | Partial | - | - | Discarded | 20 | Full | 1 | -6.6 | Selected |
| 7 | Partial | - | - | Discarded | 21 | Full | PS | - | Discarded |
| 8 | Partial | - | - | Discarded | 22 | Full | PS | - | Discarded |
| 9 | Partial | - | - | Discarded | 23 | Partial | - | - | Discarded |
| 10 | Full | 3 | -4.3 | Selected | 24 | Partial | - | - | Discarded |
| 11 | Partial | - | - | Discarded | 25 | Partial | - | - | Discarded |
| 12 | Partial | - | - | Discarded | 26 | Partial | - | - | Discarded |
| 13 | Partial | - | - | Discarded | 27 | Partial | - | - | Discarded |
| 14 | Partial | - | - | Discarded | 28 | Full | 1 | -7.2 | Selected |

[1] Full= Ligand binding completely inside the cavity; Partial: Ligand binding partially; Zero: Ligand not binding inside the cavity.
[2] Numbers: Pose number with correct stereochemistry of the ligand; PS: Poor stereochemistry of the ligand in all the poses.
[3] Selected: Ligand binding fully accommodated with in the cavity with appropriate stereochemistry; Not Selected: Ligand binding with partial or zero cavity occupancy and/or distorted stereochemistry

**SI Table 6I: Docking pose evaluation for Soly03g081320**

| Ligand No. | Cavity Binding[1] | Pose[2] | Binding Energy (kcal/mol) | Status[3] | Ligand No. | Cavity Binding[1] | Pose[2] | Binding Energy (kcal/mol) | Status |
|---|---|---|---|---|---|---|---|---|---|
| 1 | Full | 7 | -4.4 | Selected | 15 | Partial | - | - | Discarded |
| 2 | Full | 2 | -4.8 | Selected | 16 | Full | 1 | -7.2 | Selected |
| 3 | Full | 6 | -4.7 | Selected | 17 | Full | 1 | -6.9 | Selected |
| 4 | Full | 6 | -4.8 | Selected | 18 | Full | 1 | -6.5 | Selected |
| 5 | Full | 1 | -6.9 | Selected | 19 | Full | 1 | -7.7 | Selected |
| 6 | Full | 1 | -6.4 | Selected | 20 | Full | 1 | -7.4 | Selected |
| 7 | Full | PS | - | Discarded | 21 | Full | PS | - | Discarded |
| 8 | Full | PS | - | Discarded | 22 | Full | 3 | -7.8 | Selected |
| 9 | Full | PS | - | Discarded | 23 | Full | 7 | -9.2 | Selected |
| 10 | Full | 3 | -4.8 | Selected | 24 | Full | PS | - | Discarded |
| 11 | Partial | - | - | Discarded | 25 | Full | 7 | -7.3 | Selected |
| 12 | Partial | - | - | Discarded | 26 | Partial | - | - | Discarded |
| 13 | Partial | - | - | Discarded | 27 | Full | PS | - | Discarded |
| 14 | Partial | - | - | Discarded | 28 | Full | 1 | -6.0 | Selected |

[1] Full= Ligand binding completely inside the cavity; Partial: Ligand binding partially; Zero: Ligand not binding inside the cavity.
[2] Numbers: Pose number with correct stereochemistry of the ligand; PS: Poor stereochemistry of the ligand in all the poses.
[3] Selected: Ligand binding fully accommodated with in the cavity with appropriate stereochemistry; Not Selected: Ligand binding with partial or zero cavity occupancy and/or distorted stereochemistry

**SI Table 7: Analysis of ligand-induced cavity entrance geometry during 400 ns molecular dynamics simulations. (A)** Overall trajectory averages (mean ± SD) calculated over the complete 400 ns simulation. **(B–D)** Time-resolved analyses performed over consecutive 40 ns intervals for apo AT4G14500, MA-bound AT4G14500 and LPC-bound AT4G14500, respectively. The triangular entrance gate was defined by the Cα atoms of residues F214, F241, and L325. Inter-residue distances—F214–F241 (a), F214–L325 (b), and F241–L325 (c)—were measured for each frame using the Schrödinger Plot panel. Triangle area was calculated using Heron's formula: $s = (a+b+C)/2$; Area = $[(s-a)(s-b)(s-c)]^{1/2}$. Percentage area reduction relative to the apo state was calculated as: $[(Area_{apo}-Area_{ligand})/Area_{apo}]X100$, where $Area_{apo}$ and $Area_{ligand}$ are the areas of apo structure and ligand-bound structures, respectively.

**SI Table 7A: Overall trajectory statistics (400 ns)**

| System | F214–F241 distance (Å) | F214–L325 distance (Å) | F241–L325 distance (Å) | Triangle area (Å²) | Percentage Decrease in area (vs. Apo) |
|---|---|---|---|---|---|
| AT4G14500 (apo) | 15.19 ± 1.35 | 22.41 ± 2.46 | 18.00 ± 2.46 | 135.35 ± 35.42 | - |
| AT4G14500 + MA | 14.08 ± 1.15 | 17.63 ± 1.10 | 10.7 ± 1.34 | 75.13 ± 11.43 | 44.49% |
| AT4G14500 + LPC | 14.15 ± 1.06 | 19.36 ± 1.92 | 10.86 ± 2.36 | 73.58 ± 18.05 | 45.64% |

**SI Table 7B: Time-resolved cavity entrance geometry (Apo)**

| Time (ns) | F214–F241 distance (Å) | F214–L325 distance (Å) | F241–L325 distance (Å) | Triangle area (Å²) |
|---|---|---|---|---|
| 001-040 | 13.38 ± 1.00 | 17.54 ± 3.10 | 14.79 ± 3.72 | 95.03 ± 27.32 |
| 041-080 | 14.50 ± 1.07 | 21.64 ± 1.62 | 19.60 ± 1.98 | 137.43 ± 17.79 |
| 081-120 | 16.74 ± 1.22 | 23.36 ± 1.05 | 22.29 ± 1.85 | 176.29 ± 19.04 |
| 121-160 | 14.90 ± 1.17 | 23.83 ± 1.31 | 20.05 ± 2.97 | 146.37 ± 22.95 |
| 161-200 | 14.26 ± 1.52 | 22.92 ± 0.99 | 15.99 ± 2.98 | 109.73 ± 29.14 |
| 201-240 | 15.17 ± 0.89 | 23.36 ± 0.81 | 17.69 ± 2.55 | 131.91 ± 17.39 |
| 241-280 | 15.66 ± 0.78 | 23.74 ± 0.92 | 23.30 ± 3.66 | 170.24 ± 26.56 |
| 281-320 | 15.91 ± 0.68 | 23.65 ± 0.94 | 22.18 ± 4.16 | 165.79 ± 25.49 |
| 321-360 | 15.61 ± 0.68 | 23.56 ± 1.32 | 15.04 ± 1.66 | 112.68 ± 18.88 |
| 361-400 | 15.79 ± 0.60 | 20.51 ± 2.01 | 13.90 ± 1.92 | 108.46 ± 16.93 |
| **001-400** | **15.19 ± 1.35** | **22.41 ± 2.46** | **18.00 ± 2.46** | **135.35 ± 35.42** |

**SI Table 7C: Time-resolved cavity entrance geometry (AT4G14500 + MA)**

| Time (ns) | F214–F241 distance (Å) | F214–L325 distance (Å) | F241–L325 distance (Å) | Triangle area (Å²) |
|---|---|---|---|---|
| 001-040 | 13.64 ± 0.69 | 17.10 ± 1.40 | 12.06 ± 1.19 | 81.05 ± 9.75 |
| 041-080 | 13.26 ± 0.88 | 17.35 ± 1.35 | 11.44 ± 1.35 | 74.82 ± 9.81 |
| 081-120 | 12.67 ± 0.81 | 16.98 ± 0.76 | 9.67 ± 1.99 | 59.34 ± 12.97 |
| 121-160 | 13.31 ± 1.34 | 17.69 ± 0.82 | 10.55 ± 1.36 | 68.98 ± 12.46 |
| 161-200 | 14.80 ± 0.67 | 18.28 ± 1.04 | 10.58 ± 0.93 | 77.79 ± 7.87 |
| 201-240 | 14.39 ± 0.93 | 17.65 ± 1.10 | 10.45 ± 0.88 | 74.79 ± 8.01 |
| 241-280 | 14.63 ± 0.91 | 17.91 ± 0.79 | 11.00 ± 1.10 | 79.87 ± 9.12 |
| 281-320 | 14.99 ± 0.75 | 17.74 ± 0.97 | 10.73 ± 0.86 | 79.92 ± 8.14 |
| 321-360 | 14.68 ± 0.79 | 18.15 ± 0.75 | 10.72 ± 0.99 | 78.38 ± 8.65 |
| 361-400 | 14.45 ± 0.91 | 17.49 ± 1.10 | 10.64 ± 0.96 | 76.34 ± 7.97 |
| **001-400** | **14.08 ± 1.15** | **17.63 ± 1.10** | **10.7 ± 1.34** | **75.13 ± 11.43** |

**SI Table 7D: Time-resolved cavity entrance geometry (AT4G14500 + LPC)**

| Time (ns) | F214–F241 distance (Å) | F214–L325 distance (Å) | F241–L325 distance (Å) | Triangle area (Å²) |
|---|---|---|---|---|
| "01-40" | 14.14 ± 0.84 | 18.89 ± 1.88 | 11.06 ± 2.47 | 76.09 ± 18.09 |
| 41-80 | 14.03 ± 0.78 | 19.35 ± 1.95 | 9.87 ± 1.18 | 63.96 ± 13.00 |
| 81-120 | 14.84 ± 0.57 | 21.23 ± 1.16 | 11.70 ± 1.27 | 82.60 ± 11.42 |
| 121-160 | 14.45 ± 0.66 | 21.15 ± 0.93 | 11.33 ± 1.24 | 76.47 ± 12.47 |
| 161-200 | 14.49 ± 0.71 | 19.36 ± 1.98 | 11.14 ± 1.74 | 77.98 ± 15.09 |
| 201-240 | 14.60 ± 0.62 | 18.37 ± 1.50 | 10.75 ± 1.50 | 77.73 ± 11.84 |
| 241-280 | 14.56 ± 0.56 | 18.76 ± 1.72 | 10.65 ± 1.07 | 76.36 ± 8.55 |
| 281-320 | 14.63 ± 0.56 | 17.94 ± 1.52 | 7.84 ± 1.69 | 54.53 ± 13.59 |
| 321-360 | 13.41 ± 1.43 | 19.52 ± 2.06 | 10.84 ± 3.35 | 68.45 ± 25.82 |
| 361-400 | 12.44 ± 0.96 | 19.08 ± 1.31 | 13.53 ± 2.58 | 81.67 ± 22.78 |
| **001-400** | **14.15 ± 1.06** | **19.36 ± 1.92** | **10.86 ± 2.36** | **73.58 ± 18.05** |

**Supplementary Material and Methods**

**Detailed Protein Expression and Purification**

Transformed *E. coli* cells were cultured in 2 L LB medium (HiMedia) and harvested by centrifugation at 7,800 × g for 15 min at 4 °C. Cell pellets were resuspended in lysis buffer containing 20 mM Tris-HCl (pH 8.5), 10% glycerol, 5 mM β-mercaptoethanol, and 1 M NaCl, supplemented with lysozyme (0.5 mg $mL^{-1}$) and PMSF (5 mM). Cells were sonicated for 40 min at 4 °C, and lysates were clarified by centrifugation at 34,600 × g for 40 min at 4 °C. The clarified supernatants were loaded onto HisTrap HP columns (Cytiva), washed with Buffer A containing 20 mM Tris-HCl (pH 8.5), 5% glycerol, 1 M NaCl, and 5 mM β-mercaptoethanol, and proteins were eluted with 500 mM imidazole. Expression optimization conditions for the individual proteins are provided in SI Table 3.

For AT4G14500, the His-tag was removed by SUMO protease digestion. The protein was subsequently purified using a second HisTrap HP column, followed by size-exclusion chromatography using a Sephacryl 16/60 S-100 column equilibrated with 20 mM Tris-HCl (pH 8.5), 5% glycerol, 500 mM NaCl, and 5 mM β-mercaptoethanol. Purified proteins were verified by 15% SDS-PAGE and concentrated using Amicon® Ultra centrifugal filters with a 10 kDa molecular weight cutoff.

**Oligomeric-State Determination**

The oligomeric state of AT4G14500 was determined by size-exclusion chromatography. Bio-Rad molecular weight standards (vitamin B12, myoglobin, ovalbumin, γ-globulin, and thyroglobulin) were used to calibrate the Sephacryl 16/60 S-100 column. The partition coefficient (Kav) was calculated as (Ve - Vo)/(Vt - Vo) where Ve, Vo and Vt are elution volume, void volume and total volume, respectively. Thyroglobulin (670 kDa) and γ-globulin (158 kDa) were used to determine the void volume.

**Lipid Handling and Preparation**

For the protein-lipid overlay assay, myristic acid (MA), decanoic acid, and dodecanoic acid were dissolved in methanol to a concentration of 50 mM and subsequently diluted with water to 12.5 mM. Lysophosphatidylcholine (LPC), which comprises a mixture of molecular species, was dissolved in water to a nominal concentration of 20 mM. For the purpose of concentration calculation, the LPC concentration was expressed using $C_{10}H_{22}NO_7P$ as the reference species. The LPC stock solution was subsequently diluted with water to a nominal concentration of 12.5 mM. Sphingomyelin was dissolved in a water:methanol mixture (1:1:1, v/v/v) to a concentration of 24.62 mM and subsequently diluted with water to 12.5 mM. 2-Aminooctanoic acid was dissolved in an acetic acid mixture (1:1, v/v) to a concentration of 25 mM and subsequently diluted with water to 12.5 mM. N-Acetyl-L-methionine was dissolved in methanol to a concentration of 50 mM and subsequently diluted with water to 12.5 mM. For the protein–lipid overlay assay, 2 µL of each lipid solution at 12.5 mM was spotted onto the assay strip. For the internal fluorescence assay, the same lipid stock solutions described above were used, with subsequent dilutions prepared in 20 mM Tris-HCl containing 150 mM NaCl. For biolayer interferometry (BLI), the LPC stock solution described above was used, and subsequent dilutions were prepared in 20 mM sodium phosphate buffer (pH 7.5) containing 150 mM NaCl.

**Protein Modeling and Ligand Preparation**

Tertiary structures of all nine minimal START proteins were predicted using AlphaFold2 (4) with default settings. Per-residue local distance difference test (pLDDT) scores were calculated for the core START domain (SI Table 1). Cavity volumes were analyzed using the CASTp server with default parameters (55). Lipids from diverse families were selected based on the LIPID MAPS classification system. Consistent with the START-domain cavity size, small- to medium-sized lipids were included, whereas large lipids such as waxes were excluded. Plant-specific lipids were prioritized, and their biological relevance was assessed through literature review before final selection (SI Table 4). Plant hormones were also included to assess potential protein interactions. Ligand structures were drawn using ChemSketch 2021.1.2 (Advanced Chemistry Development, Inc. [ACD/Labs], Toronto, ON, Canada).

**Molecular Docking**

Molecular docking of minimal START domains with the selected ligands was performed using PyRx v0.8 (56), which integrates AutoDock Vina. Ligand structures in SDF format were obtained from PubChem or LIPID MAPS, and AlphaFold2-predicted structures were used as receptors. Protein structures were prepared by adding polar hydrogens and converting them to PDBQT format using AutoDock Tools. Ligands were energy-minimized and converted to PDBQT format using Open Babel within PyRx. Blind docking was performed with an exhaustiveness value of 8 using a grid encompassing the entire protein surface (SI Table 5). Multiple binding poses were generated and ranked by predicted binding affinity. Binding poses were further analyzed using PyMOL v2.3.2 (The PyMOL Molecular Graphics System, Version 3.0, Schrödinger, LLC).

**Confocal Microscopy**

The AT4G14500 coding sequence was amplified from *Arabidopsis thaliana* leaf cDNA using gene-specific primers (forward primer containing a 5′ CACC overhang) and cloned into the pENTR/D-TOPO vector (Invitrogen). The gene was recombined into the Gateway destination vector pK7WGF2 to generate an N-terminal GFP-tagged AT4G14500 fusion. Transient expression in *Nicotiana benthamiana* leaves was performed using *Agrobacterium tumefaciens* strain GV3101. Agrobacterium cultures were resuspended in infiltration buffer (10 mM MES, pH 5.7, 10 mM $MgCl_2$, 100 µM acetosyringone). For subcellular localization, GFP-tagged AT4G14500 ($OD_{600}$ = 0.4) was co-expressed with fluorescent organelle markers ($OD_{600}$ = 0.4) and the P19 silencing suppressor ($OD_{600}$ = 0.2). The organelle marker constructs ER-rk, G-rk, px-rk, mt-rk, pt-rk, and pm-rk, expressing mCherry-tagged markers for the endoplasmic reticulum, Golgi apparatus, peroxisomes, mitochondria, plastids, and plasma membrane, respectively, were used as previously described (57). H2B-RFP was used as the nuclear marker (58). Plants were grown at 22 °C under a 16 h light/8 h dark photoperiod, and fluorescence was examined 48 h after infiltration using a Leica TCS SP8 laser-scanning confocal microscope. GFP fluorescence was excited using a 488 nm argon laser and detected between 500–550 nm, whereas mCherry/RFP fluorescence was excited using a 561 nm DPSS laser and detected between 580–630 nm. Images were acquired using a 53.1 µm pinhole (999.82 mAiry units), and identical acquisition settings were maintained for all samples within each experiment. Co-localization between GFP-tagged AT4G14500 and the corresponding fluorescent organelle markers was quantified using the Coloc 2 plugin in Fiji by calculating Pearson's correlation coefficient (PCC). Regions of interest corresponding to individual transformed cells were analyzed from 3–6 independent biological replicates for each organelle marker under identical imaging conditions. Pearson's correlation coefficients are presented as mean ± SD.

**Reagents and materials used in this study**

| Reagents/Materials | Company Name | Catalogue No. | Purity (%) |
|---|---|---|---|
| Sodium Myristate | Tokyo Chemical Industry (TCI) | M0483 | >98.0 |
| Dodecanoic Acid | Sigma-Aldrich | L556 | >98.0 |
| Decanoic Acid | Sigma-Aldrich | C1875 | >98.0 |
| N-acetyl L-methionine | Sigma-Aldrich | 01310 | >98.5 |
| DL-2-aminooctanoic acid | Sigma-Aldrich | 217700 | >99.0 |
| Sphingomyelin | Sigma-Aldrich | 85615 | >98.0 |
| Lysophosphotidylcholine | Sigma-Aldrich | PHR2631 | |
| LPC strips | Echelon Biosciences | customized | |
| Indole-3-acetic acid | Sigma-Aldrich | I 3750 | >98.0 |
| Gibbrelic acid | Sigma-Aldrich | G7645 | >90.0 |
| Abscisic Acid | Sigma-Aldrich | A4906 | >98.0 |
| Kinetin | Sigma-Aldrich | K3378 | >98.0 |
| Jasmonic Acid | Sigma-Aldrich | J2500 | >97.0 |
| Salicylic Acid | Sigma-Aldrich | S5922 | >99.0 |
| 14-hydroxymyristic acid | Med Chem Express | HY-W012017 | >99.0 |
| Miltefosin | Sigma-Aldrich | M5571 | >98.0 |
| Edelfosine | Sigma-Aldrich | SML0332 | >95.0 |
| DPN1 | New England Biolabs | R0176 | |
| BamH1 | New England Biolabs | R0136 | |
| Xho1 | New England Biolabs | R0146 | |
| Tris, Free base | Himedia | MB029 | >99.8 |
| 2-Mercaptoethanol | Sisco Research Laboratories (SRL) | 83759 | >99.0 |
| Glycerol | Qualigen | Q15457 | |
| NaCl | Qualigen | Q27608 | >99.5 |
| Imidazole | Sisco Research Laboratories (SRL) | 61510 | >99.5 |
| Gel Filteration Marker | BioRad | 1511901 | |
| Tween 20 | Sigma-Aldrich | P2287 | |
| Skimmed milk | Sigma-Aldrich | 70166 | |
| BioRad lumiol kit | BioRad | 1705060 | |
| Anti His antibody | Santa cruz | sc-8036 | |
| HRP antibody | Santa cruz | sc-525408 | |
| MES | Sigma-Aldrich | M8250 | >99.5 |
| $MgCl_2$ | Sigma-Aldrich | M8266 | >98.0 |
| Acetosyringone | Sigma-Aldrich | D134406 | >97.0 |
| EZ-Link™ Sulfo-NHS-LC-Biotin/Biotin | Thermo Scientific | A39257 | |
| Biocytin | Sigma-Aldrich | B4261 | >98.0 |
| SSA-sensor | Forte Bio | 18-5057 | |

**References**


1. Madeira, F., Madhusoodanan, N., Lee, J., Eusebi, A., Niewielska, A., Tivey, A.R.N., Lopez, R. and Butcher, S. (2024) The EMBL-EBI Job Dispatcher sequence analysis tools framework in 2024. *Nucleic Acids Res*, **52**, W521-W525.
2. Schneider, T.D. and Stephens, R.M. (1990) Sequence logos: a new way to display consensus sequences. *Nucleic Acids Res*, **18**, 6097-6100.
3. Crooks, G.E., Hon, G., Chandonia, J.M. and Brenner, S.E. (2004) WebLogo: a sequence logo generator. *Genome Res*, **14**, 1188-1190.
4. Jumper, J., Evans, R., Pritzel, A., Green, T., Figurnov, M., Ronneberger, O., Tunyasuvunakool, K., Bates, R., Zidek, A., Potapenko, A. *et al.* (2021) Highly accurate protein structure prediction with AlphaFold. *Nature*, **596**, 583-589.
5. Lopez-Velazquez, J.G., Ayon-Reyna, L.E., Vega-Garcia, M.O., Lopez-Angulo, G., Lopez-Lopez, M.E., Lopez-Zazueta, B.A. and Delgado-Vargas, F. (2022) Caprylic acid in Vitex mollis fruit and its inhibitory activity against a thiabendazole-resistant Colletotrichum gloeosporioides strain. *Pest Manag Sci*, **78**, 5271-5280.
6. Cvak, L., Jegorov, A., Sedmera, P., Cisarova, I., Cejka, J., Kratochvil, B. and Pakhomova, S. (2005) Norleucine, a natural occurrence in a novel ergot alkaloid gamma-ergokryptinine. *Amino Acids*, **29**, 145-150.
7. Schrick, K., Bruno, M., Khosla, A., Cox, P.N., Marlatt, S.A., Roque, R.A., Nguyen, H.C., He, C., Snyder, M.P., Singh, D. *et al.* (2014) Shared functions of plant and mammalian StAR-related lipid transfer (START) domains in modulating transcription factor activity. *BMC Biol*, **12**, 70.
8. Graham, S.A. (1989) Cuphea: a new plant source of medium-chain fatty acids. *Crit Rev Food Sci Nutr*, **28**, 139-173.
9. James, A.T. (1963) The biosynthesis of long-chain saturated and unsaturated fatty acids in isolated plant leaves. *Biochim Biophys Acta*, **70**, 9-19.
10. Zhang, B., Du, H., Yang, S., Wu, X., Liu, W., Guo, J., Xiao, Y. and Peng, F. (2023) Physiological and Transcriptomic Analyses of the Effects of Exogenous Lauric Acid on Drought Resistance in Peach (Prunus persica (L.) Batsch). *Plants (Basel)*, **12**.
11. Kumaraswamy, S., Yogendra, K., Sotelo-Cardona, P., Shivanna, A., Hemalatha, S., Mohan, M. and Srinivasan, R. (2025) Non-targeted metabolomics reveals fatty acid and associated pathways driving resistance to whitefly and tomato leafminer in wild tomato accessions. *Sci Rep*, **15**, 3754.
12. Mallick, M. and Mondal, H.A. (2023) Vascular dodecanoic acid of Arabidopsis mediates an insect resistance against Myzus persicae. *Arch Insect Biochem Physiol*, **112**, e21986.
13. Kurima, K., Jimbo, H., Fujihara, T., Saito, M., Ishikawa, T. and Wada, H. (2024) High Myristic Acid in Glycerolipids Enhances the Repair of Photodamaged Photosystem II under Strong Light. *Plant Cell Physiol*, **65**, 790-797.
14. Meinnel, T., Dian, C. and Giglione, C. (2020) Myristoylation, an Ancient Protein Modification Mirroring Eukaryogenesis and Evolution. *Trends Biochem Sci*, **45**, 619-632.
15. Hu, Z., Shi, J., Feng, S., Wu, X., Shao, S. and Shi, K. (2023) Plant N-acylethanolamines play a crucial role in defense and its variation in response to elevated CO(2) and temperature in tomato. *Hortic Res*, **10**, uhac242.
16. Keereetaweep, J., Blancaflor, E.B., Hornung, E., Feussner, I. and Chapman, K.D. (2015) Lipoxygenase-derived 9-hydro(pero)xides of linoleoylethanolamide interact with ABA signaling to arrest root development during Arabidopsis seedling establishment. *Plant J*, **82**, 315-327.
17. Keereetaweep, J., Blancaflor, E.B., Hornung, E., Feussner, I. and Chapman, K.D. (2013) Ethanolamide oxylipins of linolenic acid can negatively regulate Arabidopsis seedling development. *Plant Cell*, **25**, 3824-3840.

18. Dong, W., Lv, H., Xia, G. and Wang, M. (2012) Does diacylglycerol serve as a signaling molecule in plants? *Plant Signal Behav*, **7**, 472-475.
19. Yang, Y. and Benning, C. (2018) Functions of triacylglycerols during plant development and stress. *Curr Opin Biotechnol*, **49**, 191-198.
20. Aronsson, H., Schottler, M.A., Kelly, A.A., Sundqvist, C., Dormann, P., Karim, S. and Jarvis, P. (2008) Monogalactosyldiacylglycerol deficiency in Arabidopsis affects pigment composition in the prolamellar body and impairs thylakoid membrane energization and photoprotection in leaves. *Plant Physiol*, **148**, 580-592.
21. Wi, S.J., Seo, S., Cho, K., Nam, M.H. and Park, K.Y. (2014) Lysophosphatidylcholine enhances susceptibility in signaling pathway against pathogen infection through biphasic production of reactive oxygen species and ethylene in tobacco plants. *Phytochemistry*, **104**, 48-59.
22. Yamaoka, Y., Shin, S., Lee, Y., Ito, M., Lee, Y. and Nishida, I. (2021) Phosphatidylserine Is Required for the Normal Progression of Cell Plate Formation in Arabidopsis Root Meristems. *Plant Cell Physiol*, **62**, 1396-1408.
23. Van Der Rest, B., Rolland, N., Boisson, A.M., Ferro, M., Bligny, R. and Douce, R. (2004) Identification and characterization of plant glycerophosphodiester phosphodiesterase. *Biochem J*, **379**, 601-607.
24. Umemura, K., Ogawa, N., Yamauchi, T., Iwata, M., Shimura, M. and Koga, J. (2000) Cerebroside elicitors found in diverse phytopathogens activate defense responses in rice plants. *Plant Cell Physiol*, **41**, 676-683.
25. Charruyer, A., Bell, S.M., Kawano, M., Douangpanya, S., Yen, T.Y., Macher, B.A., Kumagai, K., Hanada, K., Holleran, W.M. and Uchida, Y. (2008) Decreased ceramide transport protein (CERT) function alters sphingomyelin production following UVB irradiation. *J Biol Chem*, **283**, 16682-16692.
26. Hanada, K. (2014) Co-evolution of sphingomyelin and the ceramide transport protein CERT. *Biochim Biophys Acta*, **1841**, 704-719.
27. Liang, H., Yao, N., Song, J.T., Luo, S., Lu, H. and Greenberg, J.T. (2003) Ceramides modulate programmed cell death in plants. *Genes Dev*, **17**, 2636-2641.
28. Sun, M., Li, L., Wang, C., Wang, L., Lu, D., Shen, D., Wang, J., Jiang, C., Cheng, L., Pan, X. *et al.* (2022) Naringenin confers defence against Phytophthora nicotianae through antimicrobial activity and induction of pathogen resistance in tobacco. *Mol Plant Pathol*, **23**, 1737-1750.
29. Zhang, Z., Chen, B., Zhang, Y., Mo, X., Lu, X., Wang, T., Liang, C. and Tian, J. (2026) Integrated transcriptomic and proteomic analysis reveals inhibitory effects of naringenin application on soybean roots mainly by interfering with auxin and ROS concentrations. *BMC Plant Biol*, **26**, 247.
30. An, J., Kim, S.H., Bahk, S., Vuong, U.T., Nguyen, N.T., Do, H.L., Kim, S.H. and Chung, W.S. (2021) Naringenin Induces Pathogen Resistance Against Pseudomonas syringae Through the Activation of NPR1 in Arabidopsis. *Front Plant Sci*, **12**, 672552.
31. Shen, X., Liu, Y., Luo, X. and Yang, Z. (2019) Advances in Biosynthesis, Pharmacology, and Pharmacokinetics of Pinocembrin, a Promising Natural Small-Molecule Drug. *Molecules*, **24**.
32. Abo-Kadoum, M.A., Abouelela, M.E., Al Mousa, A.A., Abo-Dahab, N.F., Mosa, M.A., Helmy, Y.A. and Hassane, A.M.A. (2022) Resveratrol biosynthesis, optimization, induction, bio-transformation and bio-degradation in mycoendophytes. *Front Microbiol*, **13**, 1010332.
33. Baumert, A., Kuzovkina, I.N., Krauss, G., Hieke, M. and Groger, D. (1982) Biosynthesis of rutacridone in tissue cultures of Ruta graveolens L. *Plant Cell Rep*, **1**, 168-171.
34. Sultan, O.S., Kantilal, H.K., Khoo, S.P., Davamani, A.F., Eusufzai, S.Z., Rashid, F., Jamayet, N.B., Soh, J.A., Tan, Y.Y. and Alam, M.K. (2022) The Potential of alpha-Mangostin from Garcinia mangostana as an Effective Antimicrobial Agent-A Systematic Review and Meta-Analysis. *Antibiotics (Basel)*, **11**.
35. Du, Y., Fu, X., Chu, Y., Wu, P., Liu, Y., Ma, L., Tian, H. and Zhu, B. (2022) Biosynthesis and the Roles of Plant Sterols in Development and Stress Responses. *Int J Mol Sci*, **23**.

36. Ohyama, K., Suzuki, M., Kikuchi, J., Saito, K. and Muranaka, T. (2009) Dual biosynthetic pathways to phytosterol via cycloartenol and lanosterol in Arabidopsis. *Proc Natl Acad Sci U S A*, **106**, 725-730.
37. Ma, A., Diao, H., Xia, T., Sun, J., Feng, L., Stephenson, M., Osbourn, A., Wu, R. and Qi, X. (2026) Atomic basis for functional evolution of plant lanosterol synthase. *New Phytol*, **249**, 373-388.
38. Tanveer, M., Shahzad, B., Sharma, A., Biju, S. and Bhardwaj, R. (2018) 24-Epibrassinolide; an active brassinolide and its role in salt stress tolerance in plants: A review. *Plant Physiol Biochem*, **130**, 69-79.
39. Shahzad, B., Tanveer, M., Che, Z., Rehman, A., Cheema, S.A., Sharma, A., Song, H., Rehman, S.U. and Zhaorong, D. (2018) Role of 24-epibrassinolide (EBL) in mediating heavy metal and pesticide induced oxidative stress in plants: A review. *Ecotoxicol Environ Saf*, **147**, 935-944.
40. Yu, B., Wang, L., Guan, Q., Xue, X., Gao, W. and Nie, P. (2023) Exogenous 24-epibrassinolide promoted growth and nitrogen absorption and assimilation efficiency of apple seedlings under salt stress. *Front Plant Sci*, **14**, 1178085.
41. Ahsan, H., Ahad, A. and Siddiqui, W.A. (2015) A review of characterization of tocotrienols from plant oils and foods. *J Chem Biol*, **8**, 45-59.
42. Almagro, L., Correa-Sabater, J.M., Sabater-Jara, A.B. and Pedreno, M.A. (2022) Biotechnological production of beta-carotene using plant in vitro cultures. *Planta*, **256**, 41.
43. Munne-Bosch, S. (2007) Alpha-tocopherol: a multifaceted molecule in plants. *Vitam Horm*, **76**, 375-392.
44. Zhao, Y. (2012) Auxin biosynthesis: a simple two-step pathway converts tryptophan to indole-3-acetic acid in plants. *Mol Plant*, **5**, 334-338.
45. Tian, H., Xu, Y., Liu, S., Jin, D., Zhang, J., Duan, L. and Tan, W. (2017) Synthesis of Gibberellic Acid Derivatives and Their Effects on Plant Growth. *Molecules*, **22**.
46. Mo, W., Zheng, X., Shi, Q., Zhao, X., Chen, X., Yang, Z. and Zuo, Z. (2024) Unveiling the crucial roles of abscisic acid in plant physiology: implications for enhancing stress tolerance and productivity. *Front Plant Sci*, **15**, 1437184.
47. Barciszewski, J., Massino, F. and Clark, B.F. (2007) Kinetin--a multiactive molecule. *Int J Biol Macromol*, **40**, 182-192.
48. Koo, A.J. and Howe, G.A. (2009) The wound hormone jasmonate. *Phytochemistry*, **70**, 1571-1580.
49. Koo, Y.M., Heo, A.Y. and Choi, H.W. (2020) Salicylic Acid as a Safe Plant Protector and Growth Regulator. *Plant Pathol J*, **36**, 1-10.
50. Croft, S.L. and Engel, J. (2006) Miltefosine--discovery of the antileishmanial activity of phospholipid derivatives. *Trans R Soc Trop Med Hyg*, **100 Suppl 1**, S4-8.
51. Vogler, W.R., Liu, J., Volpert, O., Ades, E.W. and Bouck, N. (1998) The anticancer drug edelfosine is a potent inhibitor of neovascularization in vivo. *Cancer Invest*, **16**, 549-553.
52. Liu, C., Liu, F., Cai, J., Xie, W., Long, T.E., Turner, S.R., Lyons, A. and Gross, R.A. (2011) Polymers from fatty acids: poly(omega-hydroxyl tetradecanoic acid) synthesis and physico-mechanical studies. *Biomacromolecules*, **12**, 3291-3298.
53. Dubey, P., Sharma, P. and Kumar, V. (2017) Structural profiling of wax biopolymer from Pinus roxburghii Sarg. needles using spectroscopic methods. *Int J Biol Macromol*, **104**, 261-273.
54. Jayaram, B., Singh, T., Mukherjee, G., Mathur, A., Shekhar, S. and Shekhar, V. (2012) Sanjeevini: a freely accessible web-server for target directed lead molecule discovery. *BMC Bioinformatics*, **13 Suppl 17**, S7.
55. Tian, W., Chen, C., Lei, X., Zhao, J. and Liang, J. (2018) CASTp 3.0: computed atlas of surface topography of proteins. *Nucleic Acids Res*, **46**, W363-W367.
56. Dallakyan, S. and Olson, A.J. (2015) Small-molecule library screening by docking with PyRx. *Methods Mol Biol*, **1263**, 243-250.

57. Nelson, B.K., Cai, X. and Nebenfuhr, A. (2007) A multicolored set of in vivo organelle markers for co-localization studies in Arabidopsis and other plants. *Plant J*, **51**, 1126-1136.
58. Martin, K., Kopperud, K., Chakrabarty, R., Banerjee, R., Brooks, R. and Goodin, M.M. (2009) Transient expression in Nicotiana benthamiana fluorescent marker lines provides enhanced definition of protein localization, movement and interactions in planta. *Plant J*, **59**, 150-162.
